# Scalable dynamic characterization of synthetic gene circuits

**DOI:** 10.1101/635672

**Authors:** Neil Dalchau, Paul K. Grant, Prashant Vaidyanathan, Carlo Spaccasassi, Colin Gravill, Andrew Phillips

## Abstract

The dynamic behavior of synthetic gene circuits plays a key role in ensuring their correct function. Although there has been substantial work on modeling dynamic behavior after circuit construction, the forward engineering of dynamic behavior remains a major challenge. Previous engineering methods have focused on quantifying average behaviors of circuits over an extended time window, however this provides a static characterization of behavior that is a poor predictor of dynamics. Here we present a method for characterizing the dynamic behavior of synthetic gene circuits, using parameter inference of dynamical system models applied to time-series measurements of cell cultures growing in microtiter plates. We demonstrate that the behaviors of simple devices can be characterized dynamically and used to predict the behaviors of more complex circuits. Specifically, we compose 23 biological parts into 9 devices and use them to design 9 synthetic gene circuits in E. coli that provide core functionality for engineering cell behavior at the population level, including relays, receivers and a degrader. We embody our method in a software package and corresponding programming language. Our method supports the notion of an inference graph for iterative inference of models as new circuits are constructed, without the need to infer all models from scratch, and lays the foundation for characterizing large libraries of synthetic gene circuits in a scalable manner.

## Introduction

As increasingly complex gene circuits are designed and implemented in living cells (1), the precise dynamic behavior of these circuits is becoming increasingly important for ensuring their correct function. For instance, genetically engineered microbes must produce specific quantities of proteins at particular times to optimize the yield of compounds such as medicines (2), materials (3) and chemicals (4), and genetically engineered immune therapies must express the correct levels of proteins at the appropriate time to avoid over-stimulating the immune system and potentially killing the patient (5).

Although there has been substantial work on modeling dynamic behavior after circuit construction, the forward engineering of dynamic behavior remains a major challenge. Specifically, our ability to accurately predict the dynamic behavior of gene circuits is limited by our quantitative understanding of their properties. This understanding can be represented as a mathematical model, which can be used to optimize circuit performance and inform the design of future circuits (6, 7). Recently, mathematical modelling was also used to facilitate the design of control circuits, which can be used to ensure that a nominal circuit can continue to function in spite of perturbation (8). For models to accurately predict dynamic behaviour, their kinetic parameters, which include binding affinities, rates of transcription and translation, and rates of enzyme-catalyzed reactions, must be specified. Obtaining suitable model parameters presents a challenge, since these parameters often cannot be measured directly or derived from first principles, and must instead be inferred from observations of the dynamic behavior of circuit components over time, such as the concentrations of RNA and proteins. Furthermore, many of these quantities are not directly observable inside a living cell, requiring alternative observations such as fluorescence and absorbance over time to be used instead.

Previous engineering methods have used fluorescent reporters to obtain a static measurement of gene expression for a given a promoter (9–11). However, these methods focus on quantifying average behaviors over an extended time window, resulting in a static characterization that is a poor predictor of dynamics. More recent work has shown how to reconstruct the dynamics of promoter activity upstream of a measurable protein, using splines and differentiation of a model with assumed parameter values (12). A large number of more general parameter inference strategies for dynamical systems also exist in the literature, for deterministic (13) and stochastic models (14). Others exploit linear sub-structures in model equations to perform more efficient inference (15).

This paper presents a method for characterizing the dynamic behavior of synthetic gene circuits. The method relies on parameter inference of dynamical system models, applied to time-series measurements of cell cultures growing in microtiter plates. By comparing fluorescent protein measurements over time to the dynamic behavior of a mathematical model, we infer the values of model parameters that cannot be measured directly. We demonstrate that the behaviors of simple devices can be characterized dynamically and used to predict the behaviors of more complex circuits. Specifically, we compose 23 biological parts into 9 devices and use them to design 9 synthetic gene circuits in E. coli that provide core functionality for engineering cell behavior at the population level. The devices include Receivers capable of sensing and responding to two homoserine lactone (HSL) signaling molecules simultaneously, Relays capable of receiving one signal and producing the other signal in response, and a Degrader that degrades both signals. Our method allows computational models to be associated with gene circuits in a modular fashion, by consistently reusing models of shared genetic devices that constitute these circuits. We embody our method in a software package and corresponding programming language. Our method supports the notion of an inference graph for iterative inference of models as new circuits are constructed, without the need to infer all models from scratch. We demonstrate this by iteratively constructing a repository of models and re-using parameter values of existing devices as new devices are introduced. Propagating parameter values in this way removes the need to rerun the inference problem for an entire repository of devices each time a new device is added. Our work lays the foundation for a scalable approach to characterizing large libraries of synthetic gene circuits, by iteratively building on existing knowledge.

## Results

### Dynamic characterization method

We first present our dynamic characterization method, using the terminology of Fig. 1A. We define a genetic *device* as a sequence of genetic *parts*, which can include promoters, ribosome binding sites, protein coding regions and terminators. Devices are assembled into *circuits*, which are transformed into cell *cultures*. The dynamic behavior of each culture is measured under different *treatments*, such as different concentrations of input signals. Measurements of the culture can include culture density and the intensity of spectrally distinct fluorescent proteins. The method uses culture-level measurements in combination with dynamical systems models of intra-cellular concentrations to infer quantitative properties of parts and devices. In doing so, it enables model predictions of intracellular dynamics by reconstructing bulk culture measurements (Fig. 1B).

**Fig. 1.**
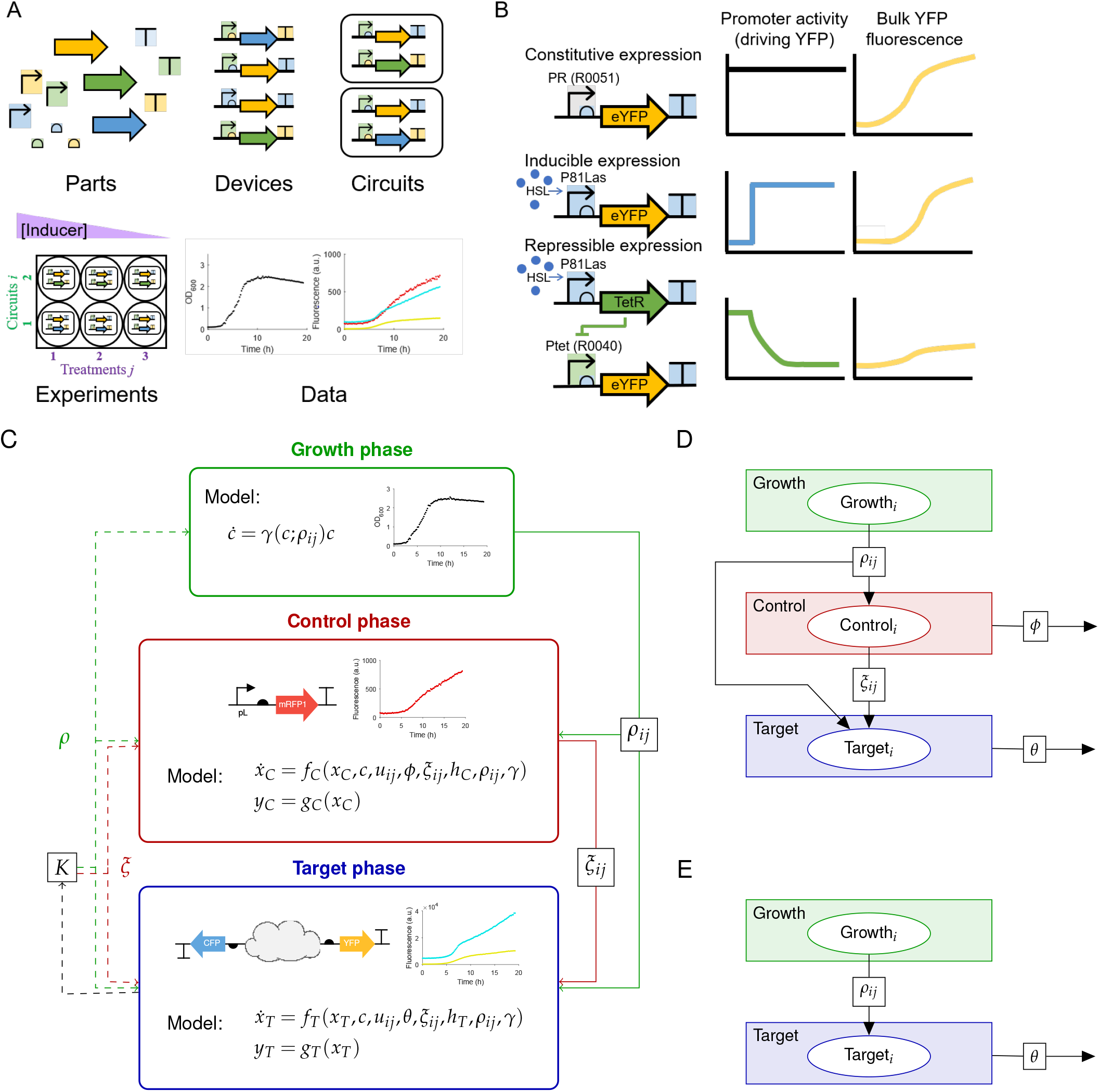
Understanding the properties of genetic circuits via dynamic bulk fluorescence measurements. **A.** A collection of genetic parts can be composed combinatorially into a set of devices that can be introduced singly or in combination into a cellular context to produce genetic circuits. The circuits (indexed by *i*) can be exposed to a variety of treatment conditions (indexed by *j*), in a set of experiments that produce data such as optical density (OD_600_) and bulk fluorescence at three different wavelengths corresponding to mRFP1, eCFP and eYFP. This hierarchical organization allows for the encoding of hypotheses about modularity and the reuse of measured properties in new contexts to allow for rational forward engineering. **B.** Dynamic bulk fluorescence is a rich data source but its relationship to the underlying promoter activity is non-trivial. Constitutive expression by constant promoter activity leads to a sigmoidal bulk fluorescence curve due to sigmoidal growth of cell density. Inducible expression leads to a (near-)step change in promoter activity, but is almost indistinguishable from constitutive expression. An inverter system drastically alters bulk fluorescence dynamics, which presents a challenge for time-window selection of static characterization. **C.** Our method seeks to infer the quantitative properties of a synthetic circuit while accounting for the feedback of the circuit consuming cellular resources and inhibiting growth. To approximate this *closed loop* system, an *open loop* approximation is proposed that allows each cell culture, expressing a given circuit *i* under a given treatment *j*, to be described by models of cell growth (*γ*) and gene expression capacity (*h*_*C*_, *h*_*T*_) that are parametrized by culture-specific parameters *u*_*ij*_, *ρ*_*ij*_ and *ξ*_*ij*_. The models of the control signal (*f*_*C*_) and the target circuit (*f*_*T*_) therefore embed these circuit-specific factors. **D.** Inference graph representing the three stages of ratiometric dynamic characterization for a single circuit *i*. The Growth node infers the growth parameters *ρ*_*ij*_ for the circuit *i* under a range of treatments *j*. This is achieved by fitting a growth model to corresponding growth data, such as optical density of the culture, for each treatment. The Control node uses the growth parameters *ρ*_*ij*_ to infer the control parameters *ξ*_*ij*_ for the same circuit *i* under the same treatments *j*. This is achieved by fitting a control model to corresponding control data, such as the concentration of fluorescent protein expressed by a constitutive promoter, for each treatment. Parameters *ϕ* shared between control models, such as the degradation of the constitutively expressed fluorescent protein, are also inferred. Finally, the Target node uses the growth and control parameters to infer parameters *θ* shared between target models, such as the strength of promoters used in the circuits. **E.** Inference graph representing the two stages of direct dynamic characterization. The graph is similar to (E) except that the Control node is omitted.

Inference of model parameters is complicated by the feedback that exists between a synthetic gene circuit and the host cell in which it operates (Fig. 1C). The circuit places a burden on the host cell resources, which in turn affects circuit behavior (16–21). In order to reduce the bias that can result from inaccurately representing the burden on host cell resources (*K* in Fig. 1C), our method does not model these resources explicitly, as attempted in (22). Instead, we model the circuit in *open loop*, by allowing parameters quantifying cell growth (*ρ*) and gene expression (*ξ*) to take on *culture-specific* values. Culture-specific cell growth incorporates resource burden indirectly, since we observe slower growth as the synthetic circuit increases its burden on the host cell. Culture-specific gene expression captures differences in transcription, translation and degradation that arise from differences in culture conditions. This might include an increased burden on the ribosome pool leading to lower translation (21, 23), increased burden on polymerases leading to lower transcription (24), or differences in the degradation of different mRNA sequences (25). Accordingly, these can include both intrinsic and extrinsic properties (11) of synthetic gene circuits. By modelling cell growth and gene expression with culture-specific values, we can arbitrarily account for the interdependence between the two (26), despite not explicitly modeling host cell resources.

In order to estimate culture-specific gene expression parameters, we developed a dynamic version of *ratiometric characterization* (11, 27). The ratiometric method seeks to separate intrinsic properties of a circuit, such as the ability of a promoter to recruit polymerase, from extrinsic properties such as cellular metabolic burden and ribosome availability. This can be achieved by using a *standard promoter* to drive expression of a spectrally distinct fluorescent protein, which is assumed to capture the same extrinsic factors that influence fluorescent protein expression driven by other promoters in the circuit.

Dynamic characterization relies on three distinct phases (Fig. 1C), where each culture is indexed by the circuit *i* that it contains and the treatment *j* under which it is measured:

1. A *growth phase* characterizes the dynamics of culture growth, by comparing simulations of a *growth model* with measurements of culture density over time. The growth model is of the form

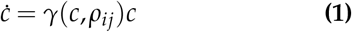

where *c*(*t*) is the culture density, *ρ*_*ij*_ are the culture-specific growth parameters and the function *γ*(*c*(*t*), *ρ*_*ij*_) describes the specific growth rate of the culture.
2. A *control phase* characterizes the gene expression of the culture, by comparing simulations of a *control model* with the expression of a control signal over time, such as the intensity of a fluorescent protein driven by a constitutive promoter, or the expression of a housekeeping gene. Although the culture may be genetically engineered to produce the control signal, we consider this to form part of the culture itself and to be distinct from the genetic circuits that are subsequently introduced, since the same control signal is used for all circuits and treatments. The control model is of the form

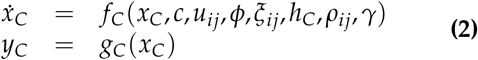

where *x*_*C*_(*t*) is a vector of unobserved state variables, *y*_*C*_(*t*) is the observable control signal, *u*_*ij*_ is the culture-specific treatment, *c*(*t*) is the culture density, *ϕ* are culture-independent control parameters and *ξ*_*ij*_ are culture-specific control parameters that characterize the *gene expression capacity* of the culture, which is described by a function *h*_*C*_(*c*(*t*), *ξ*_*ij*_, *ϕ*). The culture-specific growth parameters *ρ*_*ij*_ are incorporated into the model via the specific growth rate *γ*(*c*(*t*), *ρ*_*ij*_), which is used to model the effect of dilution due to growth.
3. A *target phase* characterizes the dynamics of the genetic circuit by comparing simulations of a *target model* with measurements of observable circuit outputs over time. The target model is of the form

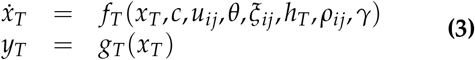

where *x*_*T*_ (*t*) is a vector of unobserved state variables, *y*_*T*_ (*t*) is a vector of observable genetic circuit outputs and *θ* are culture-independent circuit parameters. Similar to the control phase, the culture-specific growth parameters *ρ*_*ij*_ are incorporated into the model via the specific growth rate *γ*(*c*(*t*), *ρ*_*ij*_), while the culture-specific control parameters *ξ*_*ij*_ are incorporated by scaling the expression of all genes by a function *h*_*T*_ (*c*(*t*), *ξ*_*ij*_, *θ*).

Importantly, phases (1-3) represent three different measurements of the *same culture* containing a given circuit *i* under a given treatment *j*. The three phases are collectively referred to as *ratiometric* characterization. We also consider a variant of the method in which only the growth and target phases are used, referred to as *direct* characterization, which does not attempt to quantify between-culture variations in gene expression capacity. In this case *h*_*T*_ is set to 1 throughout.

We developed the notion of an *inference graph* (Supplementary Information Section S1.1) to formally represent the three phases of our method. An inference graph is defined as a *labelled directed acyclic site graph*, meaning that each edge has a label and a direction, the graph does not contain any cycles, and each node contains one or more sites:

- Each *node* in the inference graph represents a parameter inference over multiple cultures, indexed by their circuit *i* and treatment *j*, where each circuit is associated with a computational model and corresponding experimental measurements. Parameter inference seeks to minimize the deviation between the measurements and the model simulations (see Methods). The models in a given node share *global parameters* and typically represent a collection of similar circuits, such as circuits that differ only in their ribosome binding sites at specific locations.
- Each *site* in a node represents a circuit *i*, where the same model is assumed to be used consistently across all treatments *j* for that circuit. The site can also contain *local parameters* specific to that circuit.
- Each *edge* is labelled with a set of parameters that are *propagated* from the source to the target node, where each parameter is associated with a *propagation property* that denotes how the parameter is to be propagated. For example, if the property is *Fixed* then only the parameter value is propagated. If the property is *Normal* then the posterior marginal distribution of the parameter is fitted to a Normal distribution, which is then used as a prior for the parameter in the target node. Edges can occur between nodes or sites, to represent the propagation of global or local parameters, respectively.

The inference graph representing the three phases of ratiometric dynamic characterization is shown in Fig. 1D. The Growth node fits a growth model to measurements of culture density, resulting in growth parameters *ρ*_*ij*_ for each circuit *i* under treatment *j*. These parameters are then propagated to the corresponding sites in the Control and Target nodes. The Control node then fits a control model to measurements of the control signal, resulting in control parameters *ξ*_*ij*_ for the same circuit *i* under the same treatment *j*, together with parameters *ϕ* shared between the control models. The control parameters are then propagated to the corresponding sites in the Target node, which fits a target model to measurements of the target signals, resulting in parameters *θ* shared between the target models. The inference graph representing the two phases of direct dynamic characterization is shown in Fig. 1E. More generally, inference graphs allow parameter inference to be applied sequentially to circuits of increasing complexity, by reusing parameters shared between circuits in a consistent manner.

### Modeling gene circuits

Our dynamic characterization method is general in that it can support a range of modeling hypotheses. In this paper we focus on models of gene circuits derived from elementary chemical reactions, by making various assumptions about the timescales of different biological processes, culminating in only proteins and small molecules being explicitly described by ordinary differential equations (see Methods and Section S2).

Genetic devices are conveniently composable during circuit construction and model specification. However, the quantitative properties of parts and devices may not be. The rate of translation of mRNA to protein depends on the upstream ribosome binding site, while the rate of protein synthesis also depends on other factors from its surrounding context, such as the choice of upstream promoter, downstream terminator, position on the plasmid with respect to the origin of replication or other gene expression cassettes. Accordingly, one must hypothesize the rules governing whether a parameter can be shared when parts or devices are reused in other circuits. Here, we take the view that the rate of protein synthesis, which we modeled by treating transcription and translation as a single-step process (see Methods and Supplementary Information Section S2), depends on the associated promoter, ribosome binding site and protein coding region. Only when these three elements are the same are protein synthesis parameters shared. In contrast, we hypothesize that protein degradation is the same for a given protein in different genetic circuits. For inducible promoters, we assume that the regulation function parameters only depend on promoter sequence (and the concentration of the upstream regulator), and do not depend on down-stream ribosome binding site and protein coding region elements.

### Dynamic characterization case study

We demonstrated how our dynamic characterization method can be used to quantify the properties of genetic devices, by building and characterizing genetic circuits in *E. coli* that provide core functionality for engineering cell behavior at the population level. Our circuits make use of homoserine lactone (HSL) signaling components, including the Lux system from *V. fischeri* (28) and the Las system from *P. aeruginosa* (29), which have both been studied extensively in their natural contexts. They have also been used in synthetic biology contexts, either alone (30), in co-culture (31), or integrated into the same host organism (32).

In order to characterize HSL signaling devices, we designed a collection of synthetic gene circuits of increasing complexity, reusing parts in equivalent genetic contexts as much as possible (Fig. 2). For each circuit, a model was proposed that explicitly described the circuit behavior. These are summarized in the Methods section, with detailed derivations shown in Supplementary Information Section S2. In Fig. 2, code snippets show how devices can be programmatically specified as calls to modules that describe their behaviors. Circuits are defined by specifying which devices are included and by assigning models for cell culture growth and autofluorescence. Example comparisons of calibrated model simulations against measurements are shown in Fig. 2 for each circuit, and more extensive comparisons are shown in Supplementary Information Section S2.

**Fig. 2.**
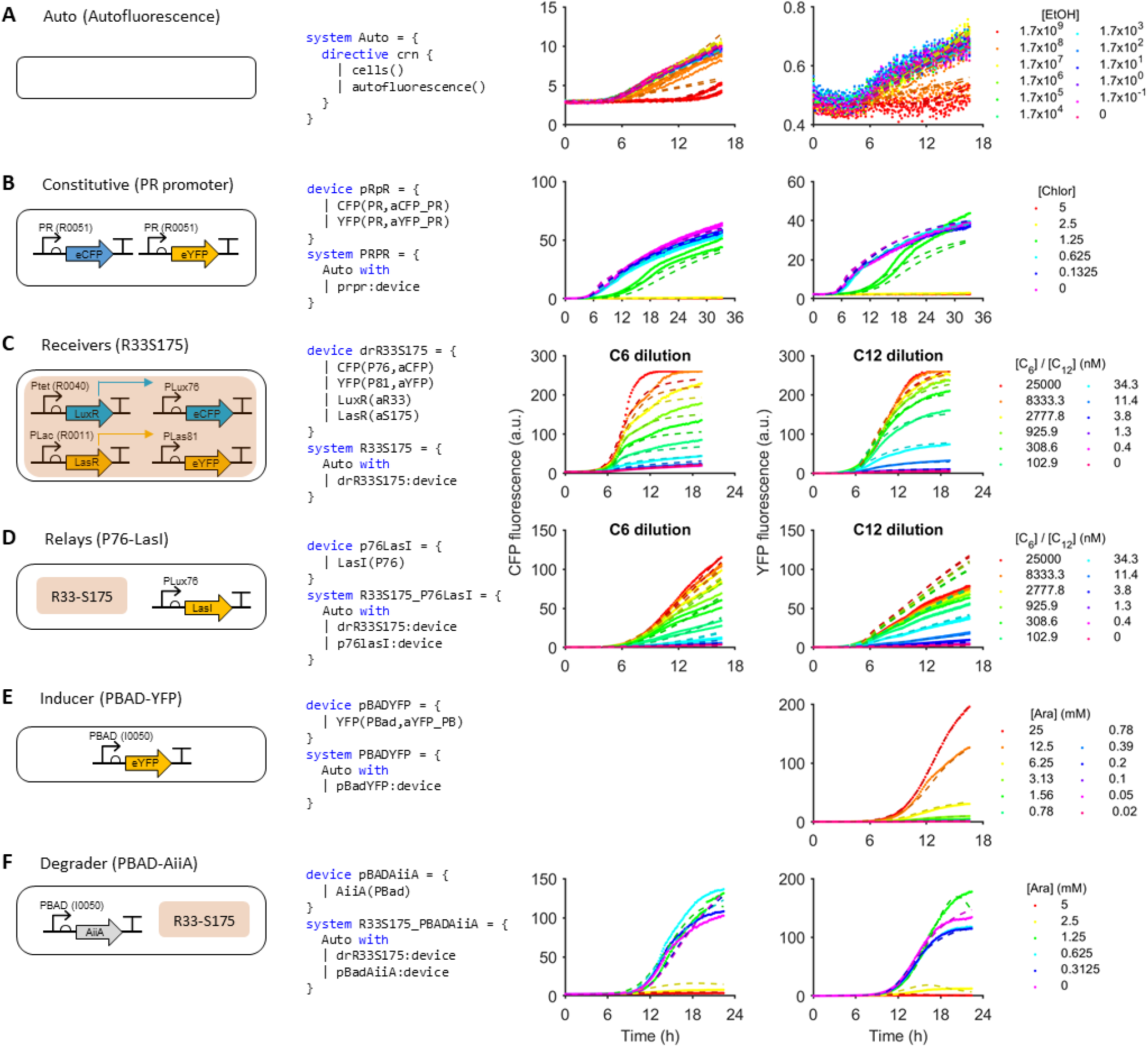
Composable design and characterization of HSL signaling parts and devices. Synthetic gene circuits are constructed and modeled in a modular way. In each row, the left column graphically depicts different circuits, the middle column shows device and circuit definitions as programs in the Genetic Engineering of Cells (GEC) language and the right column shows example measurements (solid lines) and calibrated model simulations (dashed lines). **A.** Autofluorescence characterized with cells expressing no eCFP or eYFP, but measured at 480 nm and 530 nm in response to varying concentrations of EtOH. The model of the Auto circuit specifies the cellular context through module calls `cells()` and `autofluorescence()`. **B.** Fluorescent protein degradation was characterized by expressing eCFP and eYFP with a constitutive promoter, treating with chloramphenicol, and measuring for a longer duration. The model of the PRPR device involves module calls `CFP()` and `YFP()`, which specify protein production, in addition to protein degradation and dilution (see Supplementary Information Section S1 for more details). The model of cell growth and autofluorescence is assumed to be equivalent to the model in A, and so is propagated to the system definition using the `with` keyword. **C.** HSL Receivers were characterized by measuring 4 variant circuits with different ribosome binding sites for LuxR and LasR expression, and treated with C6 or C12. Shown here is the R33-S175 circuit variant only. The model incorporates autofluorescence and fluorescent protein degradation from A and B. **D.** HSL Relays were characterized by inducibly expressing LasI in response to different concentrations of C6 via PLux81 (shown here) or LuxI in response to C12 via PLas81 (see Supplementary Information). The model incorporates elements of A, B and C. Notably, this system extends the `Auto` system using the `with` keyword, by combining the `drR33S175` and `p76lasI` devices. **E.** The PBAD promoter was characterized by inducing expression of eYFP with different concentrations of arabinose. The model incorporates elements of A and B. **F.** The HSL Degrader, PBAD-AiiA, was characterized by inducing its expression via PBAD, then measuring the response of the double receiver to different concentrations of HSL. Data-model comparisons are shown for the eCFP response to 25 *µ*M C6 (left panel) and the eYFP response to 25 *µ*M C12 (right panel) only.

We applied dynamic characterization both sequentially and simultaneously to the circuits introduced in Fig. 2, using both direct and ratiometric approaches. The inference graph for sequential direct characterization is shown in Fig. 3, while the graph for ratiometric characterization is shown in Fig. S1. Below we summarize the various stages of the inference graph in Fig. 3.

**Fig. 3.**
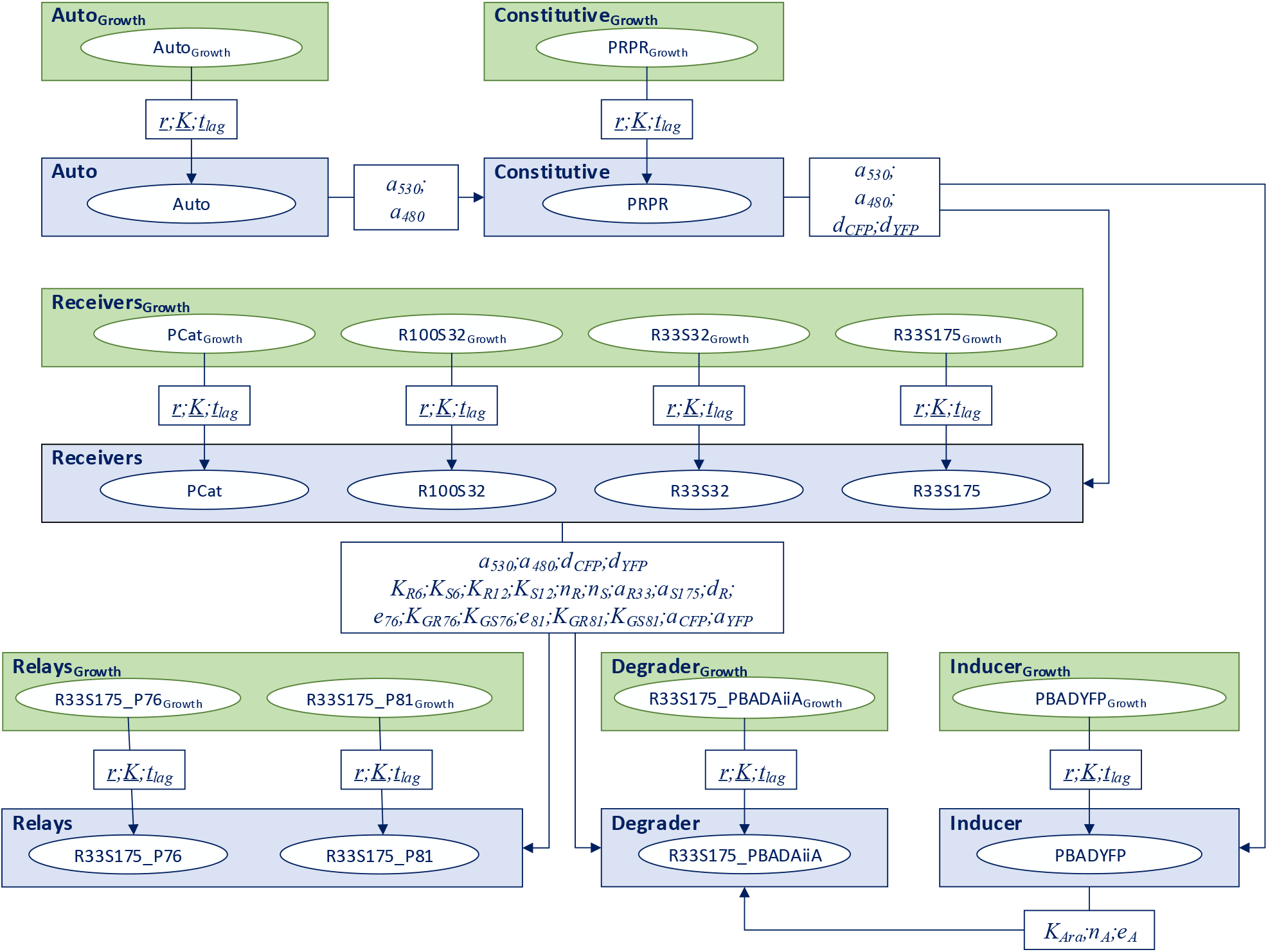
Inference graph for direct dynamic characterization of synthetic gene circuits. For conciseness, we allow edges from the same source node to share identical labels. The Auto_Growth_ node uses a growth model to infer growth parameters *r*, *K* and *t*_*lag*_ of the autofluorescence circuit, under different treatments. The parameters are propagated as vectors of equal length (shown as underlined), where the length of the vectors corresponds to the number of treatments. The parameters are propagated to the Auto node, which uses the Auto model to infer parameters *a*_530_ and *a*_480_ that characterise circuit autofluorescence. The posterior distributions of these parameters are then used as priors for the Constitutive node, which uses a model of constitutive gene expression to infer the parameters *d*_*CFP*_ and *d*_*YFP*_, together with updated posteriors of the autofluorescence parameters. The Receivers_*Growth*_ node infers growth parameters of four receiver devices simultaneously. The devices differ primarily in the ribosome binding sites used to drive expression of the two receiver proteins, LuxR and LasR. The Receivers node uses the corresponding growth parameters to infer parameters shared between the four receiver circuits. The Relays, Degrader and Inducer nodes are defined in a similar fashion, with parameters propagated as shown in the inference graph.

#### Autofluorescence

We started by characterizing cellular autofluorescence, to explicitly account for autofluorescence in circuits expressing eYFP and eCFP (Fig. 2A; Section S2.1). We pharmacologically perturbed the cells with EtOH, which above a threshold concentration led to very slow cell growth. We found that the per-cell rate of autofluorescence production at 480 nm, corresponding to eCFP measurements, was an order of magnitude higher than at 530 nm, corresponding to eYFP measurements. The autofluorescence parameters *a*_530_ and *a*_480_ inferred from these experiments were reused in subsequent circuits.

#### Constitutive promoter

Since both eCFP and eYFP were used in most of the synthetic gene circuits we constructed, we characterized their stability using a simple circuit (Fig. 2B; Section S2.2) expressing the eCFP and eYFP fluorescent proteins constitutively using the PR promoter, and measured their fluorescence for 36 hours. A model of the PRPR circuit incorporated autofluorescence terms, which shared information with the Auto circuit in Fig. 2A. The time-series measurements of PRPR enabled identification of the eCFP and eYFP degradation parameters *d*_CFP_ and *d*_YFP_.

#### Receivers

The first HSL devices we characterized were the Receivers, since these are also required to characterize the Relay and Degrader devices. We used the double receiver circuits developed in (32), which act as a dual reporter of 3-oxo-C6-HSL (C6) and 3-oxo-C12-HSL (C12; Fig. 2C; Section S2.3). This is achieved by the constitutive expression of LuxR and LasR and the inducible expression of fluorescent reporter proteins under the control of PLux promoter variants selected for orthogonal response to regulators composed of C6 bound to LuxR or C12 bound to LasR. In this way, eCFP reports the concentration of C6 and eYFP reports the concentration of C12, both across a broad range of concentrations. Because different promoters and ribosome binding sites are upstream of the coding sequences for eCFP and eYFP, as compared to the PRPR circuit, the parameters for their maximal production rate were assumed to take on new values. However, the degradation parameters were assumed to be the same as for PRPR, since the degradation rate is an intrinsic property of the protein. We used four different double receiver circuits, each incorporating a unique combination of promoter and ribosome binding sites in their LuxR and LasR expression cassettes. As explained in previous work, this enabled the effect of variable intracellular concentrations of LuxR and LasR to be modeled and therefore characterized quantitatively, which was critical for reducing chemical and genetic crosstalk (32).

#### Relays

We next characterized LuxI and LasI, the synthases of C6 and C12 respectively. HSL relay devices were designed that inducibly express the synthase of one signal in response to the other. HSL is produced intracellularly but quickly diffuses into the culture media, changing the signal concentration to which all cells are responding. This means that the promoter response to the synthesized signal is inherently time-varying and therefore cannot be accurately characterized using static methods. The first relay uses the C6-responsive pLux76 promoter to drive expression of LasI (abbreviated as P76-LasI), leading to intracellular production of C12 and subsequent induced expression of eYFP via the pLas81 promoter (Figs. 2D & S20B). The second uses the C12 responsive pLas81 promoter to drive expression of LuxI (abbreviated as P81-LuxI), leading to intracellular production of C6 and subsequent induced expression of eCFP (Fig. S20A). Again, reusing component characterization of the previously described circuits, we modeled time-series measurements of the response to different concentrations of C6 and C12 (Fig. 2D & S21), enabling quantification of the ability of LuxI and LasI to synthesize HSLs.

#### Inducer

We wished to use an orthogonal inducible system to drive expression of an HSL-degrading enzyme (see ‘Degrader’ below) but first needed an understanding of the parameters governing induction. To this end we characterized a simple circuit in which the properties of arabinose induction of the promoter PBAD could be quantified, by simply using PBAD to drive eYFP expression (Figs. 2E & S23). Incorporating the eYFP autofluorescence parameter from our Standard device, our model of the PBAD circuit closely described the experimental measurements in response to different concentrations of arabinose (Fig. S24) and enabled identification of the transfer function parameters for arabinose-induction, given by *K*_Ara_, *n*_A_ and *e*_A_ (Fig. S25).

#### Degrader

To characterize the HSL lactonase AiiA (33), we designed a circuit in which AiiA was placed under the control of the arabinose-inducible PBAD promoter (Fig. 2F), so that different intracellular levels of AiiA could be produced experimentally. Embedding the characterized PBAD and Receiver modules into the model of the AiiA circuit enabled modeling of the response to different concentrations of C6, C12 and arabinose. The parameters quantifying the shape of the transfer function of PBAD induction by arabinose were propagated from the characterization of the Arabinose circuit, though the parameter defining the maximal output 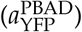 was not, because that is associated with a different downstream protein coding region. Of particular importance in reproducing the observed bulk fluorescence dynamics of the Degrader circuit was the incorporation of the effect on cell growth of high arabinose induction of AiiA, which was observable in the OD_600_ data, and therefore quantified during the cell growth phase of dynamic characterization (Figs. S28 & S30).

### Hypotheses about changes in host cell metabolism can be encoded in time-varying functions of gene expression capacity

As cells pass through different phases of growth, from lag to exponential to stationary, they undergo highly regulated changes in gene expression and metabolic state (34). These changes will have an effect on the expression of synthetic circuits and therefore on their dynamics. We observed such changes in gene expression in the eCFP and eYFP channels of our circuits that correlated with the onset of decreasing growth rate as measured by OD. Interestingly, such changes were much less prominent in the RFP channel (Fig. S36). We explored a number of hypotheses about the nature of those changes by evaluating the performance of different functional forms used to define gene expression capacity *h*_*T*_.

We considered several contrasting yet relatively simple assumptions about the functional form of the gene expression capacity *h*_*T*_ (*c*, *ξ*_*ij*_, *θ*) for the target phase (see Table 1). The simplest assumption was that gene expression capacity remains constant through time. However, because circuit activity changes over time, this assumption may break down if host cell resources become limiting when circuit activity increases. Furthermore, as bacterial cultures transition from exponential to stationary growth phase, broad changes in gene expression may need to be accounted for. For this we considered four alternative time-dependent functions (Table 1). The best-performing function differed from circuit to circuit but overall, the time-varying functions performed better than a constant gene expression capacity (Fig. S31).

**Table 1.**
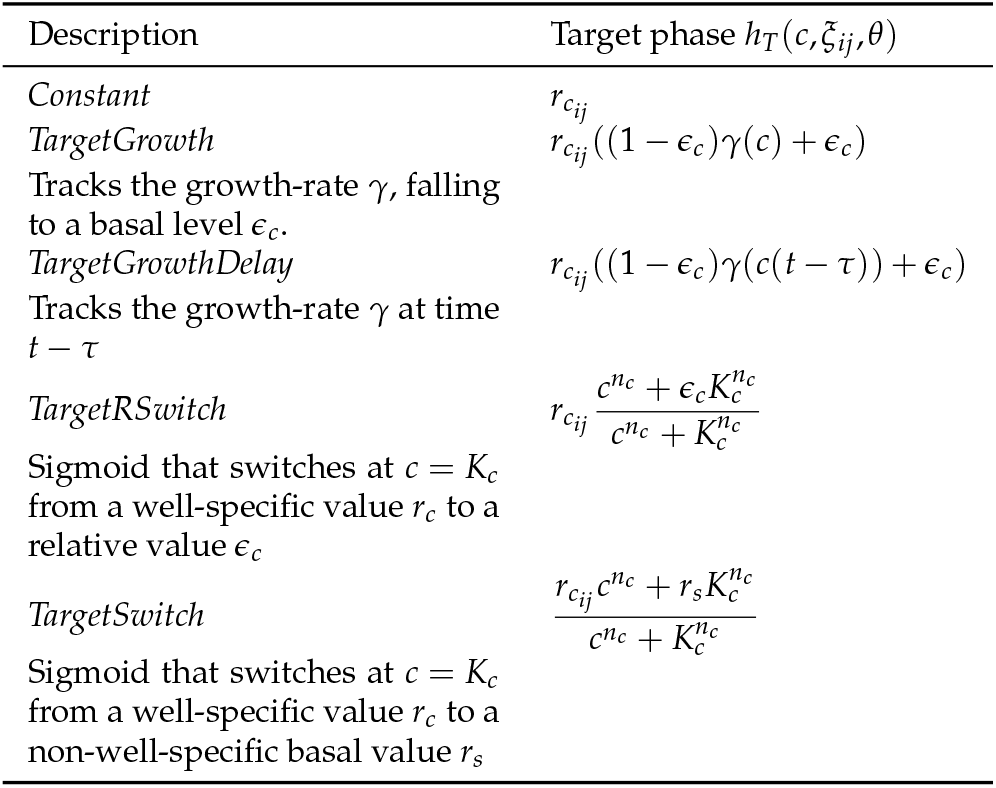
Functional forms used to define gene expression capacity *h*_*T*_(*c*). In all cases gene expression capacity *h*_*C*_ (*c*, *ξ*_*ij*_, *ϕ*) in the control phases uses the *Constant* hypothesis and its parameters *ξ*_*ij*_ = *r*_*cij*_ are inferred during this phase. Additional parameters in *h*_*C*_ (*c*, *ξ*_*ij*_, *θ*) may differ, and are therefore inferred during the target phase.

| Description | Target phase $h_T(c, \xi_{ij}, \theta)$ |
| --- | --- |
| *Constant* | $r_{c_{ij}}$ |
| *TargetGrowth*<br>Tracks the growth-rate $\gamma$, falling to a basal level $\epsilon_c$. | $r_{c_{ij}}((1 - \epsilon_c)\gamma(c) + \epsilon_c)$ |
| *TargetGrowthDelay*<br>Tracks the growth-rate $\gamma$ at time $t - \tau$ | $r_{c_{ij}}((1 - \epsilon_c)\gamma(c(t - \tau)) + \epsilon_c)$ |
| *TargetRSwitch*<br>Sigmoid that switches at $c = K_c$ from a well-specific value $r_c$ to a relative value $\epsilon_c$ | $r_{c_{ij}} \frac{c^{n_c} + \epsilon_c K_c^{n_c}}{c^{n_c} + K_c^{n_c}}$ |
| *TargetSwitch*<br>Sigmoid that switches at $c = K_c$ from a well-specific value $r_c$ to a non-well-specific basal value $r_s$ | $\frac{r_{c_{ij}} c^{n_c} + r_s K_c^{n_c}}{c^{n_c} + K_c^{n_c}}$ |

In each case, the culture-specific component 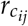 was inferred during the control phase, under a *Constant* hypothesis. Then, the non-culture-specific parameters (e.g. *є*_*c*_ in *TargetGrowth*) were inferred in the target phase alongside the target circuit parameters *θ*. As such, these hypotheses assume that the gene expression capacity for the chromosomally integrated control undergoes contrasting regulation from that of the plasmid-expressed devices. In principle, every gene expressed on the plasmid could potentially be regulated differently, which would require gene-specific functions or parameterizations. Here, we assume that the same function is applied to all genes in the target phase, to avoid a combinatorial increase in the number of parameters to infer.

### Sequential inference enables scalable dynamic characterization

Performing inference over databases containing multiple circuits and datasets can quickly become intractable as the database grows. Therefore, we have introduced the notion of an inference graph to describe how a large inference problem over multiple circuits can be decomposed into a sequential, graph-based inference strategy. In this way, when a new dataset is added to a database previously containing *n* datasets, inference can be performed on just the new dataset, as opposed to the complete database of *n* + 1 datasets. As such, sequential inference provides a scalable approach to an otherwise combinatorially intractable inference strategy. However, our sequential inference strategy provides only an approximation of the full (simultaneous) inference problem.

To determine the validity of this approximation, we compared the maximum log-likelihood score of the simultaneous characterization with the sum of the maximum log-likelihood scores over the 6 phases of the sequential characterization strategy (Fig. 4A & S31). This revealed a clear preference for sequential inference, most likely due to the flexibility afforded to model parameters when propagated from one inference problem to the next. To assess the extent to which parameters took advantage of this flexibility, we plotted the marginal distributions for each parameter resulting from inference of the 6 problems (Fig. S33 & S34). We found that the parameters associated with autofluorescence and fluorescent protein degradation were relatively stable as they were passed through the graph. Whereas parameters associated with the binding of LuxR/LasR to HSL signals and the modified PLux promoters varied by up to an order of magnitude or more. Our interpretation is that this reflects the inherent correlation between these parameters and the parameters associated with other HSL signaling circuits, characterized in inference problems downstream of the double receiver (see Fig. 3). The amount of variation also was greater in the TargetSwitch hypothesis than TargetGrowth hypothesis, which is likely due to TargetSwitch having more parameters associated with *h*_*T*_ to infer, and thus more potential for correlation and non-identifiability.

**Fig. 4.**
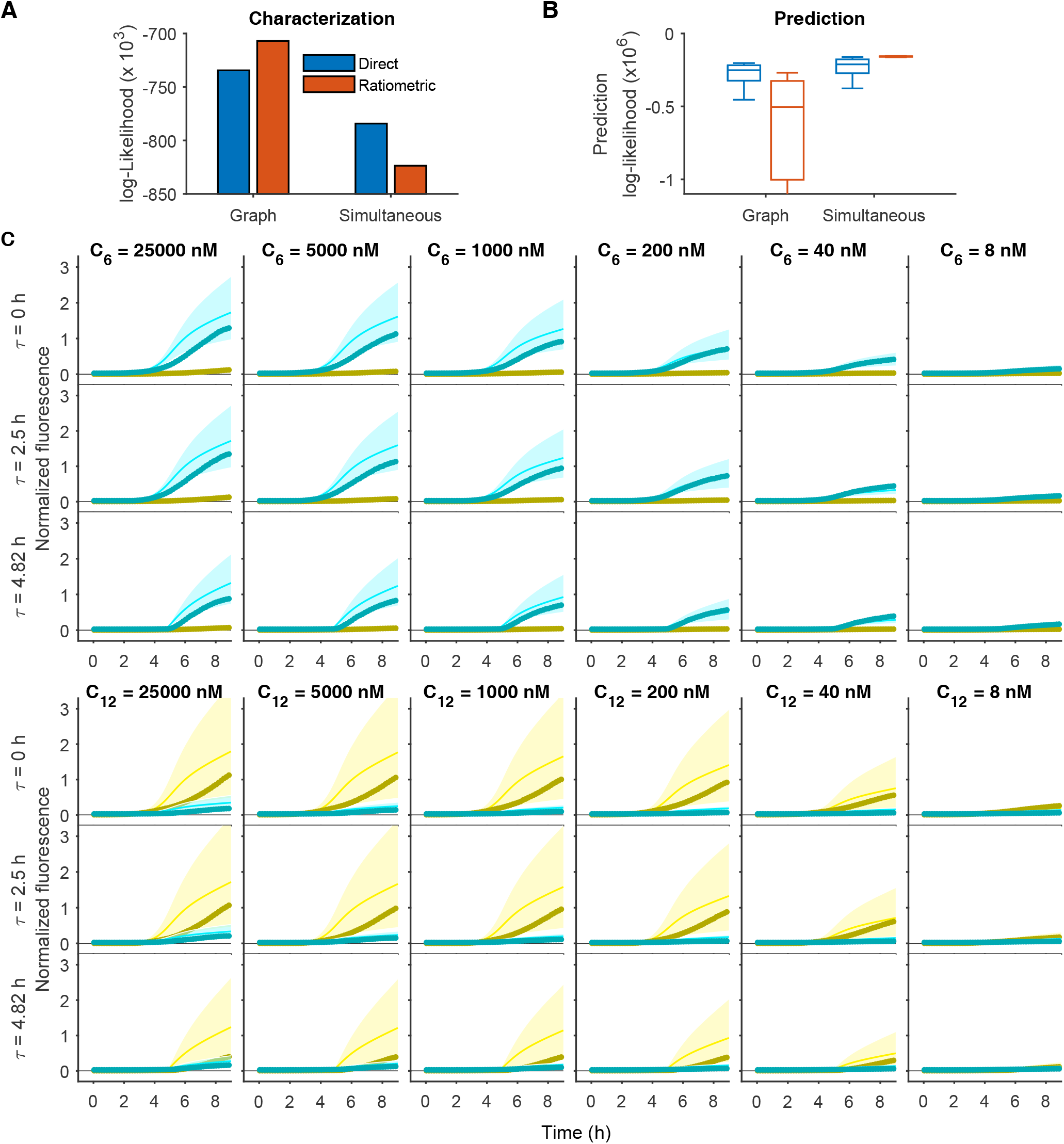
Prediction of unseen data. The predictive capabilities of each model are compared against data measuring the R33-S175 double receiver device in response to HSL treatment at different time-points. **A.** The maximum likelihood scores of dynamic characterization applied to six circuits is compared for graph-based and simultaneous inference, additionally comparing direct and ratiometric dynamic characterization. Shown here is the TargetSwitch hypothesis, while other hypotheses are compared in Fig. S31. **B.** The posterior predictive distribution for the test data in panel C is compared for the TargetSwitch models in panel A. The box-plots indicate the interquartile range of samples from the marginalized parameter posteriors. **C.** Comparison of the posterior predictive distribution for the TargetSwitch model using the direct graph method. In each column, the HSL treatment is indicated at the top, and in each row the time of HSL addition is noted on the left. eYFP and eCFP data are shown as thick darker lines (yellow and cyan respectively), with model simulations depicted as the mean (thin solid lines) and 95% credibility intervals as the shaded region.

We found that for sequential inference, in general, ratiometric dynamic characterization performed better than direct dynamic characterization in the training phase (Fig. 4A & S31G), though this preference was reversed in the simultaneous strategy (Fig. 4A & S31H). When considering the breakdown of these scores across different nodes of the inference graph, we found that PRPR, the Relays and some hypotheses for the double receiver did not benefit from the ratiometric control, but in fact led to worse fits to the data. Ideally, the ratiometric approach should be at least as good as the direct approach, which is equivalent to the ratiometric control parameters all being set to the same value. As such, the conclusions here suggest that the inferred ratiometric control parameters can sometimes inappropriately quantify gene expression capacity. Further investigation is required to determine how to more broadly specify these parameters.

### Dynamic characterization enables prediction of dynamic behavior

To test the ability of our modeling framework to predict dynamic behaviors, we measured the R33-S175 double receiver circuit in a regime previously not encountered in the training phase. The HSL treatments were applied at different times *τ* after the initialization of the assay and the commencing of measurement. This allowed us to observe how otherwise untreated cells respond to step-like perturbations. As expected, there was a lag induced in the accumulation of eYFP and eCFP expression (Fig. 4).

To quantify the predictive capability of each model hypothesis, we ran step 1 (and step 2 for ratiometric) of the dynamic characterization method. We then propagated the cell density parameters (and gene expression capacity parameters as appropriate) to a target phase. However, in the target phase, rather than running MCMC-based parameter inference, we sampled the marginal parameter posterior distributions to produce a posterior predictive distribution (see Methods). We chose to marginalize at this point, as opposed to sampling from the joint posterior of the double receiver inference results. This is because in general, we would seek to predict circuits where the underlying parameters were characterized across a range of other circuits, which means that in this method, the statistical correlations would be unavailable for sampling. Marginalization will introduce an approximation error, most likely resulting in greater uncertainty, and so using it here provides a conservative estimate of the uncertainty in the predictive distribution. We also marginalized the parameter posteriors for predicting based on the simultaneous method, for fairness.

We performed two analyses of the posterior predictive distribution. First, we calculated the log-likelihood for each sample, and summarized these distributions for each model hypothesis considered (Fig. 4B & S32). We then plotted the distribution of dynamical behaviors of the predictions for a selection of hypotheses. In Fig. 4C, we show the TargetSwitch hypothesis for the sequential application of direct dynamic characterization, which achieved a tight posterior predictive distribution. In this case, the general trend of most of the data is well reproduced. In Fig. S35, we show the TargetSwitch hypothesis again, but inferred from the simultaneous application of ratiometric dynamic characterization. Here, the predictions show a feature not exhibited by the measurements, which is that eYFP expression increases faster in the delayed HSL treatment case of *τ* = 4.82 h, whereas the measurements show significantly reduced expression compared with *τ* = 0 h and *τ* = 2.5 h. We found that this was due to the ratiometric control (mRFP1 measurements) showing increased expression, which led to higher values of the *r*_*c*_ parameters, which when propagated to the double receiver model, resulting in higher rates of eYFP and eCFP production (Fig. S36).

### Software implementation

Our method has been implemented in a dynamic characterization (DC) software package that enables a convenient specification of model hypotheses using a domain-specific programming language for genetic engineering of living cells (GEC), which allows models to be expressed as Chemical Reaction Networks (CRNs). Related software tools that incorporate MCMC-based parameter inference for CRNs have previously been used in research for molecular programming and immune system modelling (35, 36). Here, we implemented an inference graph framework that supports both simultaneous and sequential dynamic characterization, and incorporated this into the GEC language. We also developed a wrapper program that can extract time-series data from a specified format, adapted in our particular case for BMG plate-readers, and sub-select those wells that contain measurements for a device of interest. This enables a user to specify which devices to characterize, provide links to the available data, and to generate and evaluate a set of inference problems that implements our dynamic characterization method. After each phase of the method, results files are written that enable the computation to be paused and resumed (check-pointing), so that a dependency graph can be navigated conveniently. Detailed information about how to use the dynamic characterization software is provided in Supplementary Information Section S1.

## Discussion

Quantitative biological knowledge is difficult to systematize as it is often unclear clear how measured quantities relate to each other when factors such as genetic context and measurement techniques vary between experiments. Mathematical models provide a framework for explicitly encoding those relationships, allowing an internally consistent body of knowledge to be built, even when the systems being investigated are too large and complex to reason about directly. The use of mathematical models to make design decisions in the construction of synthetic gene circuits has been a central goal of the field of synthetic biology since its inception (6–8). To make this a viable approach, methods are required that allow models to match experimental data and thereby encode the quantitative biological knowledge gained from experiments. Our method allows us to build increasingly complex genetic circuits by reusing genetic parts in new contexts and predicting their behavior, by encoding our knowledge from previous experiments in mathematical models. By taking a Bayesian approach we also capture our uncertainty about the values those parameters can take.

### Dynamic characterization methods allow for quantitative understanding of inherently time-varying circuits such as HSL senders and degraders

Most methods that characterize reporter gene expression assume the reporter to be in a steady state due to production, dilution, and degradation. While this simplification can be justified in many synthetic circuits, it is unsuitable for circuits that produce or degrade intercellular signals while responding to those signals. In a culture of one of these circuits, the population of cells will be growing and continuously producing and degrading signal, meaning that the signal concentration, and the resultant reporter expression in response, will be changing throughout the experiment. Our approach was designed to capture these dynamic effects and therefore allows a direct quantitative characterization of these types of circuits for the first time.

In our experiments with the HSL degrading enzyme AiiA, inducing expression even at fairly moderate levels in combination with a receiver circuit results in strong effects on growth and gene expression capacity (Figs. S28 & S30). These effects mean attempting to determine the efficacy of degradation under various induction conditions directly from the raw data is extremely difficult. By applying our method, which can take into account both a decreased growth rate and altered gene expression capacity, we learn the rate at which HSLs are degraded for a particular level of expression of AiiA, which is consistent with both the PBAD induction measured with a YFP reporter and with our characterization of the receiver device.

### Sequential versus simultaneous inference

Our inference graph methodology supports both sequential and simultaneous inference, where simultaneous inference involves running a single inference problem over all circuits simultaneously. There are benefits and drawbacks to each.

Sequential inference enables scalable parameter inference with respect to a growing collection of genetic circuits and associated measurements. This is particularly important for circuits with large numbers of parameters, including culture-specific parameters that account for variations in cell growth and gene expression capacity. Sequential inference is achieved by propagating marginal parameter posteriors from one inference problem as the priors of another, for example using truncated Gaussian distributions (Fig. 1B-D), or by propagating maximum likelihood parameter estimates directly. Performing inference sequentially in multiple phases allows each phase to be computationally tractable. Since scientific investigation proceeds in iterations of hypothesis formulation and testing, sequential inference provides a practical solution for updating model parameters as new data arrive. In contrast, simultaneous inference can only incorporate new data by re-running inference over all existing data for all existing circuits. This becomes intractable as the number of circuits increases, due to the increased number of models to simulate and the number of samples needed for convergence.

Sequential inference enables model simulations to more closely match experimental observations, since there is flexibility of parameter values between circuits. However, this increased flexibility comes at the cost of increased uncertainty, which we observed to be greater in the posterior predictive distribution of unseen data (Fig. 4A). In contrast, simultaneous inference requires parameter values to be identical across all datasets, which can sometimes be difficult to reconcile.

Simultaneous inference enables parameters shared between multiple genetic circuits to be identified in a non-biased way. In contrast, sequential inference introduces two approximation errors, the first through marginalization of the posterior parameter distributions, and the second through approximation of these marginals with truncated Gaussian distributions, which may not be the optimal choice of parameter distribution in some cases. Therefore, one must be careful to only marginalize over MCMC chains that have reached the same local maximum of the likelihood function, and discard those stuck in other local maxima. In the case of propagating maximum likelihood estimates for parameters, such as cell growth and gene expression capacity, this leads to an under-representation of parameter uncertainties, which can in turn lead to inaccuracies in a Bayesian treatment of the target phase.

### Open versus closed loop modeling

Our models of genetic circuits assume a particular level of abstraction for describing the time-evolution of constituent components, by describing transcription and translation explicitly but ignoring host cell machinery (RNA polymerase, ribosomes, proteases, etc). More detailed models could offer an alternative starting point, but without measurements corresponding to careful manipulation of the host machinery, such models are likely to contain non-identifiable parameters and mis-specifications. Without explicit consideration of host cell resources, the way we have chosen to account for the feedback burden of circuit activity is open loop: each culture is modeled with independent parameters quantifying the rates of cell growth and gene expression. By breaking the loop in this way, our approach removes the need to speculate on the functional form of the feedback between a genetic circuit and the host cell. Such speculation would increase the complexity of models and increase the possibility of the feedback models compensating for misfits in the genetic circuit models, with poor mechanistic justification. Greater mechanistic understanding of the interaction between synthetic gene circuits and cellular burden would likely improve both parameterizations and predictions, but in the absence of such insight, open-loop arrangements such as the one proposed here might be preferable to truly closed-loop feedback models.

There are drawbacks to this kind of simplification, however. Most notably, transient changes in burden that are associated with changes in circuit activity may not be captured. This is the situation encountered in experiments with timed addition of HSL. In these experiments we might expect burden to increase dramatically at the point in time that HSL is added to the culture. A predefined function for gene expression capacity will not objectively take into account this switch in feedback burden. Indeed, the direct dynamic characterization method, which has no inter-culture variability in gene expression capacity, produces a better fit to the timed addition experiment (Fig. 4). Curiously, the RFP signal in these experiments is larger in later treatments but still well fit by a constant gene expression capacity (Fig. S36), which is not necessarily predicted by simple intuitions about the timing of burden. We also found that the direct dynamic characterization method scored more highly than the ratio-metric method for PRPR, the Relays and sometimes the double receiver, suggesting that gene expression capacity is not well accounted for by the ratiometric control in certain cases. One possibility may be due to differences in chromosomal and plasmid-derived gene expression. Alternatively, different circuits may induce different burdens on their hosts, which may or may not be observable in a single ratiometric control signal.

### Related and future work

Our approach is scalable in that inference can be performed on a new collection of related gene circuits measured simultaneously, without the need to re-run inference on all circuits in the library. However, as high throughput experimental methods continue to improve, the number of circuits that can be characterized simultaneously will become limited by the efficiency of sampling-based inference methods. Recent years have seen rapid improvement in Bayesian inference methodologies. Harnessing such improvements is a promising avenue for improving the scalability of genetic circuit characterization. For example, using stochastic gradient descent can enable more efficient approximate Bayesian inference (37). Beyond this, making explicit approximations about the distributions of parameter uncertainties (e.g. variational inference) can lead to efficient and more scalable inference schemes in the context of ODEs (38). To explore these ideas, preliminary work published in a conference proceedings has investigated the use of gradient-based inference in a variational framework (39). This approach demonstrates a substantial improvement in efficiency over MCMC, however it assumes that all parameters have Gaussian distributions and can so far only be applied to a single gene circuit with specified regions of variability. Future work is needed to generalise this work to multiple gene circuits. Such future improvements in inference algorithm efficiency could be readily incorporated into our Dynamic Characterization method.

The simple gene expression capacity functions we considered do not fully capture the complexity of the metabolic shift that occurs in the transition of a cell culture from exponential to stationary phase. This is evidenced by their inability to exactly match the shape of the eCFP and eYFP time courses. Future work could improve this fit by including more mechanistic understanding about the way gene expression is affected by metabolic shifts, or by taking a more ‘black box’ approach to generating a functional form that best fits the data (38). Machine learning approaches that combine mechanistic models where such mechanism is known with learned functions that describe unknown mechanisms seem uniquely suited for this kind of problem (39).

In summary, our method can be used to design increasingly complex gene circuits and predict their dynamic behaviors, by composing dynamically characterized genetic devices. This is achieved by encoding quantitative biological knowledge of device behavior using mathematical models parameterized by experimental data. By taking a Bayesian approach we also capture the uncertainty of model parameters, which could in future be used to determine the next experiments to perform to achieve a desired behavior.

## Methods

### Measurement of synthetic gene circuits in a microplate fluorometer

Plate fluorometer assays were conducted as previously described (32). Briefly, overnight cultures of cells containing constitutive chromosomal mRFP1 (32) and the plasmid construct of interest were diluted 1:100, grown to an OD of approximately 0.5, then diluted 1:1000 into M9 supplemented with 0.2% casamino acids and 0.4% glucose. 200*µ*l of culture was aliquoted into each well and measurements were taken every 10 min for 1,000-2,000 min in a BMG FLUOstar Omega plate fluorometer. 3-oxohexanoyl-homoserine lactone (C6-HSL, Cayman Chemicals) and 3-oxododecanoyl-homoserine lactone (C12-HSL, Cayman Chemicals) were dissolved to a concentration of 200 mM in DMSO then C6-HSL was diluted in supplemented M9 medium to the concentrations described, while C12-HSL, due to its limited solubility in aqueous media, was first diluted 1:50 in ethanol then diluted in supplemented M9 medium to the concentrations described. A 1M arabinose (Sigma) stock solution was made in water, filter sterilized, and diluted in supplemented M9 medium to the concentrations described. HSL receiver and sender plasmids were previously described in (32) and all other plasmids (summarized in Table S1) were constructed using Gibson Assembly (40) from parts obtained from the MIT Registry of Standard Biological Parts (http://parts.igem.org).

### Observer process

Because the microplate reader measurements monitor a whole culture growing over time, the observed fluorescence measurements are a sum of the single cell fluorescences in the culture. However, our models of synthetic gene circuits describe intra-cellular interactions, and therefore describe concentrations within each cell. Therefore, to compare the models with bulk fluorescence data, we use a nonlinear *observer* process. For each wavelength *w*, we describe bulk fluorescence variables as

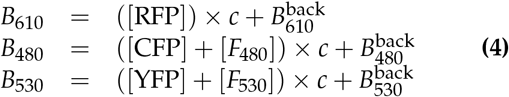

where [*F*_*w*_] is the per-cell autofluorescence at wavelength *w*, *c* is cell density (as above), and 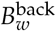 is the constant background fluorescence at wavelength. We use [RFP], [CFP] and [YFP] to denote the (modelled) intracellular concentrations of mRFP1, eCFP and eYFP respectively. Following these definitions, it remains to describe how to model the cell density (*c*) and the intracellular concentrations of fluorescent proteins ([RFP], [CFP] and [YFP]).

#### Modeling cell growth

We characterized the growth of each culture in a microtiter plate separately, to explicitly capture acceleration and deceleration of growth in response to variations in synthetic gene circuit activity and culture density. Absorbance measurements at 600 nm enable quantification of the optical density over time (OD_600_), providing a measure that is proportional to cell density (41).

To model cell density, we use the general equation

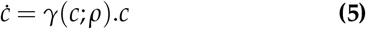

where *γ*(*c*; *ρ*) is the *specific growth rate* function that is parameterized by *ρ*. For the lag-logistic model, *γ* is defined by

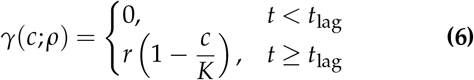

Here *r* is the per capita culture growth rate, *K* is the carrying capacity representing the maximum culture density and *t*_lag_ is the duration of the lag phase of bacterial growth. Accordingly, the parameters of cell growth for circuit *i* and culture *j* are given by *ρ*_*ij*_ = {*r*_*ij*_, *K*_*ij*_, *t*_lag,*ij*_}. The time-evolution of *c*(*t*) depends also on the initial cell density, *c*^0^:= *c*(*t* = 0), which is fixed to the intended dilution factor (0.2%) during preparation of the assay. In the measured OD_600_ signal, there is a non-cellular background signal, which we parameterize as a plate-level shared parameter *x*_*b*_. As a closed-form solution exists for the lag-logistic model, we can simply model the time evolution of the OD signal in culture *ij* as

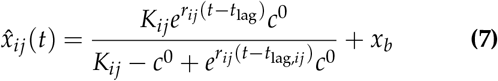

By assuming that the initial cell density is small, we have that *c*(0) ≈ 0, and so the background absorbance measurement will be approximately equal to the initial absorbance measurement. Therefore, we identify the parameter *x*_*b*_ as the average over the first time point of the OD signal in each well.

This leaves the identification of the well-specific parameters as a set of inference problems in which we seek to minimize the deviation between the data (*x*_*k*_) and simulation 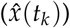. To obtain parameter estimates, we assume that the deviation between data and simulation is Gaussian-distributed with standard deviation *σ*. We seek to maximize the log-likelihood of producing the data in culture *ij* with parameters *ρ*_*ij*_

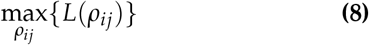

where *L*(*ρ*_*ij*_) is the sum of the log-probabilities of Gaussian observations for 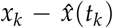, with variance *σ*^2^. For simple optimization problems over few parameters, it is possible to obtain good performance (both reliability and efficiency) using direct search methods such as the Nelder-Mead simplex algorithm (42). Here, we used the implementation in MathNet.Numerics (http://numerics.mathdotnet.com/) throughout this study for cell growth characterization.

### Control phase model derivation

We describe constitutive production of mRFP1 mRNA, followed by translation and fluorescent protein maturation by the following CRN:

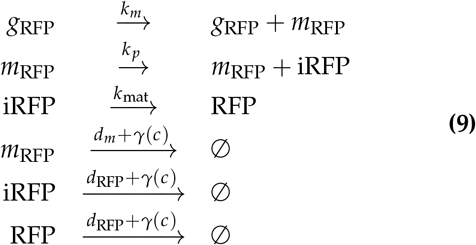

where *k*_*m*_, *k*_*p*_ and *µ*_*R*_ are transcription, translation and maturation of mRFP1 respectively, *γ* is the rate of dilution, and *d*_*m*_, *d*_RFP_ are degradation rates of mRNA and (mRFP1) protein.

From the reaction set in (Eq. 9), we can write down corresponding ordinary differential equations for the time evolution of the concentrations of each molecule, assuming mass action kinetics as follows:

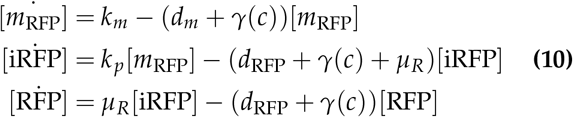

The dynamics of transcription and mRNA turnover are usually faster than translation and protein turnover, and therefore a separation of timescales argument can be applied to simplify the model equations (Eq. 10) as [*m*_RFP_]* ≈ *k*_*m*_/*d*_*m*_ +*γ*. By further assuming that the mRNA dynamics are faster than dilution, the above expression becomes constant, and can be incorporated into a single quantity incorporating the mRNA equilibrium and the translation rate *k*_*p*_.

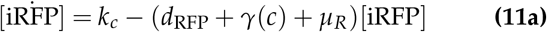

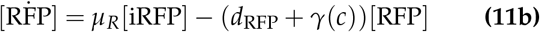

If the maturation time is faster than the turnover time for the fluorescent proteins, then an even simpler model can be derived:

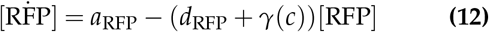

where *a*_RFP_ incorporates transcription, translation, fluorescent protein maturation and mRNA degradation rates. Since these factors are might be modulated by the growth phase and metabolic status of cell, *a*_RFP_ can be considered to represent the gene expression capacity, as introduced above. Accordingly, we introduce the open loop parameterization of the control circuit as

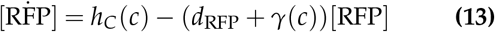

where gene expression capacity *h*_*C*_(*c*, *ξ*_*ij*_, *ϕ*) is parameterized by culture-specific parameters *ξ*_*ij*_ and global parameters *ϕ* (see Table 1 for options). The effect of dilution is described by the specific growth rate *γ*(*c*) (Eq. 6) and represents the decline in concentration as the volume of cells increases. Since we have characterized cell growth for each culture, we can use this to accurately model culture-specific dilution.

### Target phase models

Here, we summarize the derived equations for each (target phase) circuit considered in this study. Full derivation of these models can be found in Supplementary Information Section S2.

### Auto

The model of autofluorescence assumes that each cell produces material autofluorescent at the wavelengths corresponding to YFP and CFP measurement, and is subject to growth dilution in the usual way.

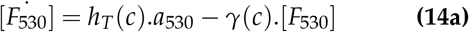

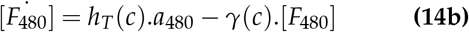

### Constitutive

The model of the Constitutive circuit incorporates autofluorescence (Eq. 14) and the following equations describing constitutive expression, degradation and growth dilution of eYFP and eCFP.

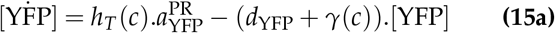

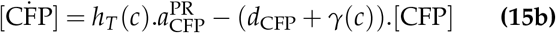

### Receivers

The receivers are a collection of 4 different circuits which differ in their ribosome binding sites upstream of LuxR and LasR expression. Accordingly, their models all have the same structure, but use RBS-specific parameters for LuxR and LasR synthesis.

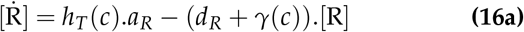

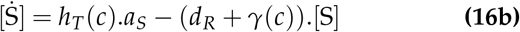

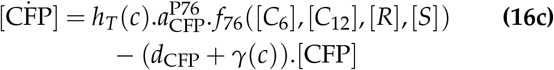

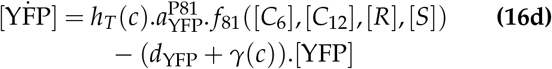

As above, the full models also incorporate the autofluorescence equations (Eq. 14).

### Relays

Two relay circuits are considered that each extend the R33-S175 Receiver circuit. The relay device P76-LasI expresses the C12 synthase LasI in response to C6 (via the P76 promoter), while the P81-LasI device expresses the C6 synthase LuxI in response to C12 (via the P81 promoter). Accordingly, their models have a common structure

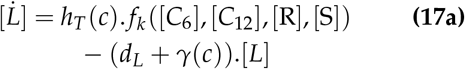

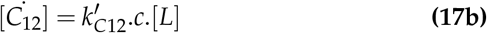

where *k* is P76 or P81, and *L* represents LuxI or LasI. Combining these equations with the equations for the R33-S175 circuit completes the description.

### Inducer

Arabinose-inducible expression of YFP uses a Hill regulation function *f*_PBAD_, with a maximal transcription rate that differs from maximal transcription of YFP of other promoters.

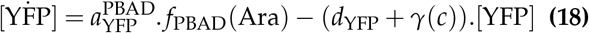

Combining these equations with the Auto circuit model completes the description.

### Degrader

The model of Arabinose-inducible AiiA uses the same Hill function as for the Inducer circuit, but maximal production of AiiA is divided out, as its concentration is not observed.

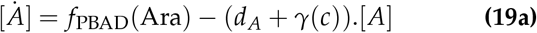

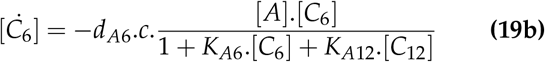

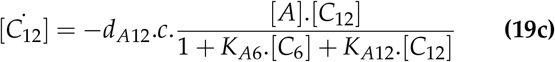

Combining these equations with the equations for the R33-S175 circuit completes the description.

### Markov chain Monte Carlo

In both the control phase and the target phase, we seek to infer the parameters of ODE models, given some observational data. For notational convenience, we define *θ* to be the vector of parameter values sought. In contrast to the optimization procedure in the cell growth phase, we use Markov chain Monte Carlo (MCMC) methods because we found that the Nelder-Mead algorithm performed poorly for inference problems over many parameters, frequently getting stuck in sub-optimal global optima of the cost function. MCMC methods are able to reduce, but not completely remove, the impact of this problem. MCMC methods also have the advantage of characterizing the uncertainty of parameter estimates, which can arise from several sources: measurement error, process error (molecular stochasticity) and model misspecification.

We used the Metropolis-Hastings algorithm as implemented in the Filzbach software (http://www.github.com/predictionmachines/filzbach) to perform MCMC parameterizations. This requires specifying a function that evaluates the log-likelihood score for a candidate parameter set *θ*, and prior distributions of each parameter, which encode our prior belief of its plausible values.

If we denote by *y*_*w,i,j*_ the bulk fluorescence measurements at wavelength *w* (*w* ∈ {480, 530, 610}) in well *i* (*i* = 1,…, *n*_*c*_) at time-points *t*_*j*_ (*j* = 1,…, *n*_*t*_), then we can define a log-likelihood function for wavelength *w* as

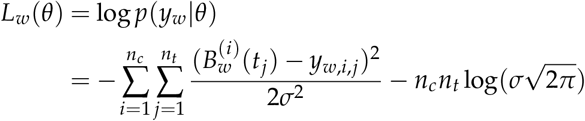

where *n*_*c*_ is the number of cultures (wells in the 96-well plate). In the control phase, we simply use *L*_610_ as the complete likelihood function, while in the target phase we use *L*_480_ + *L*_530_ when both eCFP and eYFP reporters are present in a circuit, and just one term otherwise. The parameter *σ* describes the standard deviation of the data. Here, we infer *σ* during application of MCMC.

Prior distributions encode our prior belief about the values of parameters. When characterizing a variable that has not previously been used in a model before, it can be difficult to know how to set the prior, so in this case we use a uniform distribution with wide bounds. This prevents the MCMC sampler from being swayed by an inappropriate value. When a parameter has been seen in another analysis, we propagate the marginal posterior from the previous analysis as a prior. To do this, we use truncated Gaussian distributions, where the mean and standard deviation are calculated from the previous MCMC samples, and the bounds are taken as the uniform prior bounds of that same analysis.

### Computing the posterior predictive distribution

To evaluate a model against unseen data, we approximate the posterior predictive distribution of the data, given our best estimates of the distributions of the model parameters. To do this, we formulate a model of the synthetic gene circuit being measured. In this article, we show calculations for the posterior predictive distribution of a circuit for which the parameters have already been characterized, so all are specified as truncated Gaussians, as described above. The predictions are formed by applying the cell growth phase and control phase as appropriate for dynamic characterization, but for the target phase we integrate over the prior. Accordingly, we approximate *p*(*y′|y*) by marginalizing over the posteriors of *p*(*θ|y*) to give approximate priors *π*_*θ*_ and then producing Monte Carlo samples *θ*_*k*_ ~ *π*_*θ*_, as

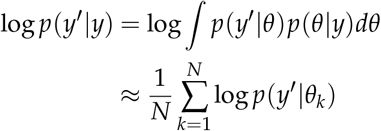

## ACKNOWLEDGEMENTS

The authors thank Fernan Federici and Timothy Rudge (Universidad Catolica de Chile) for useful discussions about ratiometric characterization. We also thank Ted Meeds (Microsoft Research) and Geoffrey Roeder (Princeton University) for discussions about Bayesian inference.

