## Supporting Information for "Scalable dynamic characterization of synthetic gene circuits"

Biological Computation group, Microsoft Research, Cambridge, CB1 2FB, UK

**S3 Supplementary Figures** **55**

**S4 Supplementary Tables** **59**

### S1 Method implementation using the GEC language

To implement our dynamic characterization method we extended the Genetic Engineering of Cells (GEC) language and its associated Chemical Reaction Network (CRN) language. This section describes the extensions made to the languages, together with a wrapper program that facilitates the preparation and scheduling of inference tasks across a graph (Fig. 3), including collecting multiple MCMC chains and using these to marginalize and approximate parameter distributions as Gaussian priors.

#### S1.1 Extensions to the CRN language

The main extension to the CRN language is the ability to define an Inference Graph, which provides a graph-based method to chain inference problems and consists of nodes and edges:

**nodes** denote inference problems. Each node consists of a set of CRN systems [1], where each system represents a model of a gene circuit, together with inference settings.

**edges** represent the propagation of parameters from one inference problem to another. Each edge is directed and consists of a source, a target and a list of parameters to be propagated together with the associated propagation property. The source indicates the node or system from which the posteriors of inferred parameters are propagated, while the target indicates the node or system in which the inferred parameters from the source are used as priors. Each parameter in an edge has a property, which indicates whether the full parameter distribution or the parameter value will be propagated.

The syntax of inference graphs is defined as follows, where italicized names denote syntax variables:

*Property* ::=<br>
| `Fixed`<br>
| `Normal`<br>
| `TruncatedNormal`

*Parameter* ::=<br>
| *Name*<br>
| *Name* = *Property*

*Edge* ::=<br>
| `edge` *Name* − > [$Parameter_1; ...; Parameter_N$] *Name*<br>
| `edge` *Name.Name* − > [$Parameter_1; ...; Parameter_N$] *Name.Name*

*Node* ::=<br>
`node` *Name* {<br>
`systems` = [$Name_1; ...; Name_N$];<br>
`settings` = *Inference_settings*;<br>
}

We use a dot notation to refer to a specific system at a given node in the graph. Note that we also allow edges between nodes directly, to indicate that the parameters are shared between all systems present at the node. When present, the inference graph is declared at the end of the CRN program.

#### S1.2 Extensions to the GEC language

To enable the generation of models for dynamic characterization we extended the GEC language [2], which was originally developed to design gene circuits using constraints expressed in terms of part types, part properties and reactions. The two main extensions are as follows:

- We extended the GEC language with the ability to specify devices. Previously, GEC only supported elementary parts including promoters (prom), ribosome binding sites (rbs), protein coding regions

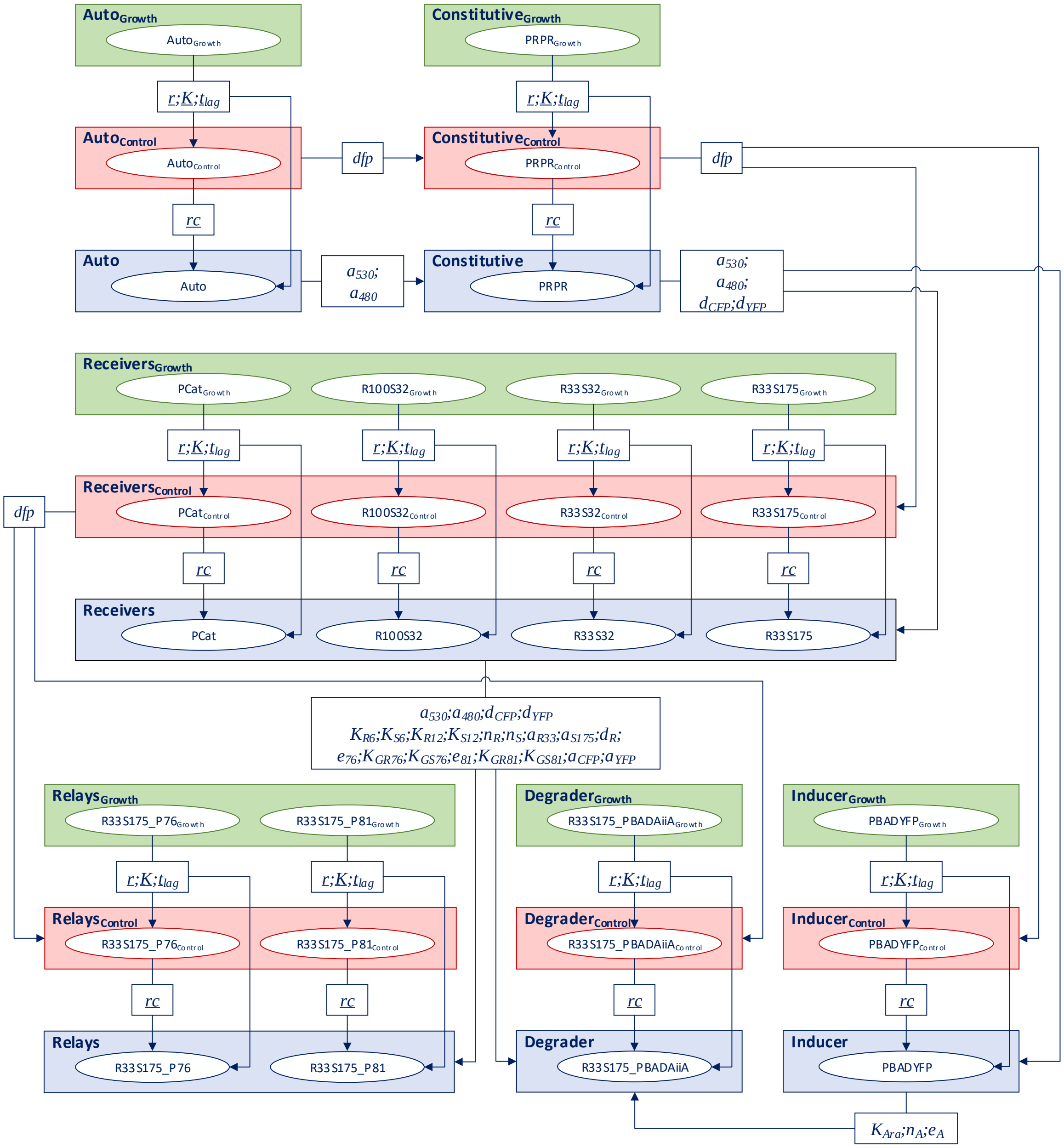

**Figure S1: Inference graph for ratiometric dynamic characterization of the case study presented in the main text.**

(pcr, also known as coding sequences), and terminators (ter). We extended the language to allow a `device` to be defined in terms of the basic parts types and other devices, and to be associated with a corresponding model expressed as a CRN.

- We extended the GEC language with the notion of an inference graph. In addition, we provided support for modules, directives and systems from the CRN language [1].

A GEC program can be divided into the following sections:

**directives** describe how a GEC program is executed and are specified using the `directive` keyword. We modified GEC to use CRN directives and introduced three directives specific to GEC:

- `RMRNADeg` - specifies the rate of mRNA degradation (the default value is 0.001)
- `crn` - specifies CRN instructions (module invocations, initial conditions and reactions), which are concatenated to the CRN instructions generated automatically from a GEC program.
- `override` - specifies CRN instructions as above, which replace those generated automatically from a GEC program.

**module definitions** are subroutines (or procedures) used in the CRN language and can have zero or more arguments. They are defined using the `module` keyword and consist of CRN instructions and other module invocations.

**device model definitions** are module definitions that describe the model associated to a given device. They are defined using the `device` keyword, followed by the device name with zero or more arguments. They consist of CRN instructions and other module invocations. The structural definition (parts and devices) of the device is specified in the GEC Parts and Device library.

**templates** are subroutines (or procedures) for the GEC language. Templates are defined using the `template` keyword and contain part types, part variables, part properties, normal reactions and transport reactions. Like modules, templates can have zero or more arguments.

**systems** are units of a GEC program that describe a specific gene circuit or biochemical process. Systems are structurally and functionally distinct from one another. A system can have directives, modules, module invocations, template invocations, parts and their properties, and GEC reactions whose scope is local to that system. A GEC program can also have the aforementioned program components outside a system. These components are considered to be *top level* and their scope is global to the entire GEC program. A system is defined using the keyword `system`.

**inference graphs** consist of nodes and edges that describe the inference tasks to be performed. The graph is defined using the `node` and `edge` keywords.

#### S1.3 Dynamic Characterization - GEC

We have extended GEC's graphical user interface to accommodate the extensions to the GEC and CRN languages introduced in the previous sections, including inference graphs, while preserving the original functionality of GEC ([2]). The latest version of the GEC web tool can now run inference problems using dynamic characterization.

Users can run inference tasks within GEC by specifying inference graphs such as the one defined for the Constant hypothesis in Listing 1, as shown in Fig. S2. GEC can also be used to visualize the inference graph (Fig. S3). Each system within a node of the graph can also be selected to view a graphical representation of the chemical reaction network model of the system.

GEC allows users to upload processed CSV (comma separated value) data files (see Supplementary Data for examples) and run dynamic characterization. Fig. S3 shows various tabs of the results generated in GEC by running dynamic characterization for the Constant hypothesis code from Listing 1. See the Supplementary Data files for instructions on running dynamic characterization within GEC.

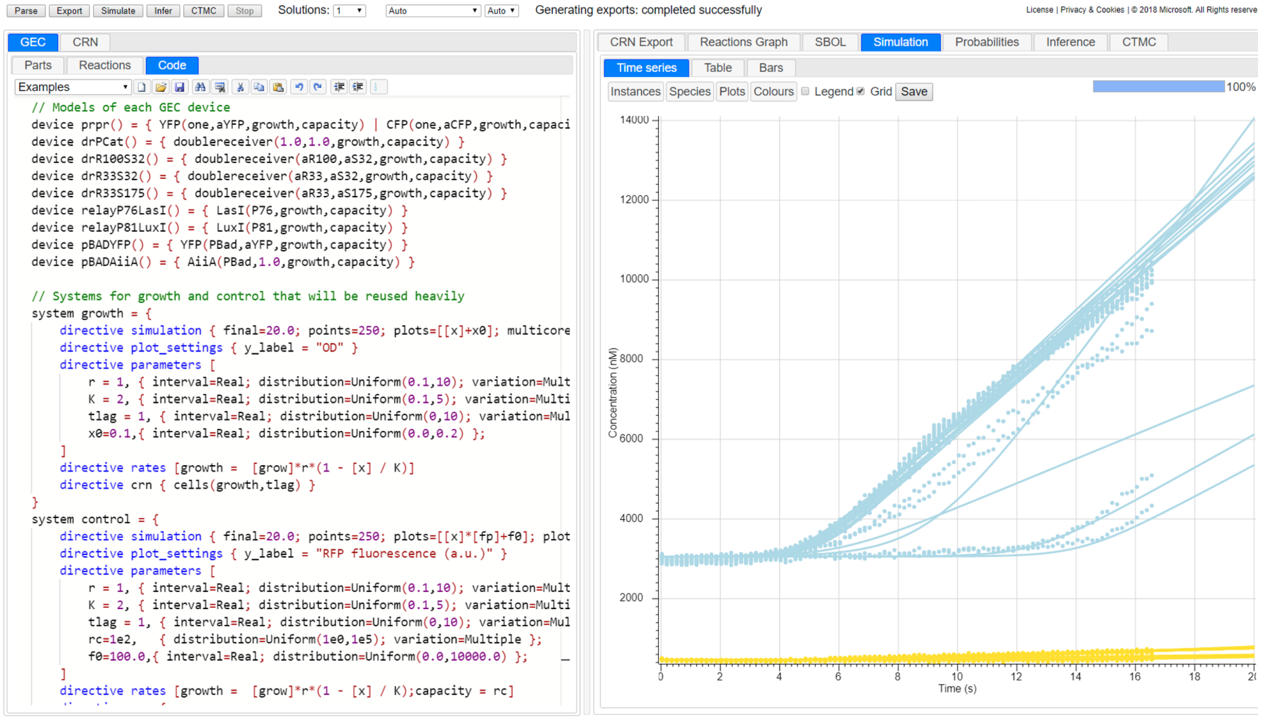

**Figure S2:** Screenshot of the GEC tool showing the GEC program used to evaluate the Constant hypothesis (left panel) and the simulations produced for the *Auto* system (right panel).

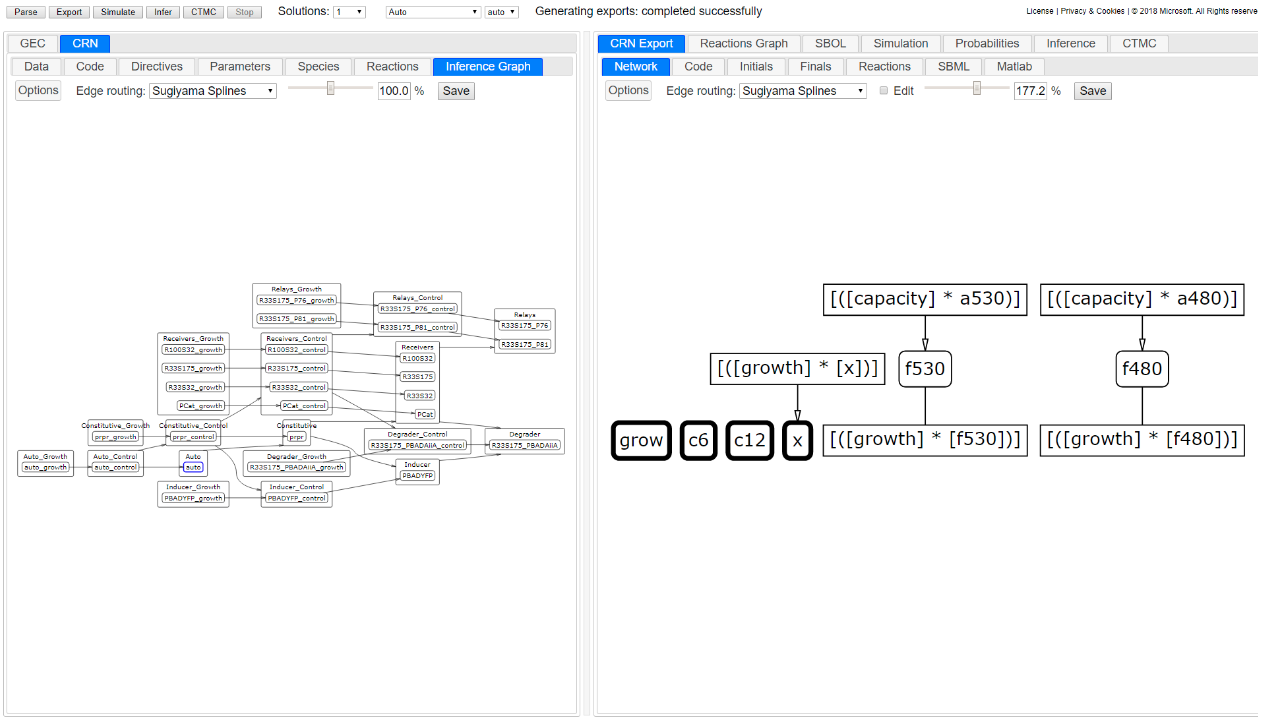

**Figure S3:** Screenshot of the GEC tool showing a graphical representation of the inference graph (left) for the Constant hypothesis and a graphical representation of the chemical reaction network (right) for the system selected in the inference graph (in this case the system *auto* within the node *Auto*).

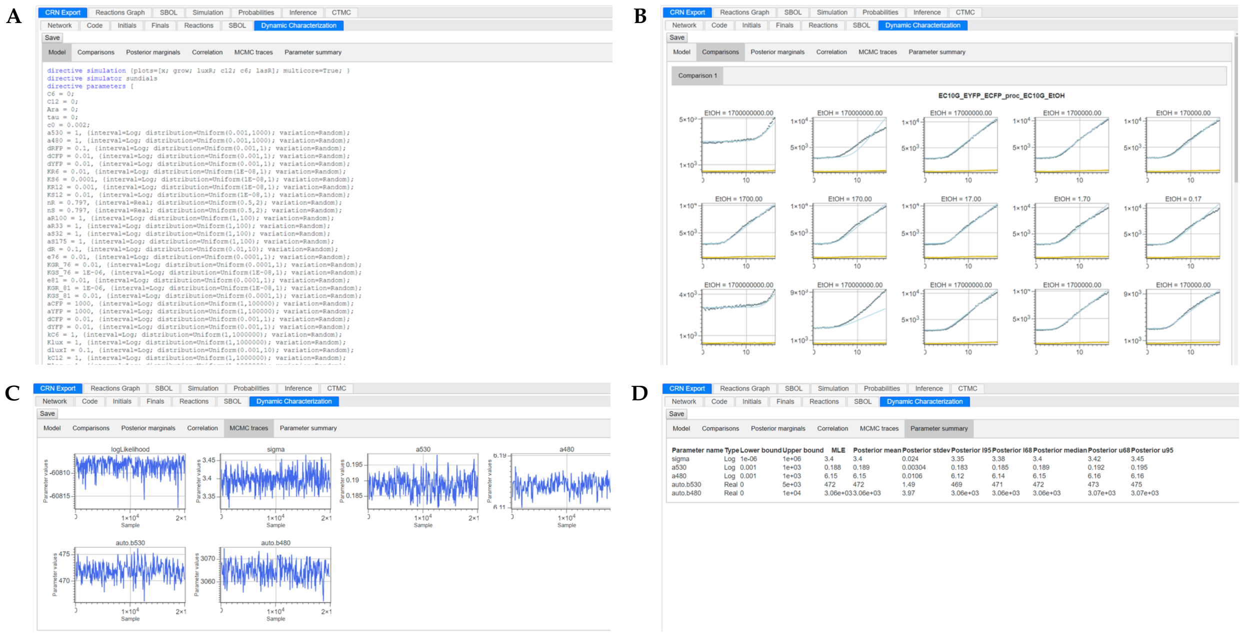

**Figure S4: Results of dynamic characterization generated in GEC.** **A**. The CRN model of a specific system within a node of the inference graph. **B**. Comparisons of the data and simulations of the maximum-likelihood parameter values produced for a specific node. **C**. Traces of the MCMC samples for a specific node of the inference graph. **D**. Summary of the inferred parameters at the end of the run.

#### S1.4 Dynamic Characterization - Command-line tool

To take advantage of high throughput computing, we make use of a command-line tool that wraps the dynamic characterization functionality of the GEC and CRN languages and enables individual inference nodes to be run in isolation. In this way, multiple MCMC chains can be generated simultaneously across multiple compute nodes of a CPU cluster or cloud computing infrastructure. It also facilitates the re-running of specific points within the inference graph, in case the model needs to be updated in any way. Examples include refining the underlying CRN of a system and modifying parameter prior distributions. This is highlighted in the figures S5, S6, and S7.

All models are written in the GEC and CRN languages, which have a default file format for observational data. The command-line tool simplifies the use of data and the specification of *sweeps*, which can be used to specify different treatment concentrations or input signals, by automatically extracting these from a more structured data file (see Supplementary Data for examples). To specify how the observational data is to be loaded for each model and each inference node, a YAML configuration file must be loaded. An example YAML file is the following:

```
homedir: ..\..\..\
outdir: GEC
hypothesis: Constant_Ratiometric
cellmodel: laglogistic
preprocess: individual
files: [proc_EC10G_EtOH]
inputs: [EtOH]
nodes:
- { node: Auto_Growth, signal: OD, targets: [ {device: EC10G, model: growth} ] }
- { node: Auto_Control, signal: RFP, targets: [ {device: EC10G, model: control} ] }
- { node: Auto, signal: YFP_CFP, targets: [ {device: EC10G, model: auto} ] }
```

In the command-line tool, devices and associated measurements are specified for inclusion, and a name is given to an inference problem. A YAML file must be provided that specifies how to load the data files appropriately, including the name(s) of the measurement signal(s) used in a given inference node, and the names of the devices associated with each model in that inference node. As an example, one can specify the characterization of the growth phase of the Auto circuit by calling:

```
DynamicCharacterization.exe graph --node Auto_Growth --yaml Ratiometric --hypothesis TargetSwitch
--seed 0
```

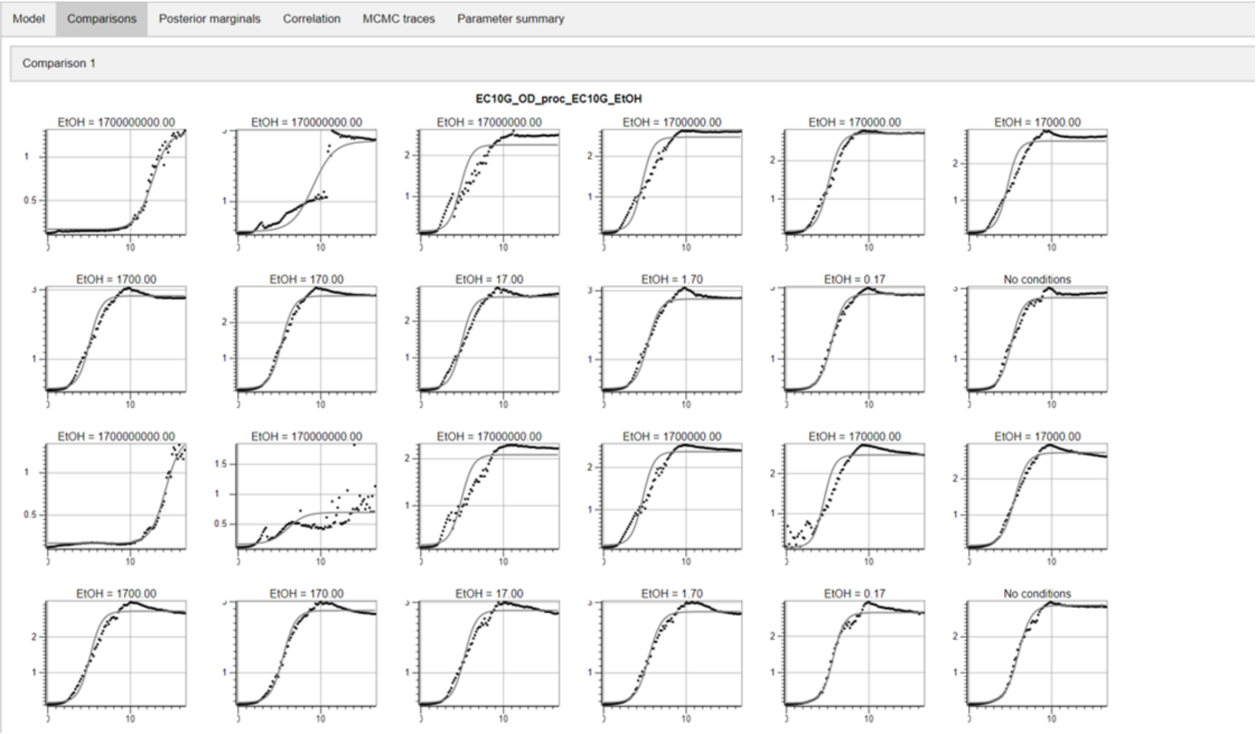

**Figure S5: Comparisions produced by the command-line tool after implementing dynamic characterization on the inference node - Auto Growth of the Constant hypothesis**

The code in Listing 1 shows the complete GEC code used to evaluate the Constant hypothesis. In the following section (S2), the model of each circuit is explained in detail.

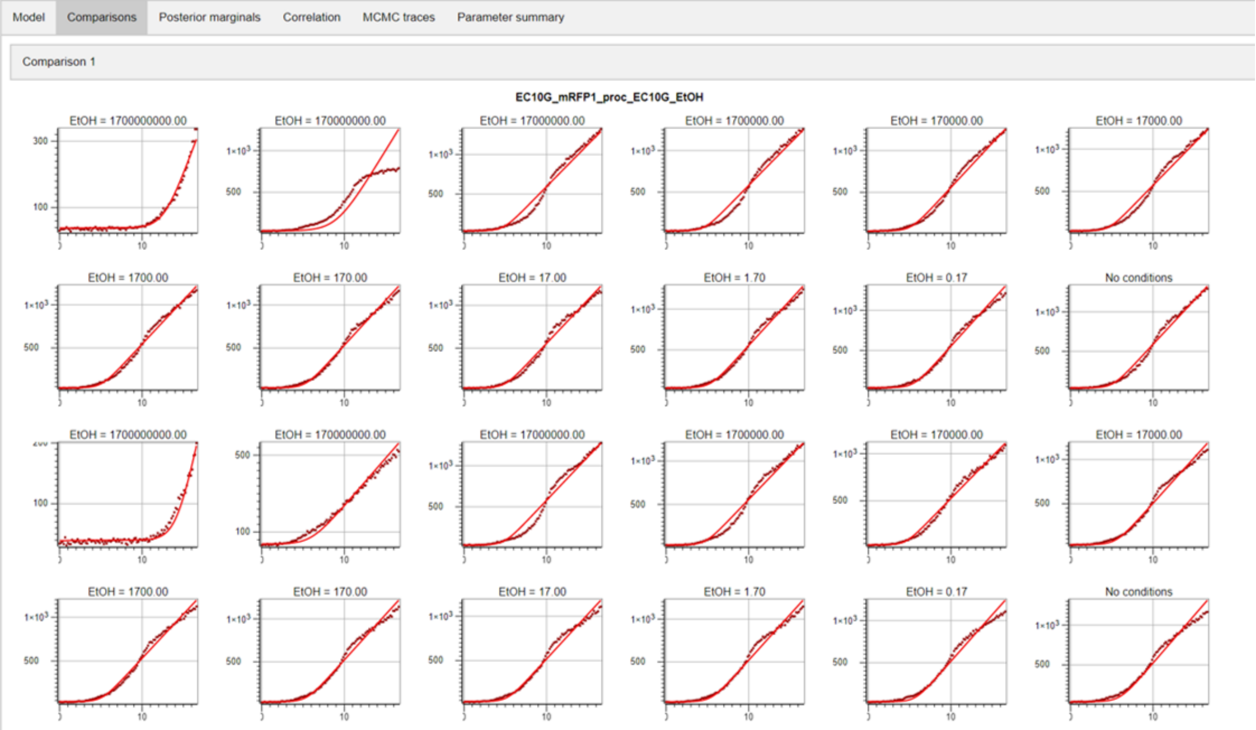

**Figure S6: Comparisions produced by the command-line tool after implementing dynamic characterization on the inference node - Auto Control of the Constant hypothesis**

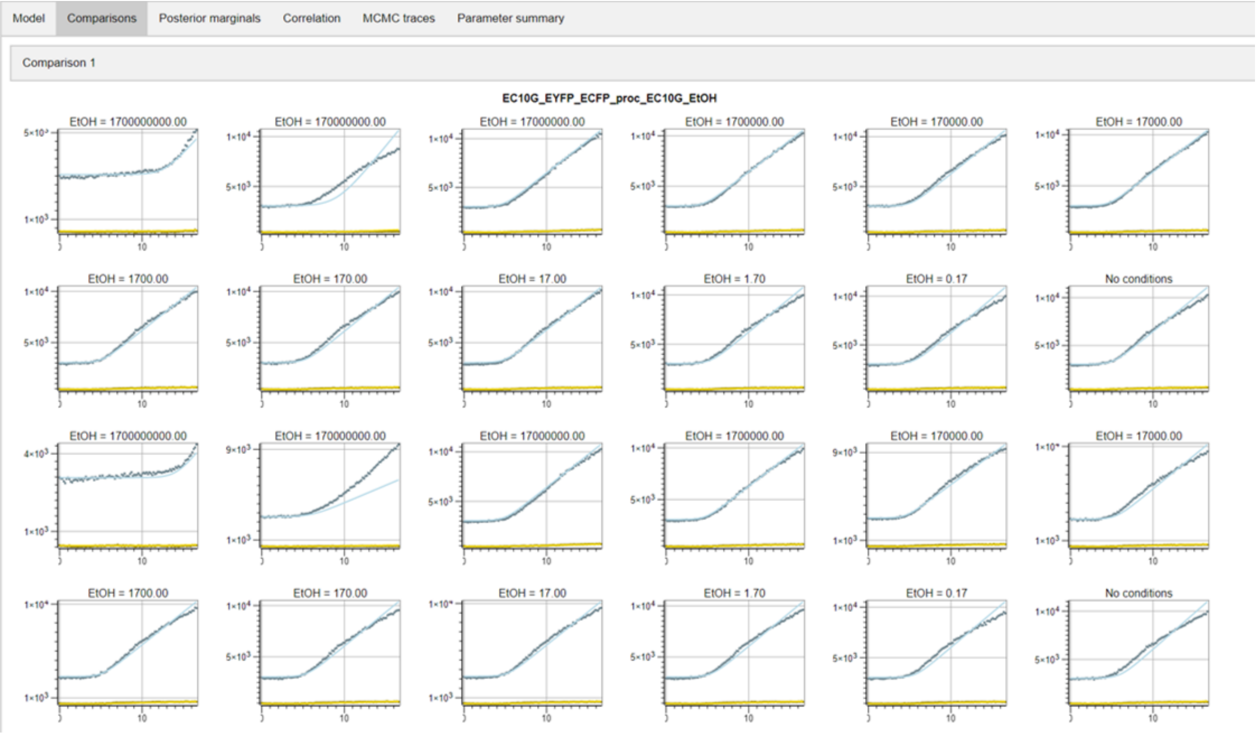

**Figure S7: Comparisions produced by the command-line tool after implementing dynamic characterization on the inference node - Auto of the Constant hypothesis**

**Listing 1:** Complete code for Constant hypothesis

```
directive simulator sundials
directive plot_settings { x_label = "Time (h)"; y_label = "Fluorescence (a.u.)" }
directive parameters [
    // Treatments
    C6=0.0; C12=0.0; Ara=0.0; tau=0.0;
    // Cell growth
    c0 = 0.002;
    // Autofluorescence
    a530=1e0,  { distribution=Uniform(1e-3,1e3) };
    a480=1e0,  { distribution=Uniform(1e-3,1e3) };
    // FP
    dRFP=0.1, { distribution=Uniform(1e-3,1e0) };
    dCFP=1e-2, { distribution=Uniform(1e-3,1e0) };
    dYFP=1e-2, { distribution=Uniform(1e-3,1e0) };
    // Double receiver
    KR6=1e-2,  { distribution=Uniform(1e-8,1e0) };
    KS6=1e-4,  { distribution=Uniform(1e-8,1e0) };
    KR12=1e-3, { distribution=Uniform(1e-8,1e0) };
    KS12=1e-2, { distribution=Uniform(1e-8,1e0) };
    nR=0.797,  { interval=Real; distribution=Uniform(0.5,2.0) };
    nS=0.797,  { interval=Real; distribution=Uniform(0.5,2.0) };
    aR100=1.0, { distribution=Uniform(1e0,1e2) };
    aR33=1.0,  { distribution=Uniform(1e0,1e2) };
    aS32=1.0,  { distribution=Uniform(1e0,1e2) };
    aS175=1.0,  { distribution=Uniform(1e0,1e2) };
    dR=0.1,    { distribution=Uniform(1e-2,1e1) };
    //dS=0.1,  { distribution=Uniform(1e-2,1e2) };
    e76=1e-2,  { distribution=Uniform(1e-4,1.0) };
    KGR_76=1e-2,{ distribution=Uniform(1e-4,1e0) };
    KGS_76=1e-6,{ distribution=Uniform(1e-8,1e0) };
    e81=1e-2,  { distribution=Uniform(1e-4,1.0) };
    KGR_81=1e-6,{ distribution=Uniform(1e-8,1e0) };
    KGS_81=1e-2,{ distribution=Uniform(1e-4,1e0) };
    aCFP=1e3,  { distribution=Uniform(1e0,1e5) };
    aYFP=1e3,  { distribution=Uniform(1e0,1e5) };
    dCFP=1e-2, { distribution=Uniform(1e-3,1e0) };
    dYFP=1e-2, { distribution=Uniform(1e-3,1e0) };
    // Relay P81-LuxI
    kC6=1e0, { distribution=Uniform(1e0,1e6) };
    Klux=1.0, { distribution=Uniform(1e0,1e6) };
    dluxI=0.1,{ distribution=Uniform(1e-3,1e1) };
    // Relay P76-LasI
    kC12=1e0, { distribution=Uniform(1e0,1e6) };
    Klas=1.0, { distribution=Uniform(1e0,1e6) };
    dlasI=0.1,{ distribution=Uniform(1e-3,1e1) };
    // PBad
    KAra=1.0, { distribution=Uniform(1e-2,1e2) };
    nA=1.0,  { interval=Real; distribution=Uniform(0.5,3.0) };
    eA=0.1,  { interval=Real; distribution=Uniform(0.0,0.5) };
    // AiiA
    dA6=1e-1, { distribution=Uniform(1e-3,1e1) };
    dA12=1e-1, { distribution=Uniform(1e-3,1e1) };
    daiiA=0.1, { distribution=Uniform(1e-3,1e1) };
]
directive simulation {multicore=True}
directive inference { burnin=200000; samples=200000; thin=200; partial = true; noise_model =
    proportional }
directive rates [
```

```
    growth = [grow]*r*(1 - [x] / K);
    capacity = rc;
    boundLuxR = [luxR]^2 * ((KR6*[c6])^nR + (KR12*[c12])^nR) / ((1.0 + KR6*[c6] + KR12*[c12])^nR);
    boundLasR = [lasR]^2 * ((KS6*[c6])^nS + (KS12*[c12])^nS) / ((1.0 + KS6*[c6] + KS12*[c12])^nS);
    P76 = (e76 + KGR_76*[boundLuxR] + KGS_76*[boundLasR]) / (1.0 + KGR_76*[boundLuxR] +
        KGS_76*[boundLasR]);
    P81 = (e81 + KGR_81*[boundLuxR] + KGS_81*[boundLasR]) / (1.0 + KGR_81*[boundLuxR] +
        KGS_81*[boundLasR]);
    PBad = (Ara^nA+eA*KAra^nA)/(Ara^nA+KAra^nA);
    one = 1.0;
]

directive sweeps [
    sweep_R33S175_PBADAiiA = [(C6,C12,Ara) = [(25000,0,5); (25000,0,2.5); (25000,0,1.25);
        (25000,0,0.625); (25000,0,0.3125); (25000,0,0); (0,25000,5); (0,25000,2.5); (0,25000,1.25);
        (0,25000,0.625); (0,25000,0.3125); (0,25000,0); (5000,0,5); (5000,0,2.5); (5000,0,1.25);
        (5000,0,0.625); (5000,0,0.3125); (5000,0,0); (0,5000,5); (0,5000,2.5); (0,5000,1.25);
        (0,5000,0.625); (0,5000,0.3125); (0,5000,0); (1000,0,5); (1000,0,2.5); (1000,0,1.25);
        (1000,0,0.625); (1000,0,0.3125); (1000,0,0); (0,1000,5); (0,1000,2.5); (0,1000,1.25);
        (0,1000,0.625); (0,1000,0.3125); (0,1000,0); (200,0,5); (200,0,2.5); (200,0,1.25);
        (200,0,0.625); (200,0,0.3125); (200,0,0); (0,200,5); (0,200,2.5); (0,200,1.25);
        (0,200,0.625); (0,200,0.3125); (0,200,0); (40,0,5); (40,0,2.5); (40,0,1.25); (40,0,0.625);
        (40,0,0.3125); (40,0,0); (0,40,5); (0,40,2.5); (0,40,1.25); (0,40,0.625); (0,40,0.3125);
        (0,40,0); (8,0,5); (8,0,2.5); (8,0,1.25); (8,0,0.625); (8,0,0.3125); (8,0,0); (0,8,5);
        (0,8,2.5); (0,8,1.25); (0,8,0.625); (0,8,0.3125); (0,8,0); (1.6,0,5); (1.6,0,2.5);
        (1.6,0,1.25); (1.6,0,0.625); (1.6,0,0.3125); (1.6,0,0); (0,1.6,5); (0,1.6,2.5);
        (0,1.6,1.25); (0,1.6,0.625); (0,1.6,0.3125); (0,1.6,0); (0,0,5); (0,0,2.5); (0,0,1.25);
        (0,0,0.625); (0,0,0.3125); (0,0,0); (0,0,5); (0,0,2.5); (0,0,1.25); (0,0,0.625);
        (0,0,0.3125); (0,0,0)]];
    sweep_C6C12_double = [(C6,C12) = [(25000,0); (8333.33333333333,0); (2777.77777777778,0);
        (925.925925925926,0); (308.641975308642,0); (102.880658436214,0); (34.2935528120713,0);
        (11.4311842706904,0); (3.81039475689681,0); (1.27013158563227,0); (0.423377195210757,0);
        (0,0); (0,25000); (0,8333.33333333333); (0,2777.77777777778); (0,925.925925925926);
        (0,308.641975308642); (0,102.880658436214); (0,34.2935528120713); (0,11.4311842706904);
        (0,3.81039475689681); (0,1.27013158563227); (0,0.423377195210757); (0,0);(25000,0);
        (8333.33333333333,0); (2777.77777777778,0); (925.925925925926,0); (308.641975308642,0);
        (102.880658436214,0); (34.2935528120713,0); (11.4311842706904,0); (3.81039475689681,0);
        (1.27013158563227,0); (0.423377195210757,0); (0,0); (0,25000); (0,8333.33333333333);
        (0,2777.77777777778); (0,925.925925925926); (0,308.641975308642); (0,102.880658436214);
        (0,34.2935528120713); (0,11.4311842706904); (0,3.81039475689681); (0,1.27013158563227);
        (0,0.423377195210757); (0,0)]];
    sweep_C6C12 = [(C6,C12) = [(25000,0); (8333.33333333333,0); (2777.77777777778,0);
        (925.925925925926,0); (308.641975308642,0); (102.880658436214,0); (34.2935528120713,0);
        (11.4311842706904,0); (3.81039475689681,0); (1.27013158563227,0); (0.423377195210757,0);
        (0,0); (0,25000); (0,8333.33333333333); (0,2777.77777777778); (0,925.925925925926);
        (0,308.641975308642); (0,102.880658436214); (0,34.2935528120713); (0,11.4311842706904);
        (0,3.81039475689681); (0,1.27013158563227); (0,0.423377195210757); (0,0)]];
    sweep_relays = [(C6,C12) = [(25000,0); (8333.333333,0); (2777.777778,0); (925.925926,0);
        (308.641975,0); (102.880658,0); (34.293553,0); (11.431184,0); (3.810395,0); (1.270132,0);
        (0.423377,0); (0,0); (0,25000); (0,8333.333333); (0,2777.777778); (0,925.925926);
        (0,308.641975); (0,102.880658); (0,34.293553); (0,11.431184); (0,3.810395); (0,1.270132);
        (0,0.423377); (0,0); (25000,0); (8333.333333,0); (2777.777778,0); (925.925926,0);
        (308.641975,0); (102.880658,0); (34.293553,0); (11.431184,0); (3.810395,0); (1.270132,0);
        (0.423377,0); (0,0); (0,25000); (0,8333.333333); (0,2777.777778); (0,925.925926);
        (0,308.641975); (0,102.880658); (0,34.293553); (0,11.431184); (0,3.810395); (0,1.270132);
        (0,0.423377); (0,0)]];
    sweep_PBAD_eYFP = [Ara = [25; 0.78; 12.5; 0.39; 6.25; 0.195; 3.125; 0.0975; 1.5625; 0.04875;
        0.78125; 0.024375]];
    sweep_EC10G_dataset0 = [(EtOH,chlor,C6,C12,Ara) = [(1700000000,0,0,0,0); (170000000,0,0,0,0);
```

```
        (17000000,0,0,0,0); (1700000,0,0,0,0); (170000,0,0,0,0); (17000,0,0,0,0); (1700,0,0,0,0);
        (170,0,0,0,0); (17,0,0,0,0); (1.7,0,0,0,0); (0.17,0,0,0,0); (0,0,0,0,0);
        (1700000000,0,0,0,0); (170000000,0,0,0,0); (17000000,0,0,0,0); (1700000,0,0,0,0);
        (170000,0,0,0,0); (17000,0,0,0,0); (1700,0,0,0,0); (170,0,0,0,0); (17,0,0,0,0);
        (1.7,0,0,0,0); (0.17,0,0,0,0); (0,0,0,0,0); (1700000000,0,0,0,0); (170000000,0,0,0,0);
        (17000000,0,0,0,0); (1700000,0,0,0,0); (170000,0,0,0,0); (17000,0,0,0,0); (1700,0,0,0,0);
        (170,0,0,0,0); (17,0,0,0,0); (1.7,0,0,0,0); (0.17,0,0,0,0); (0,0,0,0,0);
        (1700000000,0,0,0,0); (170000000,0,0,0,0); (17000000,0,0,0,0); (1700000,0,0,0,0);
        (170000,0,0,0,0); (17000,0,0,0,0); (1700,0,0,0,0); (170,0,0,0,0); (17,0,0,0,0);
        (1.7,0,0,0,0); (0.17,0,0,0,0); (0,0,0,0,0)]];
    sweep_PRPR_dataset1 = [(EtOH,chlor,C6,C12,Ara) = [(0,5,0,0,0); (0,5,0,0,0); (0,2.5,0,0,0);
        (0,2.5,0,0,0); (0,1.25,0,0,0); (0,1.25,0,0,0); (0,0.625,0,0,0); (0,0.625,0,0,0);
        (0,0.3125,0,0,0); (0,0.3125,0,0,0); (0,0,0,0,0); (0,0,0,0,0)]];
]

module CFP(P,a,growth,capacity) = {
    ->[[capacity]*a*[P]] cfp |
    cfp ->{dCFP} |
    cfp ->[[growth]*[cfp]]
}
module YFP(P,a,growth,capacity) = {
    ->[[capacity]*a*[P]] yfp |
    yfp ->{dYFP} |
    yfp ->[[growth]*[yfp]]
}
module LuxR(aR,growth,capacity) = {
    ->[[capacity]*aR] luxR |
    luxR ->{dR} |
    luxR ->[[growth]*[luxR]]
}
module LasR(aS,growth,capacity) = {
    ->[[capacity]*aS] lasR |
    lasR ->{dR} |
    lasR ->[[growth]*[lasR]]
}
module LuxI(P,growth,capacity) = {
    ->[[capacity]*[P]] luxI |
    luxI ->{dluxI} |
    luxI ->[[growth]*[luxI]] |
    ->[kC6*[capacity]*[x]*[luxI]/(1+[luxI]/Klux)] c6
}
module LasI(P,growth,capacity) = {
    ->[[capacity]*[P]] lasI |
    lasI ->{dlasI} |
    lasI ->[[growth]*[lasI]] |
    ->[kC12*[capacity]*[x]*[lasI]/(1+[lasI]/Klas)] c12
}
module AiiA(P,aI,growth,capacity) = {
    ->[[capacity]*aI*[P]] aiiA |
    aiiA ->{daiiA} |
    aiiA ->[[growth]*[aiiA]] |
    c6 -> [[x]*dA6*[c6]*[aiiA]] |
    c12 -> [[x]*dA12*[c12]*[aiiA]]
}
module cells(growth,tlag) = {
    init x c0 | init grow 1 @ tlag |
    init c6 C6 @ tau | init c12 C12 @ tau |
    ->[[growth]*[x]] x
}
```

```
module autofluorescence(growth,capacity) = {
    ->[[capacity]*a530] f530 |
    f530 ->[[growth]*[f530]] |
    ->[[capacity]*a480] f480 |
    f480 ->[[growth]*[f480]]
}
module doublereceiver(aR,aS,growth,capacity) = {
  LuxR(aR,growth,capacity) |
  LasR(aS,growth,capacity) |
  YFP(P81,aYFP,growth,capacity) |
  CFP(P76,aCFP,growth,capacity)
}

// Models of each GEC device
device prpr() = { YFP(one,aYFP,growth,capacity) | CFP(one,aCFP,growth,capacity) }
device drPCat() = { doublereceiver(1.0,1.0,growth,capacity) }
device drR100S32() = { doublereceiver(aR100,aS32,growth,capacity) }
device drR33S32() = { doublereceiver(aR33,aS32,growth,capacity) }
device drR33S175() = { doublereceiver(aR33,aS175,growth,capacity) }
device relayP76LasI() = { LasI(P76,growth,capacity) }
device relayP81LuxI() = { LuxI(P81,growth,capacity) }
device pBADYFP() = { YFP(PBad,aYFP,growth,capacity) }
device pBADAiiA() = { AiiA(PBad,1.0,growth,capacity) }

// Systems for growth and control that will be reused heavily
system growth = {
    directive simulation { final=20.0; points=250; plots=[[x]+x0]; multicore=True}
    directive plot_settings { y_label = "OD" }
    directive parameters [
        r = 1, { interval=Real; distribution=Uniform(0.1,10); variation=Multiple };
        K = 2, { interval=Real; distribution=Uniform(0.1,5); variation=Multiple };
        tlag = 1, { interval=Real; distribution=Uniform(0,10); variation=Multiple };
        x0=0.1,{ interval=Real; distribution=Uniform(0.0,0.2) };
    ]
    directive rates [growth = [grow]*r*(1 - [x] / K)]
    directive crn { cells(growth,tlag) }
}
system control = {
    directive simulation { final=20.0; points=250; plots=[[x]*[fp]+f0]; plotcolours=["#FF0000"];
         multicore=True }
    directive plot_settings { y_label = "RFP fluorescence (a.u.)" }
    directive parameters [
        r = 1, { interval=Real; distribution=Uniform(0.1,10); variation=Multiple };
        K = 2, { interval=Real; distribution=Uniform(0.1,5); variation=Multiple };
        tlag = 1, { interval=Real; distribution=Uniform(0,10); variation=Multiple };
        rc=1e2,  { distribution=Uniform(1e0,1e5); variation=Multiple };
        f0=100.0,{ interval=Real; distribution=Uniform(0.0,10000.0) };
    ]
    directive rates [growth = [grow]*r*(1 - [x] / K);capacity = rc]
    directive crn {
        cells(growth,tlag)
        | fp ->[[growth]*[fp]]    // Dilution
        | ->[[capacity]] fp       // Transcription/translation
        | fp ->{dRFP}                 // Degradation
    }
}
system target = {
    directive simulation { final=20.0; points=250; plots=[[x]*([yfp]+[f530])+b530;
         [x]*([cfp]+[f480])+b480]; plotcolours=["#FFDF00"; "#ADD8E6"]; multicore=True }
```

```
    directive parameters [
        r = 1, { interval=Real; distribution=Uniform(0.1,10); variation=Multiple };
        K = 2, { interval=Real; distribution=Uniform(0.1,5); variation=Multiple };
        tlag = 1, { interval=Real; distribution=Uniform(0,10); variation=Multiple };
        rc=1e2,  { distribution=Uniform(1e0,1e5); variation=Multiple };
        b530=1e3, { interval=Real; distribution=Uniform(0.0,5e3) };
        b480=1e3, { interval=Real; distribution=Uniform(0.0,1e4) };
    ]
    directive rates [growth = [grow]*r*(1 - [x] / K);capacity = rc]
    directive crn { cells(growth,tlag) | autofluorescence(growth,capacity) }
}

//Autofluorescence
system auto_growth = { growth with
    directive simulation {sweeps=[sweep_EC10G_dataset0] }
    directive data [EC10G_OD]
}

system auto_control = { control with
    directive simulation { sweeps=[sweep_EC10G_dataset0] }
    directive data [EC10G_mRFP1]
}

system auto = { target with
    directive simulation { plots=[[x]*[f530]+b530; [x]*[f480]+b480]; plotcolours=["#FFDF00";
        "#ADD8E6"]; sweeps=[sweep_EC10G_dataset0] }
    directive data [EC10G_EYFP_ECFP]
}

//Constitutive
system prpr_growth = { growth with
    directive simulation { final=36.0; sweeps=[sweep_PRPR_dataset1] }
    directive data [PRPR_OD]
}

system prpr_control = { control with
    directive simulation { final=36.0; sweeps=[sweep_PRPR_dataset1] }
    directive data [PRPR_mRFP1]
}

system prpr = { target with
    directive simulation {final=36.0; sweeps=[sweep_PRPR_dataset1]}
    directive parameters [
        r = 1, { interval=Real; distribution=Uniform(0.1,10); variation=Multiple };
        K = 2, { interval=Real; distribution=Uniform(0.1,5); variation=Multiple };
        tlag = 1, { interval=Real; distribution=Uniform(0,10); variation=Multiple };
        rc=1e2,  { distribution=Uniform(1e0,1e5); variation=Multiple };
        aCFP=1e3,  { distribution=Uniform(1e0,1e5); variation=Random };
        aYFP=1e3,  { distribution=Uniform(1e0,1e5); variation=Random };
        b530=1e3, { interval=Real; distribution=Uniform(0.0,5e3) };
        b480=1e3, { interval=Real; distribution=Uniform(0.0,1e4) };
    ]
    directive data [PRPR_EYFP_ECFP]
    prpr:device
}

// DR (PCat-PCat)
system PCat_growth = { growth with
```

```
    directive simulation {sweeps=[sweep_C6C12_double]}
    directive data [PCat_OD]
}

system PCat_control = { control with
    directive simulation {sweeps=[sweep_C6C12_double]}
    directive data [PCat_mRFP1]
}

system PCat = { target with
    directive simulation {sweeps=[sweep_C6C12_double]}
    directive data [PCat_EYFP_ECFP]
    drPCat:device
}

// DR (R100-S32)
system R100S32_growth = { growth with
    directive simulation {sweeps=[sweep_C6C12]}
    directive data [R100S32_OD]
}

system R100S32_control = { control with
    directive simulation {sweeps=[sweep_C6C12]}
    directive data [R100S32_mRFP1]
}

system R100S32 = { target with
    directive simulation {sweeps=[sweep_C6C12]}
    directive data [R100S32_EYFP_ECFP]
    drR100S32:device
}

// DR (R33-S32)
system R33S32_growth = { growth with
    directive simulation {sweeps=[sweep_C6C12]}
    directive data [R33S32_OD]
}

system R33S32_control = { control with
    directive simulation {sweeps=[sweep_C6C12]}
    directive data [R33S32_mRFP1]
}
system R33S32 = { target with
    directive simulation {sweeps=[sweep_C6C12]}
    directive data [R33S32_EYFP_ECFP]
    drR33S32:device
}

// DR (R33-S175)
system R33S175_growth = { growth with
    directive simulation {sweeps=[sweep_C6C12]}
    directive data [R33S175_OD]
}

system R33S175_control = { control with
    directive simulation {sweeps=[sweep_C6C12]}
    directive data [R33S175_mRFP1]
}
system R33S175 = { target with
```

```
    directive simulation {sweeps=[sweep_C6C12]}
    directive data [R33S175_EYFP_ECFP]
    drR33S175:device
}

// Timed additions circuit (for prediction)
system doublereceiverTimed = { R33S175 with directive parameters [ sigma = 1.0 ] }

// Relays
system R33S175_P76_growth = { growth with
    directive simulation {sweeps=[sweep_relays]}
    directive data [R33S175_P76LasI_OD]
}

system R33S175_P76_control = { control with
    directive simulation {sweeps=[sweep_relays]}
    directive data [R33S175_P76LasI_mRFP1]
}
system R33S175_P76 = { R33S175 with
    directive simulation {sweeps=[sweep_relays]}
    directive data [R33S175_P76LasI_EYFP_ECFP]
    relayP76LasI:device
}

system R33S175_P81_growth = { growth with
    directive simulation {sweeps=[sweep_relays]}
    directive data [R33S175_P81LuxI_OD]
}

system R33S175_P81_control = { control with
    directive simulation {sweeps=[sweep_relays]}
    directive data [R33S175_P81LuxI_mRFP1]
}
system R33S175_P81 = { R33S175 with
    directive simulation {sweeps=[sweep_relays]}
    directive data [R33S175_P81LuxI_EYFP_ECFP]
    relayP81LuxI:device
}

// PBad - Inducer
system PBADYFP = { target with
    directive simulation { plots=[[x]*([yfp]+[f530])+b530]; plotcolours=["#FFDF00"];
        sweeps=[sweep_PBAD_eYFP] }
    directive parameters [
        r = 1, { interval=Real; distribution=Uniform(0.1,10); variation=Multiple };
        K = 2, { interval=Real; distribution=Uniform(0.1,5); variation=Multiple };
        tlag = 1, { interval=Real; distribution=Uniform(0,10); variation=Multiple };
        rc=1e2,  { distribution=Uniform(1e0,1e5); variation=Multiple };
        aYFP=1.0, { distribution=Uniform(1e0,1e5) };
        b530=1e3, { interval=Real; distribution=Uniform(0.0,5e3) };
    ]
    directive data [PBAD_eYFP_EYFP]
    pBADYFP:device
}

system PBADYFP_growth = { growth with
    directive simulation { sweeps=[sweep_PBAD_eYFP] }
    directive data [PBAD_eYFP_OD]
}
```

```
system PBADYFP_control = { control with
    directive simulation { sweeps=[sweep_PBAD_eYFP] }
    directive data [PBAD_eYFP_mRFP1]
}

// AiiA - Degrader
system R33S175_PBADAiiA = { R33S175 with
    directive simulation {sweeps=[sweep_R33S175_PBADAiiA]}
    directive deterministic {reltolerance=1e-12; abstolerance=1e-10}
    directive data [R33S175_PBADAiiA_EYFP_ECFP]
    | pBADAiiA:device
}

// AiiA - Degrader Growth
system R33S175_PBADAiiA_growth = { growth with
    directive simulation {sweeps=[sweep_R33S175_PBADAiiA]}
    directive data [R33S175_PBADAiiA_OD]
}

// AiiA - Degrader Control
system R33S175_PBADAiiA_control = { control with
    directive simulation {sweeps=[sweep_R33S175_PBADAiiA]}
    directive data [R33S175_PBADAiiA_mRFP1]
}

// Auto
node Auto_Growth { systems = [auto_growth] }
node Auto_Control { systems = [auto_control] }
node Auto { systems = [auto]; inference = { burnin=20000; samples=20000; thin=50; partial=false } }
edge Auto_Growth.auto_growth ->[r=Fixed;K=Fixed;tlag=Fixed] Auto_Control.auto_control
edge Auto_Control.auto_control ->[r=Fixed;K=Fixed;tlag=Fixed;rc=Fixed] Auto.auto

// PRPR - Constitutive
node Constitutive_Growth { systems = [prpr_growth] }
node Constitutive_Control { systems = [prpr_control] }
node Constitutive { systems = [prpr]; inference = { partial=false } }
edge Constitutive_Growth.prpr_growth ->[r=Fixed;K=Fixed;tlag=Fixed]
    Constitutive_Control.prpr_control
edge Constitutive_Control.prpr_control ->[r=Fixed;K=Fixed;tlag=Fixed;rc=Fixed] Constitutive.prpr
edge Auto_Control ->[dRFP] Constitutive_Control
edge Auto ->[a530;a480] Constitutive

// Receivers
node Receivers_Growth { systems = [PCat_growth; R100S32_growth; R33S32_growth; R33S175_growth] }
node Receivers_Control { systems = [PCat_control; R100S32_control; R33S32_control;
    R33S175_control] }
node Receivers { systems = [PCat; R100S32; R33S32; R33S175]; inference = { partial=false } }
edge Receivers_Growth.PCat_growth ->[r=Fixed;K=Fixed;tlag=Fixed] Receivers_Control.PCat_control
edge Receivers_Growth.R100S32_growth ->[r=Fixed;K=Fixed;tlag=Fixed]
    Receivers_Control.R100S32_control
edge Receivers_Growth.R33S32_growth ->[r=Fixed;K=Fixed;tlag=Fixed] Receivers_Control.R33S32_control
edge Receivers_Growth.R33S175_growth ->[r=Fixed;K=Fixed;tlag=Fixed]
    Receivers_Control.R33S175_control
edge Receivers_Control.PCat_control ->[r=Fixed;K=Fixed;tlag=Fixed;rc=Fixed] Receivers.PCat
edge Receivers_Control.R100S32_control ->[r=Fixed;K=Fixed;tlag=Fixed;rc=Fixed] Receivers.R100S32
edge Receivers_Control.R33S32_control ->[r=Fixed;K=Fixed;tlag=Fixed;rc=Fixed] Receivers.R33S32
edge Receivers_Control.R33S175_control ->[r=Fixed;K=Fixed;tlag=Fixed;rc=Fixed] Receivers.R33S175
edge Constitutive_Control ->[dRFP] Receivers_Control
```

```
edge Constitutive ->[a530; a480; dYFP; dCFP] Receivers

// Relays
node Relays_Growth { systems = [R33S175_P76_growth; R33S175_P81_growth] }
node Relays_Control { systems = [R33S175_P76_control; R33S175_P81_control] }
node Relays { systems = [R33S175_P76; R33S175_P81]; inference = { partial=false } }
edge Relays_Growth.R33S175_P76_growth ->[r=Fixed;K=Fixed;tlag=Fixed]
    Relays_Control.R33S175_P76_control
edge Relays_Growth.R33S175_P81_growth ->[r=Fixed;K=Fixed;tlag=Fixed]
    Relays_Control.R33S175_P81_control
edge Relays_Control.R33S175_P76_control ->[r=Fixed;K=Fixed;tlag=Fixed;rc=Fixed] Relays.R33S175_P76
edge Relays_Control.R33S175_P81_control ->[r=Fixed;K=Fixed;tlag=Fixed;rc=Fixed] Relays.R33S175_P81
edge Receivers_Control ->[dRFP] Relays_Control
edge Receivers ->[
   a530;a480;dYFP;dCFP;
   aR33;aS175;aYFP;aCFP;e76;e81;KGR_76;KGS_76;KGR_81;KGS_81;KR6;KS6;KR12;KS12;nR;nS;dR;
] Relays

// PBad - Inducer
node Inducer_Growth { systems = [PBADYFP_growth] }
node Inducer_Control { systems = [PBADYFP_control] }
node Inducer { systems = [PBADYFP]; inference = { partial=false } }
edge Inducer_Growth.PBADYFP_growth ->[r=Fixed;K=Fixed;tlag=Fixed] Inducer_Control.PBADYFP_control
edge Inducer_Control.PBADYFP_control ->[r=Fixed;K=Fixed;tlag=Fixed;rc=Fixed] Inducer.PBADYFP
edge Constitutive_Control ->[dRFP] Inducer_Control
edge Constitutive ->[a530;dYFP] Inducer

// AiiA - Degrader
node Degrader_Growth { systems = [R33S175_PBADAiiA_growth] }
node Degrader_Control { systems = [R33S175_PBADAiiA_control] }
node Degrader { systems = [R33S175_PBADAiiA]; inference = { partial=false } }
edge Degrader_Growth.R33S175_PBADAiiA_growth ->[r=Fixed;K=Fixed;tlag=Fixed]
    Degrader_Control.R33S175_PBADAiiA_control
edge Degrader_Control.R33S175_PBADAiiA_control ->[r=Fixed;K=Fixed;tlag=Fixed;rc=Fixed]
    Degrader.R33S175_PBADAiiA
edge Receivers_Control ->[dRFP] Degrader_Control
edge Receivers ->[
   a530;a480;dYFP;dCFP;
   aR33;aS175;aYFP;aCFP;e76;e81;KGR_76;KGS_76;KGR_81;KGS_81;KR6;KS6;KR12;KS12;nR;nS;dR;
] Degrader
edge Inducer ->[KAra; nA; eA] Degrader
```

### S2 Models of cells expressing synthetic gene circuits

This section explains the models of the cell lines that were built and tested for the Constant hypothesis. Each subsection focuses on a system specified in the code in Listing 1. For the sake of convenience and readability, we have unrolled (expanded) the systems to be a self-contained system definition. Tables S2 and S3 list the parts and devices that were built, tested, and used in this case study.

#### S2.1 No plasmid (Autofluorescence)

The simplest possible cell line to characterize is one in which there is no synthetic gene circuit at all. However, applying dynamic characterization to no circuit is useful to characterize how autofluorescence depends on cell density and gene expression capacity. Therefore, our chromosomal RFP-expressing cells were measured under a range of conditions, to explore how gene expression capacity influenced time-series measurements at fluorescence wavelengths corresponding to eYFP and eCFP .

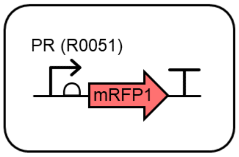

**Figure S8: Design of chromosomal RFP.**

##### Model definition

The model we used for autofluorescence assumes that the rate of autofluorescence is proportional to gene expression capacity, $h(c)$, and that the fluorescent material dilutes with cell growth. Bulk autofluorescence is then quantified in exactly the same way as it is when there is a synthetic gene circuit inside the cells, using equation (4) of the main text. As such, the equations for intracellular autofluorescence corresponding to eYFP and eCFP are

$$\frac{dc}{dt} = \gamma(c).c \tag{S1a}$$
$$\frac{d[F_{530}]}{dt} = a_{530}.h(c) - \gamma(c)[F_{530}] \tag{S1b}$$
$$\frac{d[F_{480}]}{dt} = a_{480}.h(c) - \gamma(c)[F_{480}] \tag{S1c}$$

To compare with experimental measurements, we consider the bulk fluorescence given by

$$B_{480} = c.[F_{480}] + b_{480} \tag{S2a}$$
$$B_{530} = c.[F_{530}] + b_{530} \tag{S2b}$$

where $[F_{480}]$ and $[F_{530}]$ are modeled as in (14). This is equivalent to the observer process described in the Methods section, but with [YFP] and [CFP] set to zero, as they are not expressed in this circuit.

##### Implementation with GEC/CRN language

To implement the combination of modeling cell density and autofluorescence in the GEC language, we define a module that can be called as part of the implementation of any other circuit. We also use *rate* expressions to define $\gamma$ (`growth`) and $h(c)$ (`capacity`, here the TargetSwitch hypothesis), as these are reused in multiple reactions across different modules/systems.

```
directive rates [
   growth = [grow]*r*(1 - [x] / K);
```

```
    capacity = rc;
]
module cells(growth,tlag) = {
    init x c0 | init grow 1 @ tlag |
    init c6 C6 @ tau | init c12 C12 @ tau |
    ->[[growth]*[x]] x
}
module autofluorescence(growth,capacity) = {
    ->[[capacity]*a530] f530 |
    f530 ->[[growth]*[f530]] |
    ->[[capacity]*a480] f480 |
    f480 ->[[growth]*[f480]]
}
```

##### Characterization experiment

We infer the Auto model parameters by using uniform priors on $a_{530}$, $a_{480}$, $b_{530}$, $b_{480}$ and any parameters that are specific to the gene expression capacity hypothesis being considered. The parameters within $\gamma$ are taken to be the maximum likelihood estimates from the cell density characterization phase, and the $\xi$ are either taken to be the maximum likelihood estimates from the control characterization phase or are inferred within the target phase, depending on the hypothesis in question.

According, the overall model for the Auto circuit is simply that instantiation of the `cells` module above, along with the specification of the parameters to be inferred:

```
system auto = {
    directive simulation { final=36.0; points=250; plots=[[x]*[f530]+b530; [x]*[f480]+b480];
        plotcolours=["#FFDF00"; "#ADD8E6"] }
    directive parameters [
        r = 1, { interval=Real; distribution=Uniform(0.1,10); variation=Multiple };
        K = 2, { interval=Real; distribution=Uniform(0.1,5); variation=Multiple };
        tlag = 1, { interval=Real; distribution=Uniform(0,10); variation=Multiple };
        rc=1e2,  { interval=Log; distribution=Uniform(1e0,1e5); variation=Multiple };
        b530=1e3, { interval=Real; distribution=Uniform(0.0,5e3); variation=Random };
        b480=1e3, { interval=Real; distribution=Uniform(0.0,1e4); variation=Random };
    ]
    directive rates [growth = [grow]*r*(1 - [x] / K);capacity = rc]
    directive crn { cells(growth,tlag) | autofluorescence(growth,capacity) }
}
```

#### Characterization results

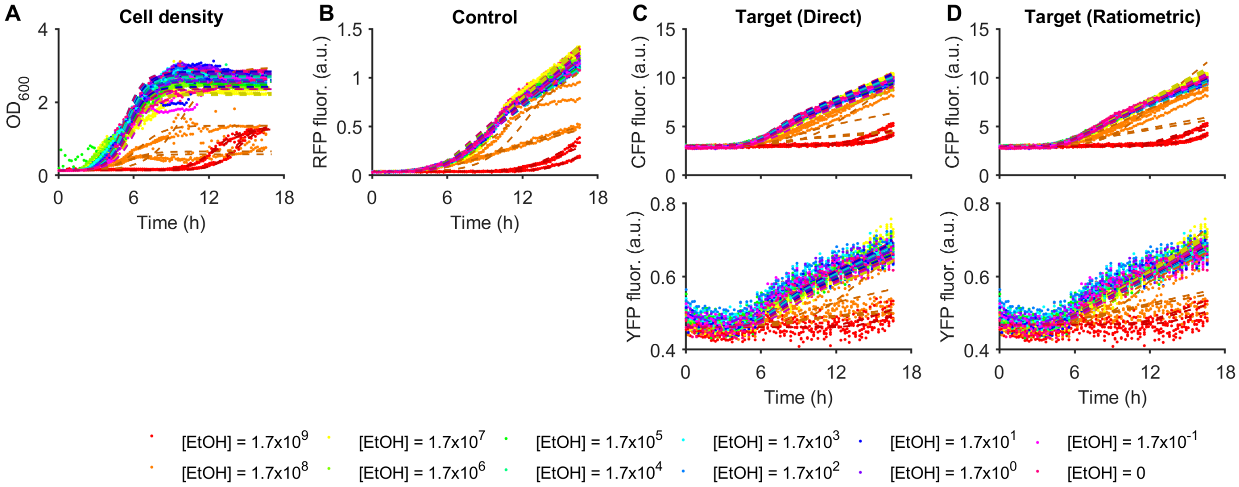

**Figure S9: Model-data comparison for the Auto circuit with TargetSwitch hypothesis.** Measurements of the Auto circuit are compared against simulations of the maximum likelihood parameter set from the TargetSwitch hypothesis.

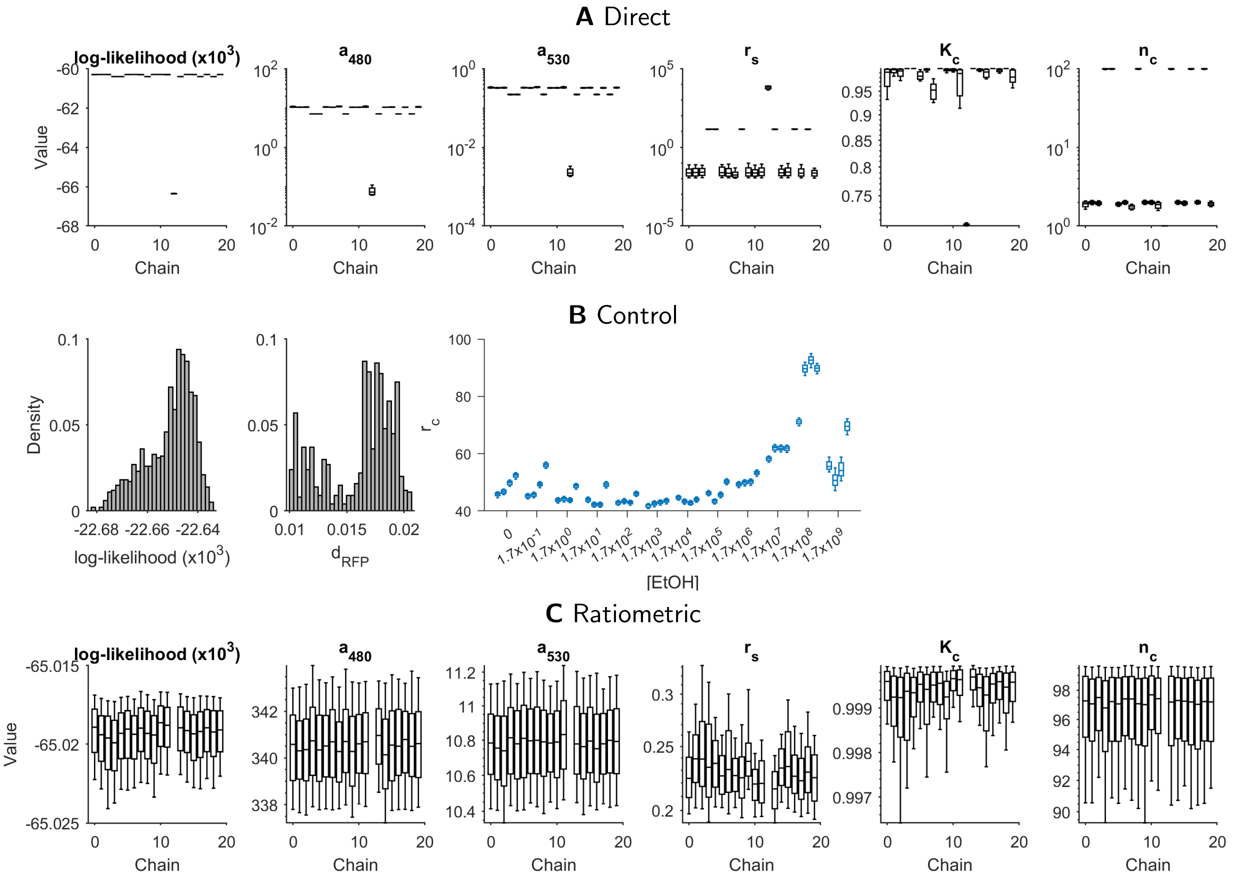

**Figure S10: Marginal parameter posterior distributions for Auto with TargetSwitch hypothesis. A.** Inference of circuit parameters using the direct (non-ratiometric) method, which corresponds to assuming that $r_c = 1$ for all time-series traces. **B.** The control phase identifies values for $r_c$ for each time-series, to be used in ratiometric characterization. **C.** Inference of circuit parameters using the ratiometric method, using the values of $r_c$ in B.

#### S2.2 Constitutive expression via the PR promoter

The simplest circuit that we consider in this article uses constitutive promoters to drive expression of eYFP and eCFP . Using such a simple circuit enables us to characterize properties of the fluorescent proteins, which can then be reused to aide characterization of circuits using the same fluorescent proteins.

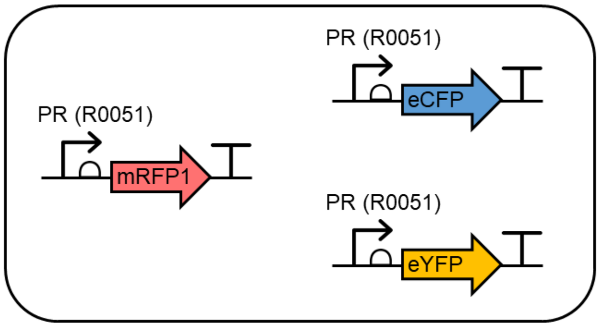

**Figure S11: Design of constitutive expression circuit.**

##### Model definition

As this is the first (non-trivial) synthetic gene circuit that we describe, we include a detailed derivation. We start by considering the chemical reactions introduced by the circuit in Fig. S11.

If we denote by $g$ the plasmid containing the PL:eYFP and PL:eCFP cassettes, then we can write a system of chemical reactions that describe transcription, translation and fluorescent protein maturation as

$$g \xrightarrow{a_{r0011}} g + \text{m\_b0034\_eYFP} \tag{S3a}$$

$$\text{m\_b0034\_eYFP} \xrightarrow{d_{\text{m\_b0034\_eYFP}}} \varnothing \tag{S3b}$$

$$\text{m\_b0034\_eYFP} \xrightarrow{k_{b0034} \cdot h(c)} \text{m\_b0034\_eYFP} + \text{eYFPi} \tag{S3c}$$

$$\text{eYFPi} \xrightarrow{d_{\text{YFP}}} \varnothing \tag{S3d}$$

$$\text{eYFPi} \xrightarrow{\mu_{\text{YFP}}} \text{eYFP} \tag{S3e}$$

$$\text{eYFP} \xrightarrow{d_{\text{YFP}}} \varnothing \tag{S3f}$$

$$g \xrightarrow{a_{r0011}} g + \text{m\_b0034\_eCFP} \tag{S3g}$$

$$\text{m\_b0034\_eCFP} \xrightarrow{d_{\text{m\_b0034\_eCFP}}} \varnothing \tag{S3h}$$

$$\text{m\_b0034\_eCFP} \xrightarrow{k_{b0034} \cdot h(c)} \text{m\_b0034\_eCFP} + \text{eCFPi} \tag{S3i}$$

$$\text{eCFPi} \xrightarrow{d_{\text{CFP}}} \varnothing \tag{S3j}$$

$$\text{eCFPi} \xrightarrow{\mu_{\text{CFP}}} \text{eCFP} \tag{S3k}$$

$$\text{eCFP} \xrightarrow{d_{\text{CFP}}} \varnothing \tag{S3l}$$

where $h(c)$ is the gene expression capacity, as introduced in the Methods section of the main text.

Translating to ODEs, and replacing $[g]$ with $N$, the plasmid copy number, we obtain

$$\frac{dc}{dt} = \gamma(c).c \tag{S4a}$$

$$\frac{d[\text{m\_b0034\_eYFP}]}{dt} = a_{r0011}.N - (d_{\text{m\_b0034\_eYFP}} + \gamma(c))[\text{m\_b0034\_eYFP}] \tag{S4b}$$

$$\frac{d[\text{eYFPi}]}{dt} = k_{b0034}.h(c).[\text{m\_b0034\_eYFP}] - (d_{\text{YFP}} + \mu_{\text{YFP}} + \gamma(c))[\text{eYFP}] \tag{S4c}$$

$$\frac{d[\text{eYFP}]}{dt} = \mu_{\text{YFP}}[\text{eYFPi}] - (d_{\text{YFP}} + \gamma(c))[\text{eYFP}] \tag{S4d}$$

$$\frac{d[\text{m\_b0034\_eCFP}]}{dt} = a_{r0011}.N - d_{\text{m\_b0034\_eCFP}}[\text{m\_b0034\_eCFP}] \tag{S4e}$$

$$\frac{d[\text{eCFPi}]}{dt} = k_{b0034}.h(c).[\text{m\_b0034\_eCFP}] - (d_{\text{CFP}} + \mu_{\text{CFP}} + \gamma(c))[\text{eCFPi}] \tag{S4f}$$

$$\frac{d[\text{eCFP}]}{dt} = \mu_{\text{CFP}}[\text{eCFPi}] - (d_{\text{CFP}} + \gamma(c))[\text{eCFP}] \tag{S4g}$$

where $\gamma(c)$ is the cellular dilution of each molecular concentration.

**Assumption 1:** By assuming that mRNA dynamics are fast, we can remove the mRNA species from the model entirely. That is, we equate (S4b) and (S4e) to zero, solve for [m_b0034_eYFP] and [m_b0034_eCFP], and then substitute into the remaining equations. This results in the reduced model

$$\frac{dc}{dt} = \gamma(c).c \tag{S5a}$$

$$\frac{d[\text{eYFPi}]}{dt} = a_{\text{YFP}}.h(c) - (d_{\text{YFP}} + \mu_{\text{YFP}} + \gamma(c))[\text{eYFP}] \tag{S5b}$$

$$\frac{d[\text{eYFP}]}{dt} = \mu_{\text{YFP}}[\text{eYFPi}] - (d_{\text{YFP}} + \gamma(c))[\text{eYFP}] \tag{S5c}$$

$$\frac{d[\text{eCFPi}]}{dt} = a_{\text{CFP}}.h(c) - (d_{\text{CFP}} + \mu_{\text{CFP}} + \gamma(c))[\text{eCFPi}] \tag{S5d}$$

$$\frac{d[\text{eCFP}]}{dt} = \mu_{\text{CFP}}[\text{eCFPi}] - (d_{\text{CFP}} + \gamma(c))[\text{eCFP}] \tag{S5e}$$

where

$$a_{\text{YFP}} = \frac{a_{r0011}.N.k_{b0034}}{d_{\text{m\_b0034\_eYFP}} + \gamma(c)} \approx \frac{a_{r0011}.N.k_{b0034}}{d_{\text{m\_b0034\_eYFP}}}, \qquad a_{\text{CFP}} = \frac{a_{r0011}.N.k_{b0034}}{d_{\text{m\_b0034\_eCFP}} + \gamma(c)} \approx \frac{a_{r0011}.N.k_{b0034}}{d_{\text{m\_b0034\_eCFP}}} \tag{S6}$$

Here, we remove $\gamma(c)$ from the denominator because dilution will normally be slower than mRNA degradation, and making this simplification results in $a_{\text{YFP}}$ and $a_{\text{CFP}}$ being constant. Finally, we note here that despite simplifying the parametrization (removing $a_{r0011}$, $k_{b0034}$, $d_{\text{m\_b0034\_eYFP}}$ and $d_{\text{m\_b0034\_eCFP}}$), we should keep in mind that there is a dependency of $a_{\text{YFP}}$ and $a_{\text{CFP}}$ on the biological parts in each cassette.

**Assumption 2:** By assuming that fluorescent protein maturation is fast, we can remove the equations for the concentration of immature fluorescent proteins. Let [YFP] be the sum of the concentrations of immature and mature fluorescent proteins. Adding (S5b) to (S5c), we obtain

$$\frac{d}{dt}([\text{eYFPi}] + [\text{eYFP}]) = a_{\text{YFP}}.h(c) - (d_{\text{YFP}} + \gamma(c))([\text{eYFPi}] + [\text{eYFP}])$$

$$\implies \frac{d[\text{YFP}]}{dt} = a_{\text{YFP}}.h(c) - (d_{\text{YFP}} + \gamma(c))[\text{YFP}] \tag{S7}$$

and similarly for eCFP . If we assume that the immature form is instantly converted to the mature form, then the concentration of the mature form is equal to the total concentration. Therefore, the resultant system of equations is given by

$$\frac{dc}{dt} = \gamma(c).c \tag{S8a}$$

$$\frac{d[\text{YFP}]}{dt} = a_{\text{YFP}}.h(c) - (d_{\text{YFP}} + \gamma(c))[\text{YFP}] \tag{S8b}$$

$$\frac{d[\text{CFP}]}{dt} = a_{\text{CFP}}.h(c) - (d_{\text{CFP}} + \gamma(c))[\text{CFP}] \tag{S8c}$$

We compare with experimental measurements using the observer process of (4).

##### Implementation with GEC/CRN language

To implement the PRPR circuit in the GEC language, we define modules for the eYFP and eCFP proteins, which can then be called by multiple systems (note that module definitions are identical in both GEC and CRN). We require an argument `P` for the normalized promoter activity (rate expression) and another argument `a` for the maximal production rate. This enables circuits that use different promoters or ribosome binding sites for eYFP and eCFP to provide different contexts in the module call. The rate expressions `growth` and `capacity` are used equivalently to the Auto circuit.

```
module CFP(P,a,growth,capacity) = {
    ->[[capacity]*a*[P]] cfp |
    cfp ->{dCFP} |
    cfp ->[[growth]*[cfp]]
}
module YFP(P,a,growth,capacity) = {
    ->[[capacity]*a*[P]] yfp |
    yfp ->{dYFP} |
    yfp ->[[growth]*[yfp]]
}
device prpr() = { YFP(one,aYFP,growth,capacity) | CFP(one,aCFP,growth,capacity) }
```

It's also important to note that this code has incorporated the assumptions above concerning fluorescent protein maturation and mRNA expression. Alternative assumptions can be readily explored in this framework by modifying these module definitions.

##### Characterization experiment

We measured the PRPR circuit in response to varying concentrations of chloramphenicol, an inhibitor of protein translation. By inhibiting protein translation, we directly alter the gene expression capacity term in our models, enabling us to test whether $h(c)$ can capture such an effect implicitly. Unfortunately, chloramphenicol also considerably alters cell growth, so separating its effects on cell growth and gene expression is a challenge for the methods we are demonstrating in this paper.

We infer the PRPR model parameters by using (uninformative) uniform priors on the previously uncharacterized parameters $a_{\text{CFP}}$, $a_{\text{YFP}}$, $d_{\text{CFP}}$ and $d_{\text{YFP}}$, and (informative) truncated Gaussian priors on $a_{480}$ and $a_{530}$, with mean and standard deviation taken from the marginal posteriors of the Auto circuit characterization. These parameters are globally specified, with propagation of marginal posteriors as edges of the inference graph (see Listing 1). We also use uninformative priors on the background fluorescence parameters (specified below as local parameters `b480` and `b530`), without propagating their marginal posteriors from previous circuits.

In the GEC language, the overall model for the PRPR circuit combines instantiation of the `cells` and autofluorescence modules and the prpr device. In the following system definition, a device invocation `prpr` implies that the system must use the device model definition `device prpr()`. The structural definition of the `prpr` device (specified in the GEC library and detailed in Table S3) is : pr; rbs34; eyfp; ter1; pr; rbs34; ecfp; ter1. The part definitions for these DNA components can be found in Table S2. This applies for all device invocations in systems in all subsequent sections.

```
system prpr = {
   directive simulation { final=36.0; points=250; plots=[[x]*([yfp]+[f530])+b530;
       [x]*([cfp]+[f480])+b480]; plotcolours=["#FFDF00"; "#ADD8E6"] }
   directive rates [growth = [grow]*r*(1 - [x] / K);capacity = rc]
   directive crn { cells(growth,tlag) | autofluorescence(growth,capacity) }
   directive parameters [
      r = 1, { interval=Real; distribution=Uniform(0.1,10); variation=Multiple };
      K = 2, { interval=Real; distribution=Uniform(0.1,5); variation=Multiple };
      tlag = 1, { interval=Real; distribution=Uniform(0,10); variation=Multiple };
      rc=1e2,  { interval=Log; distribution=Uniform(1e0,1e5); variation=Multiple };
      aCFP=1e3,  { interval=Log; distribution=Uniform(1e0,1e5); variation=Random };
      aYFP=1e3,  { interval=Log; distribution=Uniform(1e0,1e5); variation=Random };
      b530=1e3, { interval=Real; distribution=Uniform(0.0,5e3); variation=Random };
      b480=1e3, { interval=Real; distribution=Uniform(0.0,1e4); variation=Random };
   ]
   prpr:device
}
```

Here, system-local parameters aCFP and aYFP replace the global parameters (see Listing 1). This is to enable the PR promoter in combination with the b0034 rbs to take on a different maximum production rate than the PLux-driven production rates of the other devices. The other important parameters for this device are dCFP and dYFP, and are globally specified (Listing 1), since they are shared between devices.

##### Characterization results

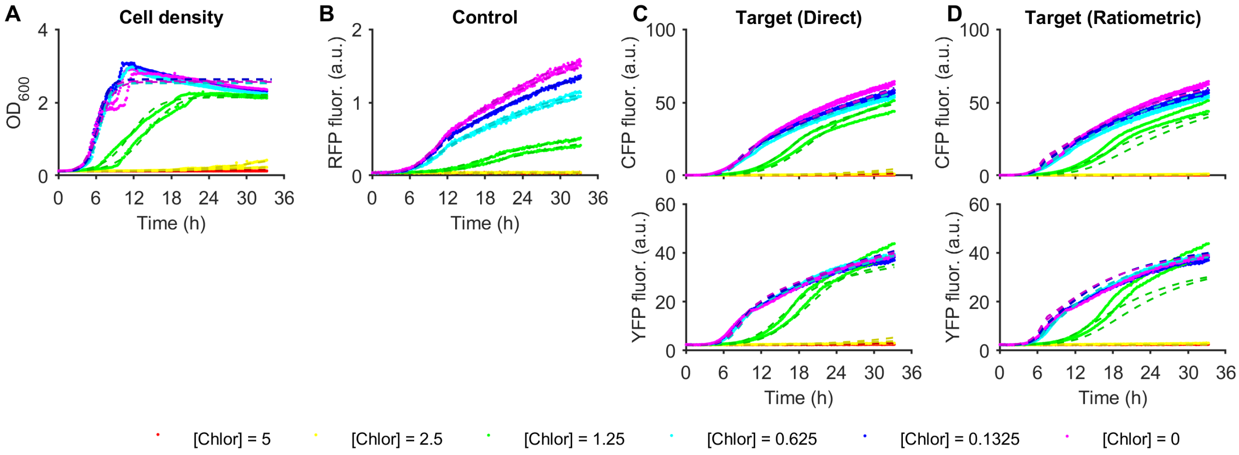

**Figure S12: Model-data comparison for the PRPR circuit with TargetSwitch hypothesis.** Measurements of the PRPR circuit are compared against simulations of the maximum likelihood parameter set from the TargetSwitch hypothesis.

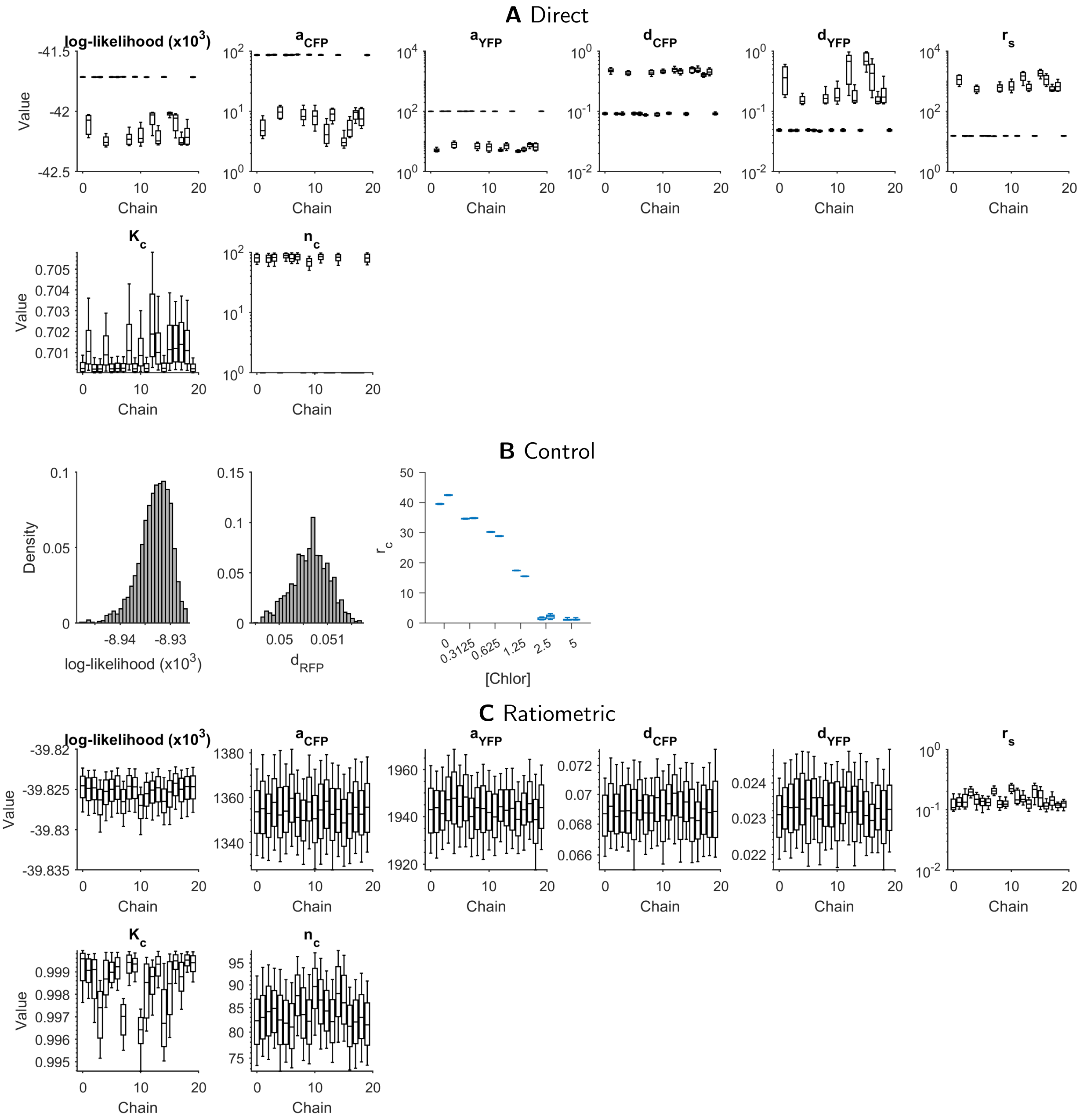

**Figure S13: Marginal parameter posterior distributions for PRPR with TargetSwitch hypothesis. A.** Inference of circuit parameters using the direct (non-ratiometric) method, which corresponds to assuming that $r_c = 1$ for all time-series traces. **B.** The control phase identifies values for $r_c$ for each time-series, to be used in ratiometric characterization. **C.** Inference of circuit parameters using the ratiometric method, using the values of $r_c$ in B.

#### S2.3 HSL Double Receivers

We consider the dynamic characterization of the AHL double receiver (DR) circuit introduced in [3]. In this device, two variations of the wild-type PLux promoter, PLux76 and PLas81, were engineered to bind preferentially to activated luxR and lasR complexes respectively. As LuxR favours binding of C6 and LasR favours binding of C12, optimized expression of LuxR and LasR can lead to near-orthogonal intracellular detection of C6 and C12. The double receiver device was originally measured with PLux76 upstream of the coding sequence for eCFP , and PLas81 upstream of the coding sequence for eYFP . A plasmid containing all of this machinery was then inserted into cells chromosomally expressing mRFP1, which could be used as a ratiometric control.

##### Model definition

As the modelling philosophy we are considering here describes dynamics of proteins, the model of DR circuits in this paper differs from model described previously [3], as the concentrations of luxR and lasR become dynamic quantities affected by growth dilution. Furthermore, in the previous paper, we parametrized the double receiver by first parametrizing circuits that only have one PLux promoter. Here, we seek to parametrize the DR circuits directly, by inferring parameters against eYFP and eCFP measurements simultaneously. As such, the model must consider that luxR and lasR-based regulators (bound to HSLs) might bind to more than one (PLux) promoter.

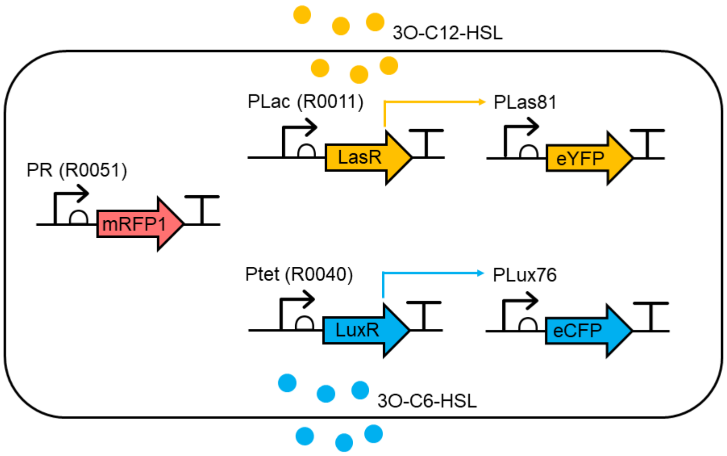

**Figure S14: Design of a DR circuit.**

We denote by $C_k$ the HSL molecule with length $k$ carbon chain, and by $G_i$ the PLux76 and PLas81 promoters. Then similar to the derivation in [3], we can specify all of the reactions between the HSLs, LuxR and LasR, and eventual binding of transcriptional regulators to PLux76/PLas81.

$$\begin{aligned}
\text{luxR} + C_k &\underset{u_{Rk}}{\overset{b_{Rk}}{\rightleftharpoons}} \text{luxR-}C_k \\
\text{lasR} + C_k &\underset{u_{Sk}}{\overset{b_{Sk}}{\rightleftharpoons}} \text{lasR-}C_k \\
\text{luxR-}C_k + \text{luxR-}C_k &\underset{u_{Dk}}{\overset{b_{Dk}}{\rightleftharpoons}} \text{luxR-}C_k\text{-luxR-}C_k \\
\text{lasR-}C_k + \text{lasR-}C_k &\underset{u_{Ek}}{\overset{b_{Ek}}{\rightleftharpoons}} \text{lasR-}C_k\text{-lasR-}C_k \\
G_i + \text{luxR-}C_k\text{-luxR-}C_k &\underset{u_{GiDk}}{\overset{b_{GiDk}}{\rightleftharpoons}} G_i\text{-luxR-}C_k\text{-luxR-}C_k \\
G_i + \text{lasR-}C_k\text{-lasR-}C_k &\underset{u_{GiEk}}{\overset{b_{GiEk}}{\rightleftharpoons}} G_i\text{-lasR-}C_k\text{-lasR-}C_k
\end{aligned} \tag{S9}$$

Constitutive expression of luxR, lasR is described by

$$\varnothing \xrightarrow{a_{\text{luxR}}} \text{luxR} \qquad \varnothing \xrightarrow{a_{\text{lasR}}} \text{lasR} \tag{S10}$$

Inducible expression of eCFP and eYFP by $\text{P}_{\text{OLux}}$ and $\text{P}_{\text{OLas}}$ respectively is described by

$$\begin{aligned}
G_{\text{pLux76}} &\xrightarrow{a0_{\text{pLux76}}} G_{\text{pLux76}} + \text{m_b0034_eCFP} \\
G_{\text{pLux76}}\text{-luxR-}C_k\text{-luxR-}C_k &\xrightarrow{a1_{\text{pLux76}}} G_{\text{pLux76}}\text{-luxR-}C_k\text{-luxR-}C_k + \text{m_b0034_eCFP} \\
G_{\text{pLux76}}\text{-lasR-}C_k\text{-lasR-}C_k &\xrightarrow{a1_{\text{pLux76}}} G_{\text{pLux76}}\text{-lasR-}C_k\text{-lasR-}C_k + \text{m_b0034_eCFP} \\
G_{\text{pLas81}} &\xrightarrow{a0_{\text{pLas81}}} G_{\text{pLas81}} + \text{m_b0034_eCFP} \\
G_{\text{pLas81}}\text{-luxR-}C_k\text{-luxR-}C_k &\xrightarrow{a0_{\text{pLas81}}} G_{\text{pLas81}}\text{-luxR-}C_k\text{-luxR-}C_k + \text{m_b0034_eYFP} \\
G_{\text{pLas81}}\text{-lasR-}C_k\text{-lasR-}C_k &\xrightarrow{a1_{\text{pLas81}}} G_{\text{pLas81}}\text{-lasR-}C_k\text{-lasR-}C_k + \text{m_b0034_eCFP}
\end{aligned} \tag{S11}$$

The full DR circuit comprises reactions (S9), (S10) and (S11), and the same reactions as shown for the PRPR circuit that describe mRNA translation for eCFP and eYFP .

To produce a simplified ODE model amenable to parameter inference, we start with the equations describing luxR and lasR protein, their complexes involving $C_6$ and $C_{12}$, and the bound/unbound promoters. Applying also Assumption 1 from before, we arrive a set of equations for the non-mRNA species as

$$\frac{dc}{dt} = \gamma(c).c \tag{S12a}$$

$$\frac{d[\text{luxR}]}{dt} = a_R + u_{Rk}[\text{luxR-}C_k] - [\text{luxR}](\gamma(c) + d_R + b_{Rk}[C_k]) \tag{S12b}$$

$$\frac{d[\text{lasR}]}{dt} = a_S + u_{Sk}[\text{lasR-}C_k] - [\text{lasR}](\gamma(c) + d_S + b_{Sk}[C_k]) \tag{S12c}$$

$$\frac{d[\text{luxR-}C_k]}{dt} = b_{Rk}[\text{luxR}][C_k] + 2u_{Dk}[\text{luxR-}C_k\text{-luxR-}C_k] - [\text{luxR-}C_k](\gamma(c) + d_R^1 + u_{Rk} + 2b_{Dk}[\text{luxR-}C_k]) \tag{S12d}$$

$$\frac{d[\text{lasR-}C_k]}{dt} = b_{Sk}[\text{lasR}][C_k] + 2u_{Ek}[\text{lasR-}C_k\text{-lasR-}C_k] - [\text{lasR-}C_k](\gamma(c) + d_S^1 + u_{Sk} + 2b_{Ek}[\text{lasR-}C_k]) \tag{S12e}$$

$$\begin{aligned}\frac{d[\text{luxR-}C_k\text{-luxR-}C_k]}{dt} &= b_{Dk}[\text{luxR-}C_k]^2 + u_{GiDk}\sum_i[G_i\text{-luxR-}C_k\text{-luxR-}C_k]\ldots \\ &\quad - [\text{luxR-}C_k\text{-luxR-}C_k]\left(\gamma(c) + d_R^1 + u_{Dk} + b_{GiDk}\sum_i[G_i]\right)\end{aligned} \tag{S12f}$$

$$\begin{aligned}\frac{d[\text{lasR-}C_k\text{-lasR-}C_k]}{dt} &= b_{Ek}[\text{lasR-}C_k]^2 + u_{GiEk}\sum_i[G_i\text{-lasR-}C_k\text{-lasR-}C_k]\ldots \\ &\quad - [\text{lasR-}C_k\text{-lasR-}C_k]\left(\gamma(c) + d_R^1 + u_{Ek} + b_{GiEk}\sum_i[G_i]\right)\end{aligned} \tag{S12g}$$

$$\frac{d[G_i\text{-luxR-}C_k\text{-luxR-}C_k]}{dt} = b_{GiDk}[G_i][\text{luxR-}C_k\text{-luxR-}C_k] - (\gamma(c) + u_{GiDk})[G_i\text{-luxR-}C_k\text{-luxR-}C_k] \tag{S12h}$$

$$\frac{d[G_i\text{-lasR-}C_k\text{-lasR-}C_k]}{dt} = b_{GiEk}[G_i][\text{lasR-}C_k\text{-lasR-}C_k] - (\gamma(c) + u_{GiEk})[G_i\text{-lasR-}C_k\text{-lasR-}C_k] \tag{S12i}$$

$$\begin{aligned}\frac{d[G_i]}{dt} &= \gamma(c).n_G + u_{GiDk}[G_i\text{-luxR-}C_k\text{-luxR-}C_k] + u_{GiEk}[G_i\text{-lasR-}C_k\text{-lasR-}C_k]\ldots \\ &\quad - [G_i](\gamma(c) + b_{GiDk}[\text{luxR-}C_k\text{-luxR-}C_k] + b_{GiEk}[\text{lasR-}C_k\text{-lasR-}C_k])\end{aligned} \tag{S12j}$$

Here, luxR-$C_k$ refers to a luxR-HSL heterodimer and luxR-$C_k$-luxR-$C_k$ is the tetramer comprising two luxR-$C_k$ complexes (no cross-binding of C6 and C12 dimers). Similarly, lasR-$C_k$ and lasR-$C_k$-lasR-$C_k$ are equivalent lasR-HSL complexes. The luxR and lasR proteins are assumed to be stabilized by the binding of signal, as in [4]. Therefore, rates $d_j^1 < d_j$ ($j = R, S$) are the slower rates for signal-bound luxR/lasR proteins.

Solving most of the above system equal to zero, we obtain the quasi-equilibrium

$$[G_i\text{-luxR-}C_k\text{-luxR-}C_k]^* = K_{GDk}[G_i][\text{luxR-}C_k\text{-luxR-}C_k] \tag{S13a}$$

$$[\text{luxR-}C_k\text{-luxR-}C_k]^* = K_{Dk}[\text{luxR-}C_k]^2 \tag{S13b}$$

$$[\text{luxR-}C_k]^* = K_{Rk}[\text{luxR}][C_k] \tag{S13c}$$

where $K_{Rk} = \dfrac{b_{Rk}}{\gamma + u_{Rk}}$, $K_{Dk} = \dfrac{b_{Dk}}{u_{Dk}}$ and $K_{GiDk} = \dfrac{b_{GiDk}}{u_{GiDk}}$. Therefore (also symmetry of luxR and lasR),

$$[G_i\text{-luxR-}C_k\text{-luxR-}C_k]^* = K_{GiDk}K_{Dk}\left(K_{Rk}[C_k][\text{luxR}]\right)^2 \tag{S14a}$$

$$[G_i\text{-lasR-}C_k\text{-lasR-}C_k]^* = K_{GiEk}K_{Ek}\left(K_{Sk}[C_k][\text{lasR}]\right)^2 \tag{S14b}$$

where the new $K$'s are defined as above.

In the reduced system, total luxR is described by the equations

$$\frac{d[\text{luxR}]_T}{dt} = \frac{d[\text{luxR}]}{dt} + \sum_k \left( \frac{d[\text{luxR-}C_k]}{dt} + 2\frac{d[\text{luxR-}C_k\text{-luxR-}C_k]}{dt} + 2\sum_i \frac{d[G_i\text{-luxR-}C_k\text{-luxR-}C_k]}{dt} \right) \tag{S15a}$$

$$= a_R - \gamma[\text{luxR}] - d_R[\text{luxR}]\dots$$

$$- \sum_k \left( (\gamma + d_R^1)[\text{luxR-}C_k] + (\gamma + d_R^1)[\text{luxR-}C_k\text{-luxR-}C_k] + \gamma \sum_i [G_i\text{-luxR-}C_k\text{-luxR-}C_k] \right) \tag{S15b}$$

$$= a_R - \gamma[\text{luxR}]_T - d_R^1[\text{luxR}]_T - (d_R - d_R^1)[\text{luxR}] \tag{S15c}$$

The final term needs careful attention, as we expect a difference between $d_R$ and $d_R^1$. Nevertheless, the obvious approximation is to ignore this difference, and model $[\text{luxR}]_T$ explicitly with no dependence on $[R]$. We can write down expressions for the fraction of $[\text{luxR}]_T$ that is bound to signal, dimerized, etc., using the equilibrium relationships above. E.g.

$$[\text{luxR}]_T = [\text{luxR}] + \sum_k \left( [\text{luxR-}C_k] + 2[\text{luxR-}C_k\text{-luxR-}C_k] + 2\sum_i [G_i\text{-luxR-}C_k\text{-luxR-}C_k] \right)$$

$$= [\text{luxR}] + \sum_k \left( K_{Rk}[\text{luxR}][C_k] + 2K_{Dk}K_{Rk}^2[\text{luxR}]^2[C_k]^2 + \sum_i 2K_{GiDk}[G_i]K_{Dk}K_{Rk}^2[\text{luxR}]^2[C_k]^2 \right)$$

When $C_k$ is low, total LuxR is closely approximated by free LuxR, $[\text{luxR}]_T \approx [\text{luxR}]$. But when $C_k$ is high, $[\text{luxR}]_T$ should be partitioned between the $[\text{luxR-}C_k\text{-luxR-}C_k]$ and $[G_i\text{-luxR-}C_k\text{-luxR-}C_k]$ species. Therefore, to simplify the analysis, we propose modelling this by using the assumption

$$[\text{luxR}]_T \approx [\text{luxR}] + \sum_k [\text{luxR-}C_k] = [\text{luxR}] \left( 1 + \sum_k K_{Rk}[C_k] \right) \tag{S16}$$

which still captures the saturation of luxR by $C_k$, using the approximations

$$[\text{luxR}] \approx [\text{luxR}]_T.\frac{1}{1 + \sum_k K_{Rk}[C_k]} \tag{S17a}$$

$$[\text{luxR-}C_k] \approx [\text{luxR}]_T.\frac{K_{Rk}[C_k]}{1 + \sum_k K_{Rk}[C_k]} \tag{S17b}$$

$$[\text{luxR-}C_k\text{-luxR-}C_k] \approx K_{Dk}[\text{luxR}]_T^2 \left( \frac{K_{Rk}[C_k]}{1 + \sum_k K_{Rk}[C_k]} \right)^2 \tag{S17c}$$

$$[G_i\text{-luxR-}C_k\text{-luxR-}C_k] \approx K_{GR}^{(i)}[G_i][\text{luxR}]_T^2 \left( \frac{K_{Rk}[C_k]}{1 + \sum_k K_{Rk}[C_k]} \right)^2 \tag{S17d}$$

where $K_{GR}^{(i)} = K_{GiDk}K_{Dk}$ is assumed to be independent of which signal is bound ($k$), equivalent to the derivation in [3].

The approximation also allows for saturation of $G_i$. By taking advantage of the conservation law $[G_i] + [G_i\text{.luxR-}C_6\text{-luxR-}C_6] + [G_{12}\text{-luxR-}C_]\text{-luxR-}C_]] + [G_6\text{-lasR-}C_]\text{-lasR-}C_]] + [G_{12}\text{-lasR-}C_]\text{-lasR-}C_]] = N_i$, we can derive the rate of production of mRNA as a function of $[R]_T$, $[S]_T$ $[C_6]$ and $[C_{12}]$. For notational convenience we drop the square brackets and $T$ subscript, which allows us to write

$$f_i(R,S,C_6,C_{12}) = \frac{\epsilon^{(i)} + K_{GR}^{(i)} R^2 \left(\frac{K_{Rk}C_k}{1+\sum_k K_{Rk}C_k}\right)^{n_R} + K_{GS}^{(i)} S^2 \left(\frac{K_{Sk}C_k}{1+\sum_k K_{Sk}C_k}\right)^{n_S}}{1 + K_{GR}^{(i)} R^2 \left(\frac{K_{Rk}C_k}{1+\sum_k K_{Rk}C_k}\right)^{n_R} + K_{GS}^{(i)} S^2 \left(\frac{K_{Rk}C_k}{1+\sum_k K_{Rk}C_k}\right)^{n_S}} \tag{S18}$$

where $K_{G_i\text{-luxR}}$ and $K_{G_i\text{-lasR}}$ are scaled to incorporate $K_{Dk}$ and $K_{Ek}$ respectively, and we introduce alternative exponents $n_R$ and $n_S$, analogous to the usage of $n$ in [3]. Accordingly, we obtain the following system of equations

$$\frac{dc}{dt} = \gamma(c).c \tag{S19a}$$

$$\frac{d[R]}{dt} = a_R.h(c) - (d_R + \gamma(c))[R] \tag{S19b}$$

$$\frac{d[S]}{dt} = a_S.h(c) - (d_R + \gamma(c))[S] \tag{S19c}$$

$$\frac{d[\text{CFP}]}{dt} = a_{\text{CFP}}.h(c).f_{76}([C_6],[C_{12}],[R],[S]) - (d_{\text{CFP}} + \gamma(c))[\text{CFP}] \tag{S19d}$$

$$\frac{d[\text{YFP}]}{dt} = a_{\text{YFP}}.h(c).f_{81}([C_6],[C_{12}],[R],[S]) - (d_{\text{YFP}} + \gamma(c))[\text{YFP}] \tag{S19e}$$

where the variables $R$ and $S$ now represent total luxR and lasR respectively, and $\gamma$ is as defined in (6).

##### Implementation with GEC/CRN language

The DR circuit introduces expression of LuxR and LasR proteins, so we define modules for their synthesis, taking the same strategy as done for eYFP /eCFP in the PRPR  circuit above. However, as we only use the same promoters across all circuits in this paper, and they are used in a non-inducible way, there is no need to have a rate expression that multiplies the overall expression rate.

```
module LuxR(aR,growth,capacity) = {
    ->[[capacity]*aR] luxR |
    luxR ->{dR} |
    luxR ->[[growth]*[luxR]]
}
module LasR(aS,growth,capacity) = {
    ->[[capacity]*aS] lasR |
    lasR ->{dR} |
    lasR ->[[growth]*[lasR]]
}
```

The overall models for the DR circuits combines calls to these LuxR/LasR modules with calls to the eYFP /eCFP modules, but the latter uses more complex rate expressions that describe the complex inducible nature of the modified PLux promoters. As different DR circuits have different ribosome binding sites for LuxR and LasR expression, we use different values of `aR` and `aS`. Finally, we specify the initial conditions for C6 (species `c6`) and C12 (species `c12`) using parameters `C6` and `C12`, which are set differently for each trace.

```
directive rates [
    boundLuxR = [luxR]^2 * ((KR6*[c6])^nR + (KR12*[c12])^nR) / ((1.0 + KR6*[c6] + KR12*[c12])^nR);
    boundLasR = [lasR]^2 * ((KS6*[c6])^nS + (KS12*[c12])^nS) / ((1.0 + KS6*[c6] + KS12*[c12])^nS);
    P76 = (e76 + KGR_76*[boundLuxR] + KGS_76*[boundLasR]) / (1.0 + KGR_76*[boundLuxR] +
        KGS_76*[boundLasR]);
```

```
    P81 = (e81 + KGR_81*[boundLuxR] + KGS_81*[boundLasR]) / (1.0 + KGR_81*[boundLuxR] +
        KGS_81*[boundLasR]);
]
device drPcat() = { LuxR(1.0,growth,capacity) | LasR(1.0,growth,capacity) |
    YFP(P81,aYFP,growth,capacity) | CFP(P76,aCFP,growth,capacity) }
device drRS100S32() = { LuxR(aRS100,growth,capacity) | LasR(aS32,growth,capacity) |
    YFP(P81,aYFP,growth,capacity) | CFP(P76,aCFP,growth,capacity) }
device drR33S32() = { LuxR(aR33,growth,capacity) | LasR(aS32,growth,capacity) |
    YFP(P81,aYFP,growth,capacity) | CFP(P76,aCFP,growth,capacity) }
device drR33S175() = { LuxR(aR33,growth,capacity) | LasR(aS175,growth,capacity) |
    YFP(P81,aYFP,growth,capacity) | CFP(P76,aCFP,growth,capacity) }
```

##### Characterization experiment

To characterize the LuxR and LasR signalling components, we used measurements of the response of four DR circuits from [3] to treatment with C6 and C12 over 3-fold dilutions. The maximum LuxR and LasR production rates were normalized to the values corresponding to the Pcat promoters, as done previously [3], thus setting the scale for unobserved concentrations of LuxR and LasR.

We used (uninformative) uniform priors on the previously uncharacterized parameters, and (informative) truncated Gaussian priors on $f_{480}$, $f_{530}$, $d_{\mathrm{CFP}}$ and $d_{\mathrm{YFP}}$ with mean and standard deviation taken from the marginal posteriors of the PRPR circuit characterization. These parameters are globally specified, with propagation of marginal posteriors as edges of the inference graph (see Listing 1). We also use uninformative priors on the $B_k^{\mathrm{back}}$ parameters (specified below as local parameters b530 and b480), without propagating their marginal posteriors from previous circuits.

As in the previous section, each system definition incorporates device invocations in combination with module calls for cell growth and autofluorescence.

```
// DR (Pcat-Pcat)
system dr_PcatPcat = {
    directive simulation { final=36.0; points=250; plots=[[x]*([yfp]+[f530])+b530;
        [x]*([cfp]+[f480])+b480]; plotcolours=["#FFDF00"; "#ADD8E6"] }
    directive parameters [
        r = 1, { interval=Real; distribution=Uniform(0.1,10); variation=Multiple };
        K = 2, { interval=Real; distribution=Uniform(0.1,5); variation=Multiple };
        tlag = 1, { interval=Real; distribution=Uniform(0,10); variation=Multiple };
        rc=1e2,  { interval=Log; distribution=Uniform(1e0,1e5); variation=Multiple };
        b530=1e3, { interval=Real; distribution=Uniform(0.0,5e3); variation=Random };
        b480=1e3, { interval=Real; distribution=Uniform(0.0,1e4); variation=Random };
    ]
    directive rates [growth = [grow]*r*(1 - [x] / K);capacity = rc]
    directive crn { cells(growth,tlag) | autofluorescence(growth,capacity) }
    | drPcat:device
}

// DR (R100-S32)
system dr_R100S32 = {
    directive simulation { final=36.0; points=250; plots=[[x]*([yfp]+[f530])+b530;
        [x]*([cfp]+[f480])+b480]; plotcolours=["#FFDF00"; "#ADD8E6"] }
    directive parameters [
        r = 1, { interval=Real; distribution=Uniform(0.1,10); variation=Multiple };
        K = 2, { interval=Real; distribution=Uniform(0.1,5); variation=Multiple };
        tlag = 1, { interval=Real; distribution=Uniform(0,10); variation=Multiple };
        rc=1e2,  { interval=Log; distribution=Uniform(1e0,1e5); variation=Multiple };
        b530=1e3, { interval=Real; distribution=Uniform(0.0,5e3); variation=Random };
        b480=1e3, { interval=Real; distribution=Uniform(0.0,1e4); variation=Random };
    ]
    directive rates [growth = [grow]*r*(1 - [x] / K);capacity = rc]
```

```
    directive crn { cells(growth,tlag) | autofluorescence(growth,capacity) }
    | drRS100S32:device
}

// DR (R33-S32)
system dr_R33S32 = {
    directive simulation { final=36.0; points=250; plots=[[x]*([yfp]+[f530])+b530;
        [x]*([cfp]+[f480])+b480]; plotcolours=["#FFDF00"; "#ADD8E6"] }
    directive parameters [
        r = 1, { interval=Real; distribution=Uniform(0.1,10); variation=Multiple };
        K = 2, { interval=Real; distribution=Uniform(0.1,5); variation=Multiple };
        tlag = 1, { interval=Real; distribution=Uniform(0,10); variation=Multiple };
        rc=1e2,  { interval=Log; distribution=Uniform(1e0,1e5); variation=Multiple };
        b530=1e3, { interval=Real; distribution=Uniform(0.0,5e3); variation=Random };
        b480=1e3, { interval=Real; distribution=Uniform(0.0,1e4); variation=Random };
    ]
    directive rates [growth = [grow]*r*(1 - [x] / K);capacity = rc]
    directive crn { cells(growth,tlag) | autofluorescence(growth,capacity) }
    | drR33S32:device
}

// DR (R33-S175)
system dr_R33S175 = {
    directive simulation { final=36.0; points=250; plots=[[x]*([yfp]+[f530])+b530;
        [x]*([cfp]+[f480])+b480]; plotcolours=["#FFDF00"; "#ADD8E6"] }
    directive parameters [
        r = 1, { interval=Real; distribution=Uniform(0.1,10); variation=Multiple };
        K = 2, { interval=Real; distribution=Uniform(0.1,5); variation=Multiple };
        tlag = 1, { interval=Real; distribution=Uniform(0,10); variation=Multiple };
        rc=1e2,  { interval=Log; distribution=Uniform(1e0,1e5); variation=Multiple };
        b530=1e3, { interval=Real; distribution=Uniform(0.0,5e3); variation=Random };
        b480=1e3, { interval=Real; distribution=Uniform(0.0,1e4); variation=Random };
    ]
    directive rates [growth = [grow]*r*(1 - [x] / K);capacity = rc]
    directive crn { cells(growth,tlag) | autofluorescence(growth,capacity) }
    | drR33S175:device
}
```

##### Characterization results

In the subsequent figures we first show model-data comparisons for the TargetSwitch hypothesis applied using direct (Fig. S15) and ratiometric characterization (Figs. S16 & S17). Next, we show the marginal posterior distributions for each parameter (Figs. S18 & S19).

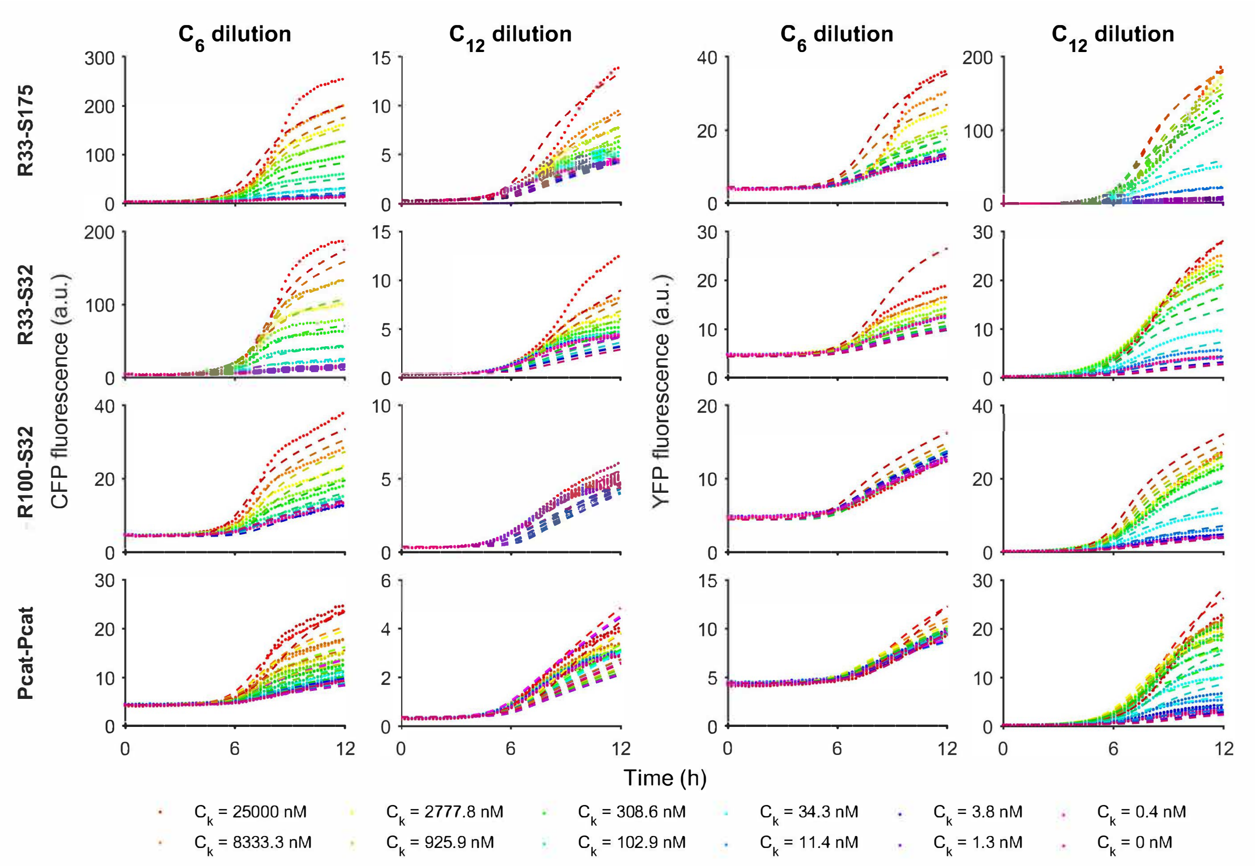

**Figure S15: Model-data comparison for Receivers using direct dynamic characterization with the TargetSwitch hypothesis.**

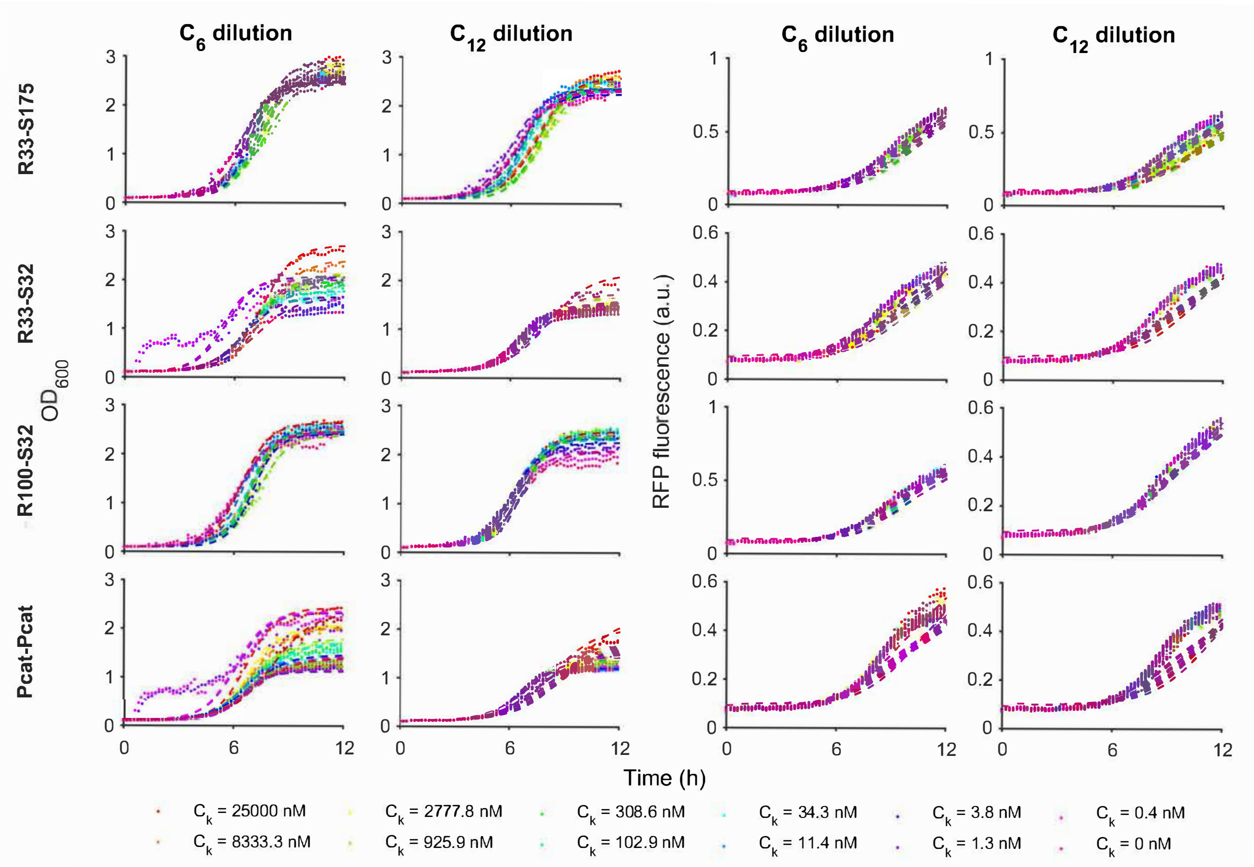

**Figure S16: Model-data comparison for Receivers using ratiometric dynamic characterization with the TargetSwitch hypothesis (Growth and Control phases).**

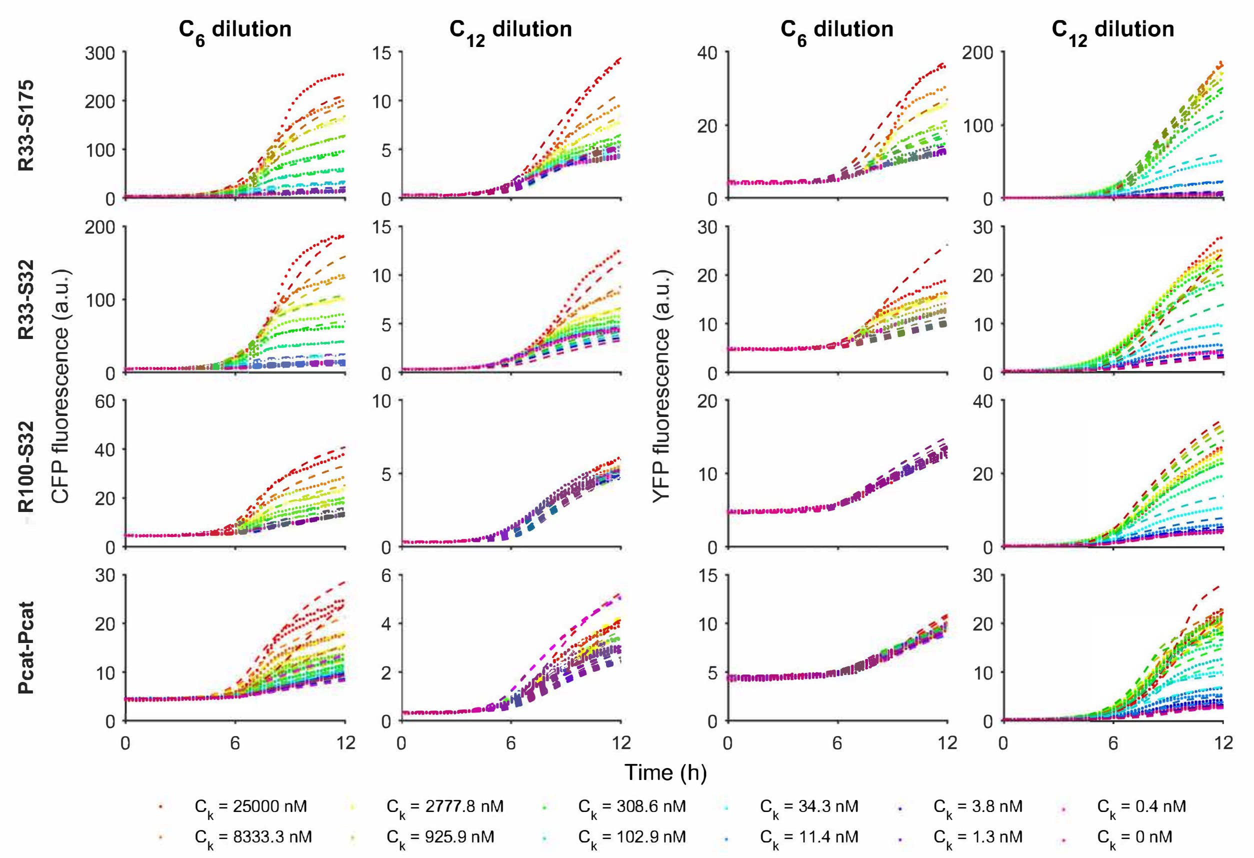

**Figure S17: Model-data comparison for Receivers using ratiometric dynamic characterization with the TargetSwitch hypothesis.**

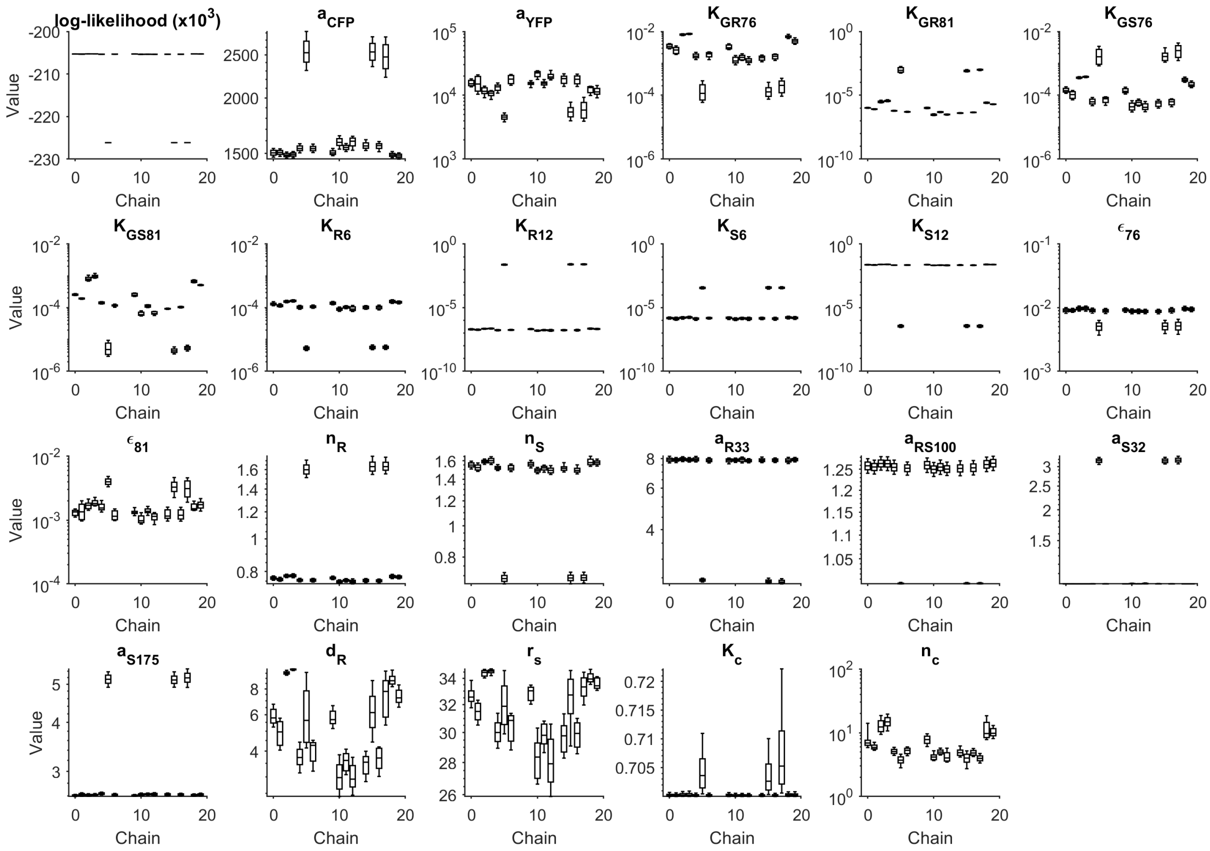

**Figure S18: Marginal parameter posterior distributions for the DR circuit with TargetSwitch using the direct method.** Inference of circuit parameters using the direct (non-ratiometric) method, which corresponds to assuming that $r_c = 1$ for all time-series traces.

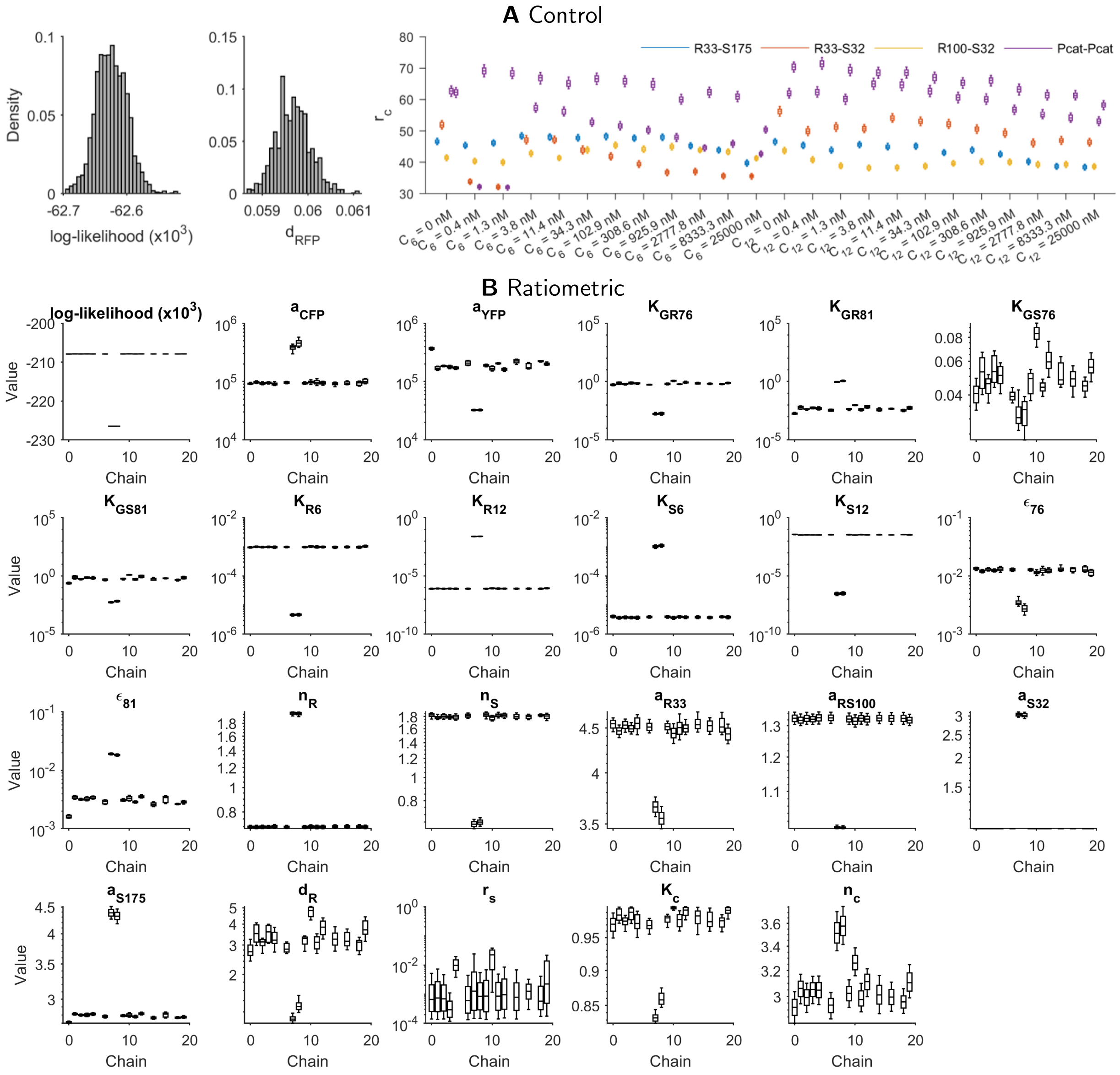

**Figure S19: Marginal parameter posterior distributions for the DR circuit with TargetSwitch using the ratiometric method. A.** The control phase identifies values for $r_c$ for each time-series, to be used in ratiometric characterization. **B.** Inference of circuit parameters using the ratiometric method, using the values of $r_c$ in B.

#### S2.4 AHL senders (Relays)

To characterize the dynamics of AHL senders, we constructed two *relay* signalling devices, each perceiving a first signal that acts as an inducer for a second signal. The first relay device responds to C6 via binding of constitutively expressed LuxR and activation of the engineered PLux76 promoter, then synthesizes the C12-producing LasI enzyme and eCFP . C12 is then detected via binding of constitutively expressed LasR and activation of the engineered PLas81 promoter, which synthesizes eYFP . Accordingly, we can monitor the promoter activity of both stages of the relay signals. The second device performs receive-send-receive in the other order, sensing C12 via PLas81, which synthesizes the C6-producing LuxI enzyme (and eYFP ), with C6 then detected via PLux76-eCFP . Both relay circuits embed the double receiver module, along with an HSL sender for C6 or for C12 (Fig. S20). As such, the parameter posterior distributions learned during characterization of the double receiver module are relevant prior distributions for these circuits.

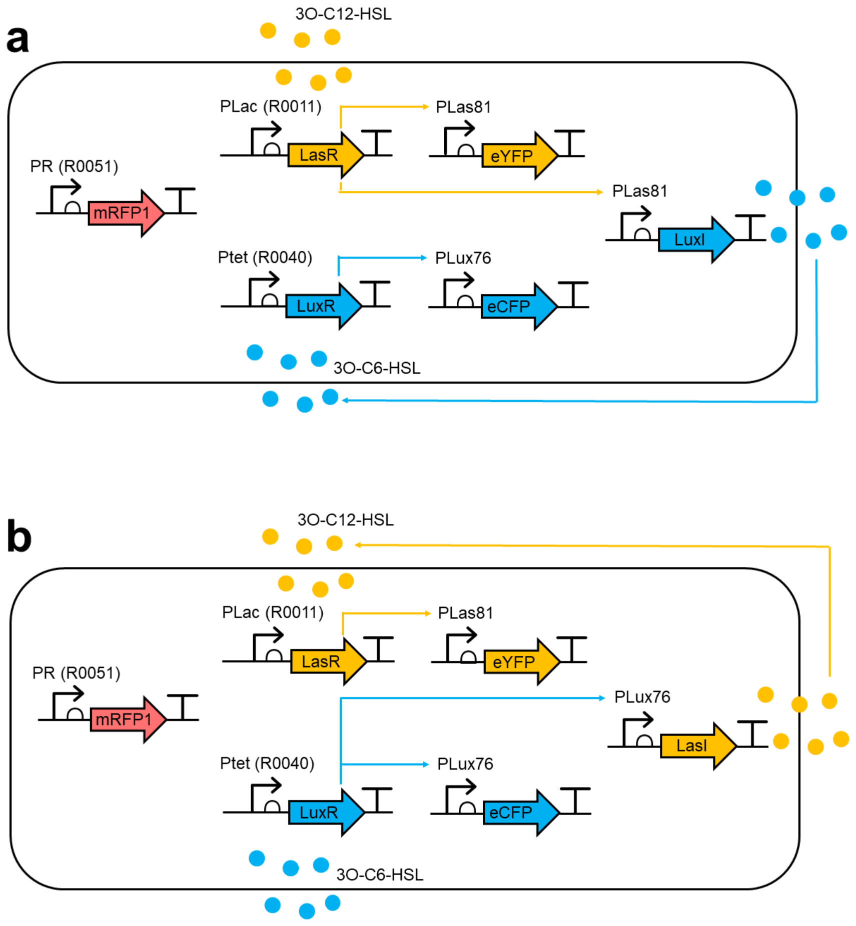

**Figure S20: Relay circuit network diagrams. A**. The PLas81-LuxI relay device, with double reporter. **B**. The PLux76-LasI device, with double reporter.

##### Model definition

The models for the relay circuits are modular: the double receiver model from Section S2.3, is extended by equations for luxI and lasI, and additional terms for $C_6$ and $C_{12}$ synthesis. Building on the approach above, we can write down an equation for the intracellular production of LuxI or LasI. Each then synthesizes C6 and C12 respectively, which we model as a linear function of [luxI] or [lasI]. In the models described above, it was assumed that transport between the intracellular and extracellular compartments was fast. As there was no *de novo* synthesis of HSL, and often a large extracellular excess, the system dynamics would unlikely be critically sensitive to this approximation, as any lag could be accounted for by adjustments in the other parameters. However, in the relay circuits, where HSLs are being produced, the transport between intracellular and extracellular compartments must be considered more carefully. For the C6-LasI-C12 relay, the equations for the mass of each molecule in the intracellular (denoted by subscript $i$) and extracellular (denoted by subscript $e$) compartments is given by

$$\frac{d(V_i.[\text{lasI}])}{dt} = V_i.h\left(\frac{V_i}{V_{\text{tot}}}\right).\rho_L.f_{76}([\text{C}_6],[\text{C}_{12}],[\text{luxR}],[\text{lasR}]) - V_i.d_L[\text{lasI}] \tag{S20a}$$

$$\frac{d(V_i.[\text{C}_{12}]_i)}{dt} = V_i.k_{\text{C12}}.[\text{lasI}] + \eta_{12}\left([\text{C}_{12}]_e - [\text{C}_{12}]_i\right) \tag{S20b}$$

$$\frac{d(V_e.[\text{C}_{12}]_e)}{dt} = \eta_{12}\left([\text{C}_{12}]_i - [\text{C}_{12}]_e\right) \tag{S20c}$$

where $\rho_L$ is the relative production rate of lasI (relative to eCFP production; see above), and $k_{C12}$ is the rate of C12 synthesis by lasI. Here, $V_i$ and $V_e$ are the volumes of intracellular and extracellular compartments respectively. Accordingly, cell density $c$ is simply the ratio of $V_i$ to $V_i + V_e =: V_{\text{tot}}$.

To arrive at a set of equations for the derivatives of the concentrations of each molecule (in each compartment), we expand the left-hand sides using the chain rule. Then, defining $\gamma := \frac{1}{V_i}\frac{dV_i}{dt}$ as the specific growth rate, or *dilution* factor, assuming that $V_e = V_{\text{tot}} - V_i$, and rescaling [lasI] according to $[\text{lasI}] = \frac{L}{\rho_L}$ we obtain

$$\frac{dL}{dt} = h\left(\frac{V_i}{V_{\text{tot}}}\right).f_{76}([\text{C}_6],[\text{C}_{12}],[\text{luxR}],[\text{lasR}]) - (d_L + \gamma)L \tag{S21a}$$

$$\frac{d[\text{C}_{12}]_i}{dt} = k'_{\text{C12}}.L + \frac{\eta_{12}}{V_i}\left([\text{C}_{12}]_e - [\text{C}_{12}]_i\right) - \gamma[\text{C}_{12}]_i \tag{S21b}$$

$$\frac{d[\text{C}_{12}]_e}{dt} = \frac{\eta_{12}}{V_{\text{tot}} - V_i}\left([\text{C}_{12}]_i - [\text{C}_{12}]_e\right) + \frac{\gamma V_i}{V_{\text{tot}} - V_i} \tag{S21c}$$

where $k'_{\text{C12}} = \frac{k_{C12}}{\rho_L}$.

As we had no reliable estimate of HSL transport rates *a priori*, we started by using the model that combines (S21) with (S19). However, we also considered the assumption that HSL transport is fast, resulting in the relations $[C_k]_e = [C_k]_i = [C_k]$ for $k = 6, 12$. By summing equations (S20b) and (S20c), and using an equivalent rescaling for [lasI] as before, we obtain

$$\frac{d(V_{\text{tot}}.[\text{C}_{12}])}{dt} = \frac{d(V_i.[\text{C}_{12}]_i)}{dt} + \frac{d(V_e.[\text{C}_{12}]_e)}{dt} = V_i.k_{\text{C12}}.L$$

which when rearranged and gives an alternative to model (S21) as

$$\frac{dL}{dt} = h(c).f_{76}([\text{C}_6],[\text{C}_{12}],[\text{luxR}],[\text{lasR}]) - (d_L + \gamma)L \tag{S22a}$$

$$\frac{d[\text{C}_{12}]}{dt} = k'_{\text{C12}}.c.L \tag{S22b}$$

with $k'_{\text{C12}}$ and $L$ as defined above.

**Saturation of HSL substrate usage.** If the luxI or lasI enzymes are expressed very highly, it might be that substrate becomes the limiting factor for HSL synthesis (as opposed to enzyme). In which case, a saturating production term should be used, which is parametrized by a new parameter $K_L$, the concentration of luxI/lasI at which HSL production is half-maximal. We can define this function as

$$f(L) = k'_{\text{C12}}.\frac{L}{L + K_L} \tag{S23}$$

where $k'_{\text{C12}}$ is the rescaled (maximal) per-enzyme rate of HSL synthesis from lasI, as above.

Finally, we note that the equations above can be trivially concatenated with the equations of the double receiver device described in the previous section to produce a full model of each relay device.

##### Implementation with GEC/CRN language

We define modules for LuxI and LasI enzymes, which includes their modelled enzymatic synthesis of C6 and C12. As before, we use a rate expression argument to arbitrarily assign a promoter transfer function to the production rates of each enzyme. However, since LuxI and LasI are never observed, their maximal production rates can be *divided out*, as explained above, and without loss of generality set to a value 1.

```
module LuxI(P,growth,capacity) = {
    ->[[capacity]*[P]] luxI |
    luxI ->{dluxI} |
    luxI ->[[growth]*[luxI]] |
    ->[kC6*[capacity]*[x]*[luxI]/(1+[luxI]/Klux)] c6
}
module LasI(P,growth,capacity) = {
    ->[[capacity]*[P]] lasI |
    lasI ->{dlasI} |
    lasI ->[[growth]*[lasI]] |
    ->[kC12*[capacity]*[x]*[lasI]/(1+[lasI]/Klas)] c12
}

device drR33S175() = { LuxR(aR33,growth,capacity) | LasR(aS175,growth,capacity) |
     YFP(P81,aYFP,growth,capacity) | CFP(P76,aCFP,growth,capacity) }
device relayP76LasI() = { LasI(P76,growth,capacity) }
device relayP81LuxI() = { LuxI(P81,growth,capacity) }
```

##### Characterization experiment

To characterize the Relay devices, we measured each in response to C6 and C12 treatments. We used (uninformative) uniform priors on the previously uncharacterized parameters ($k_{C6}$, $k_{C12}$, $K_{lux}$, $K_{las}$, $d_{luxI}$ and $d_{lasI}$), and (informative) truncated Gaussian priors on all remaining parameters, with mean and standard deviation taken from the marginal posteriors of the DR circuits characterization. We also use uninformative priors on the background fluorescence parameters (specified below as local parameters b530 and b480), without propagating their marginal posteriors from previous circuits.

To instantiate the relay circuits, we use device definitions as above, this time taking advantage of device modularity.

```
system relay_P76LasI = {
    directive simulation { final=36.0; points=250; plots=[[x]*([yfp]+[f530])+b530;
         [x]*([cfp]+[f480])+b480]; plotcolours=["#FFDF00"; "#ADD8E6"] }
    directive parameters [
        r = 1, { interval=Real; distribution=Uniform(0.1,10); variation=Multiple };
        K = 2, { interval=Real; distribution=Uniform(0.1,5); variation=Multiple };
        tlag = 1, { interval=Real; distribution=Uniform(0,10); variation=Multiple };
        rc=1e2,  { interval=Log; distribution=Uniform(1e0,1e5); variation=Multiple };
        b530=1e3, { interval=Real; distribution=Uniform(0.0,5e3); variation=Random };
        b480=1e3, { interval=Real; distribution=Uniform(0.0,1e4); variation=Random };
    ]
    directive rates [growth = [grow]*r*(1 - [x] / K);capacity = rc]
    directive crn { cells(growth,tlag) | autofluorescence(growth,capacity) }
    | drR33S175:device
    | relayP76LasI:device
}

system relay_P81LuxI = {
    directive simulation { final=36.0; points=250; plots=[[x]*([yfp]+[f530])+b530;
         [x]*([cfp]+[f480])+b480]; plotcolours=["#FFDF00"; "#ADD8E6"] }
    directive parameters [
        r = 1, { interval=Real; distribution=Uniform(0.1,10); variation=Multiple };
```

```
        K = 2, { interval=Real; distribution=Uniform(0.1,5); variation=Multiple };
        tlag = 1, { interval=Real; distribution=Uniform(0,10); variation=Multiple };
        rc=1e2,  { interval=Log; distribution=Uniform(1e0,1e5); variation=Multiple };
        b530=1e3, { interval=Real; distribution=Uniform(0.0,5e3); variation=Random };
        b480=1e3, { interval=Real; distribution=Uniform(0.0,1e4); variation=Random };
    ]
    directive rates [growth = [grow]*r*(1 - [x] / K);capacity = rc]
    directive crn { cells(growth,tlag) | autofluorescence(growth,capacity) }
    | drR33S175:device
    | relayP81LuxI:device
}
```

##### Characterization results

In the subsequent figures we first show model-data comparisons for the TargetSwitch hypothesis applied using direct (Fig. S21A) and ratiometric characterization (Fig. S21B,C). Next, we show the marginal posterior distributions for each parameter (Fig. S22).

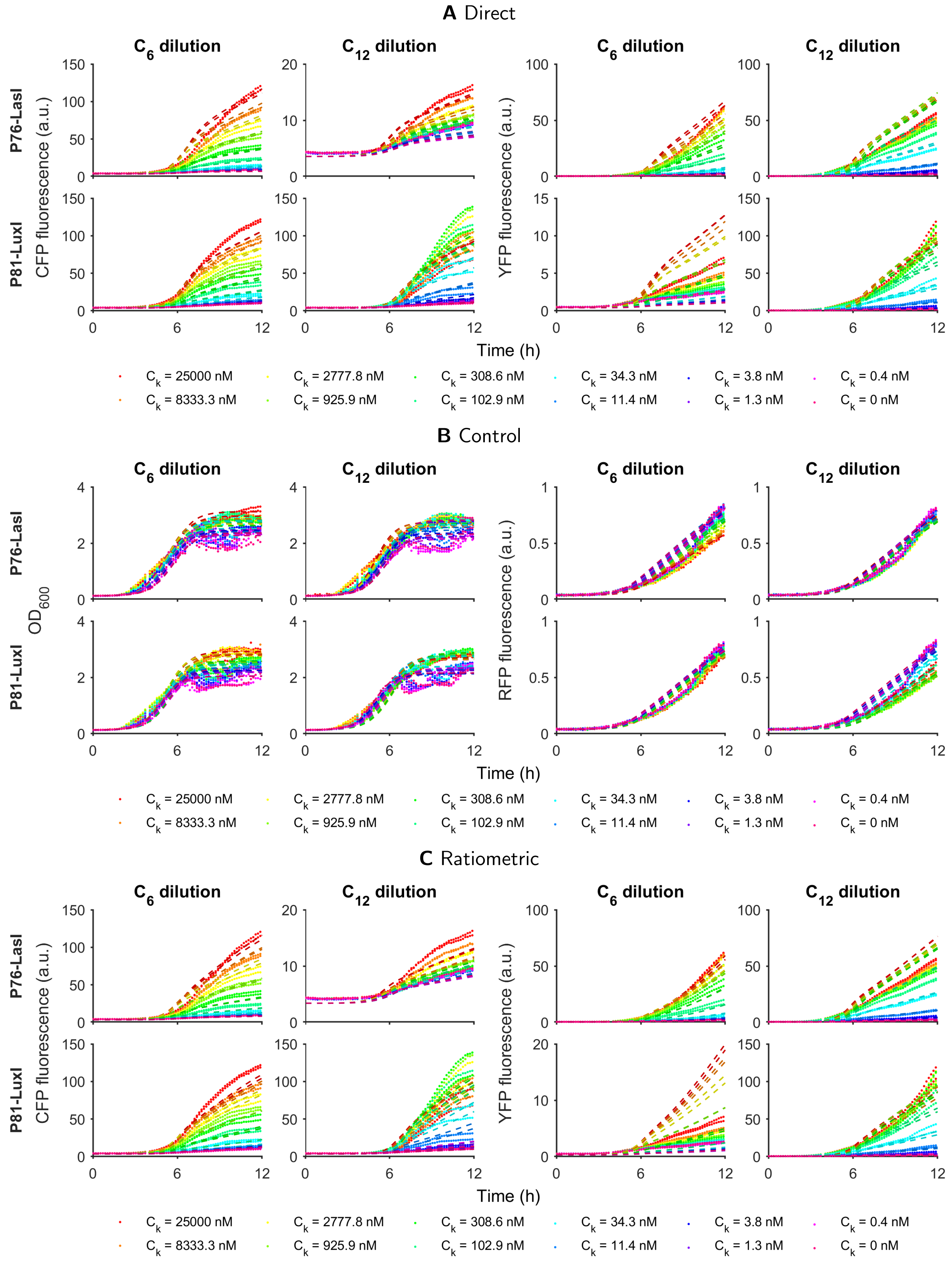

**Figure S21: Model-data comparison for the Relays circuit with TargetSwitch.** Measurements of the Relays circuit are compared against simulations of the maximum likelihood parameter set from the TargetGrowth hypothesis.

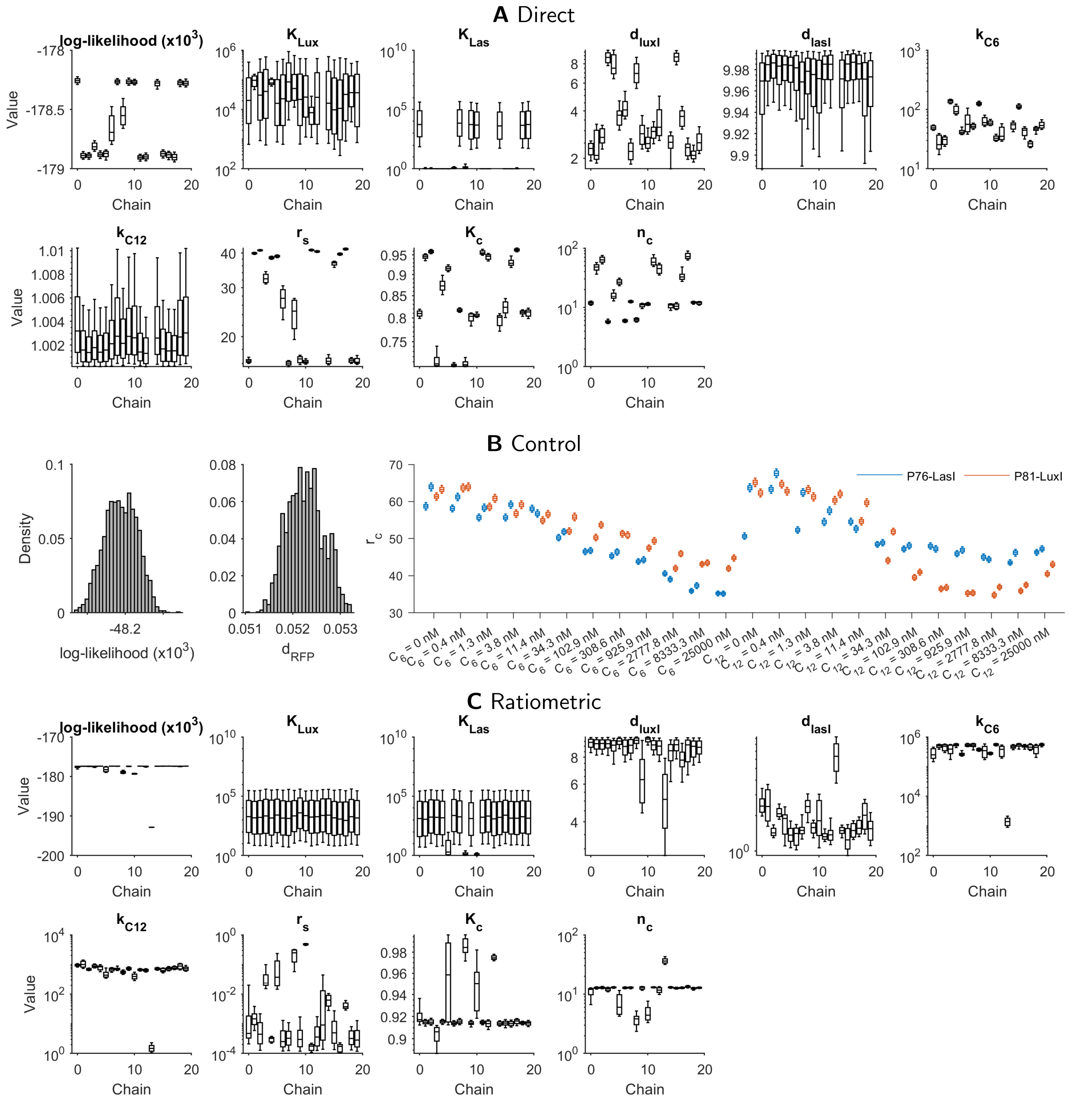

**Figure S22: Marginal parameter posterior distributions for Relays with TargetSwitch hypothesis.** **A.** Inference of circuit parameters using the direct (non-ratiometric) method, which corresponds to assuming that $r_c = 1$ for all time-series traces. **B.** The control phase identifies values for $r_c$ for each time-series, to be used in ratiometric characterization. **C.** Inference of circuit parameters using the ratiometric method, using the values of $r_c$ in B.

#### S2.5 Inducible expression (PBAD)

The arabinose-PBAD system is commonly used to control expression in synthetic gene circuits. The PBAD promoter is part of the arabinose operon, which also involves the araC protein. When arabinose is added to cell cultures at different concentrations, it increases expression of a downstream transcript. In this study, we have used PBAD to control AHL lactonase expression (see Section S2.6 for details).

To characterize the relationship between PBAD and arabinose in vivo, we constructed the Inducer circuit, which has the coding sequence for eYFP downstream of PBAD, and measured its response to varying concentrations of arabinose.

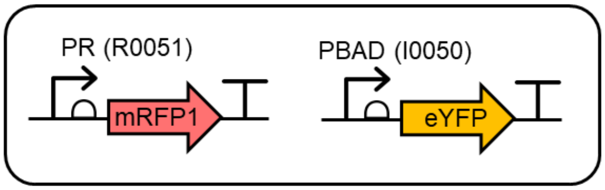

**Figure S23: Design of Inducer circuit.**

##### Characterization experiment

We measured the Inducer circuit in response to varying concentrations of the arabinose inducer. We used (uninformative) uniform priors on the previously uncharacterized parameters, and (informative) truncated Gaussian priors on $a_{480}$ and $a_{530}$ with mean and standard deviation taken from the marginal posteriors of the Constitutive circuit characterization. We also use uninformative priors on the background fluorescence parameters, without propagating their marginal posteriors from previous circuits.

##### Implementation with GEC/CRN language

As the module for describing eYFP expression has already been defined, it can be used directly to describe PBAD induction. All that is required is to define a rate expression for the transfer function that describes PBAD activity in response to varying concentrations of arabinose, and define a new parameter that quantifies maximal YFP production in the context of a PBAD promoter (with B0034 ribosome binding site). Accordingly, the Inducer circuit is described by

```
directive rates [
    PBAD = (Ara^nA+eA*KAra^nA)/(Ara^nA+KAra^nA);
]

device pBADYFP() = { YFP(PBAD,aYFP,growth,capacity) }

system PBADYFP = { target with
    directive simulation { final=36.0; points=250; plots=[[x]*([yfp]+[f530])+b530];
         plotcolours=["#FFDF00"] }
    directive parameters [
        r = 1, { interval=Real; distribution=Uniform(0.1,10); variation=Multiple };
        K = 2, { interval=Real; distribution=Uniform(0.1,5); variation=Multiple };
        tlag = 1, { interval=Real; distribution=Uniform(0,10); variation=Multiple };
        rc=1e2,  { interval=Log; distribution=Uniform(1e0,1e5); variation=Multiple };
        aYFP=1.0, { interval=Log; distribution=Uniform(1e0,1e5); variation=Random };
        b530=1e3, { interval=Real; distribution=Uniform(0.0,5e3); variation=Random };
    ]
    directive rates [growth = [grow]*r*(1 - [x] / K);capacity = rc]
    directive crn { cells(growth,tlag) | autofluorescence(growth,capacity) }
    pBadYFP:device
}
```

#### Characterization results

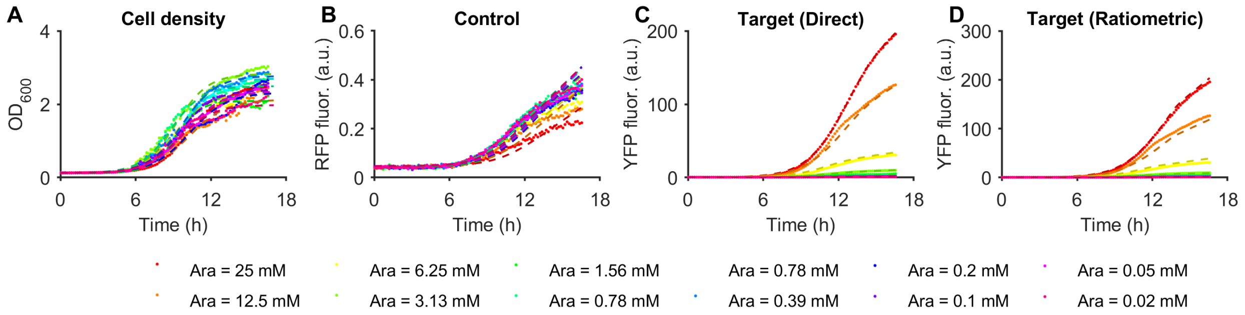

**Figure S24: Model-data comparison for the Inducer circuit with TargetSwitch hypothesis.** Measurements of the PBADYFP circuit are compared against simulations of the maximum likelihood parameter set from applying ratiometric dynamic characterization with the TargetSwitch hypothesis.

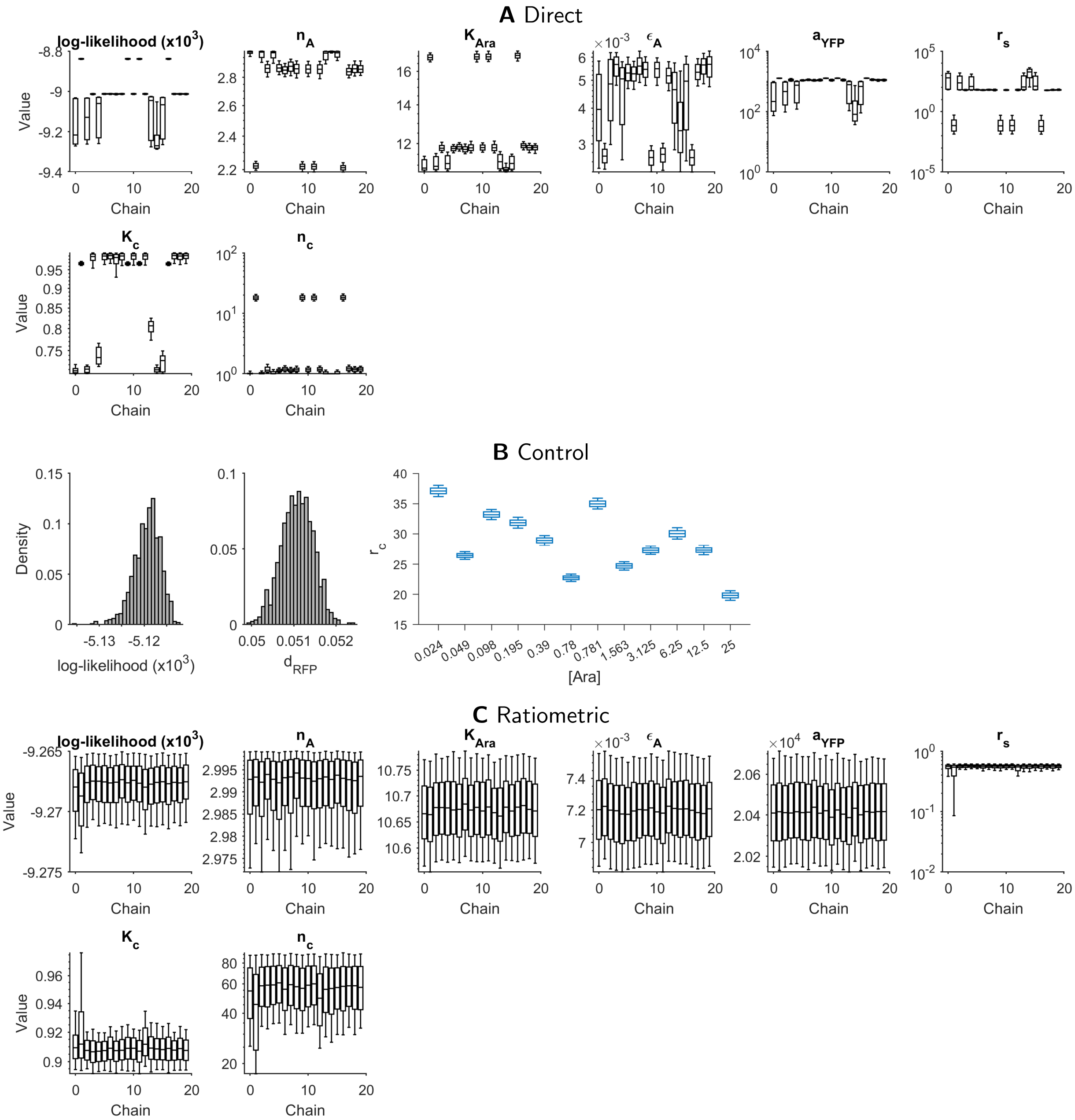

**Figure S25: Marginal parameter posterior distributions for the Inducer circuit with TargetSwitch hypothesis. A.** Inference of circuit parameters using the direct (non-ratiometric) method, which corresponds to assuming that $r_c = 1$ for all time-series traces. **B.** The control phase identifies values for $r_c$ for each time-series, to be used in ratiometric characterization. **C.** Inference of circuit parameters using the ratiometric method, using the values of $r_c$ in B.

#### S2.6 HSL lactonase (AiiA)

AHL molecules can be degraded by AHL lactonase enzymes. The most commonly used lactonase in synthetic biology applications is AiiA [5, 6], which originates from *Bacillus Thurigensis*. AiiA is mostly non-specific; catalytic activity has been measured against a large variety of AHLs with varying length carbon chains [7].

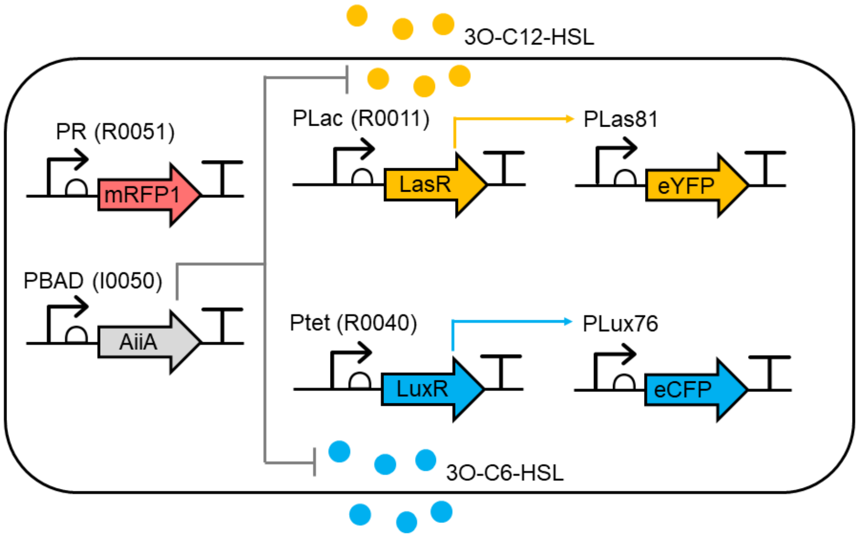

**Figure S26: Network diagram for the Degrader circuit.**

To characterize the AiiA lactonase activity in a synthetic gene circuit, we constructed a PBAD-AiiA plasmid, and combined this with the R33-S175 plasmid to define an arabinose-inducible Degrader circuit (Fig. S26).

It is fairly straightforward to write down a few reactions describing the production, degradation and catalytic activity of AiiA. Assuming constitutive production, we propose:

$$\varnothing \underset{d_A}{\overset{a_A \cdot f_{\text{Pbad}}(\text{Ara})}{\rightleftharpoons}} \text{AiiA} \tag{S24a}$$

$$C_6 + \text{AiiA} \underset{u_{A6}}{\overset{b_{A6}}{\rightleftharpoons}} C_6.\text{AiiA} \xrightarrow{d_{A6}} \text{AiiA} \tag{S24b}$$

$$C_{12} + \text{AiiA} \underset{u_{A12}}{\overset{b_{A12}}{\rightleftharpoons}} C_{12}.\text{AiiA} \xrightarrow{d_{A12}} \text{AiiA} \tag{S24c}$$

As AHLs may translocate between cells and the extracellular medium, and AiiA is only expressed inside cells, we must carefully consider the volume and concentration of these molecules over time, as before.

Translating the above reactions to ODEs, we obtain

$$\frac{d(V_i.[A])}{dt} = V_i.a_A.f_{\text{Pbad}}(\text{Ara}) + V_i.(u_{A6} + d_{A6})[A.C_6] + V_i.(u_{A12} + d_{A12})[A.C_{12}] \dots \tag{S25a}$$

$$- V_i.(d_A + b_{A6}[C_6]_i + b_{A12}[C_{12}]_i)[A] \tag{S25b}$$

$$\frac{d(V_i.[C_6]_i)}{dt} = V_i.u_{A6}.[A.C_6] + \eta_6\left([C_6]_e - [C_6]_i\right) - V_i.(d_C + b_{A6}[A])[C_6]_i \tag{S25c}$$

$$\frac{d(V_i.[C_{12}]_i)}{dt} = V_i.u_{A12}.[A.C_{12}] + \eta_{12}\left([C_{12}]_e - [C_{12}]_i\right) - V_i.(d_C + b_{A12}[A])[C_{12}]_i \tag{S25d}$$

$$\frac{d(V_i.[A.C_6])}{dt} = V_i.b_{A6}[A][C_6]_i - V_i.(u_{A6} + d_{A6})[A.C_6] \tag{S25e}$$

$$\frac{d(V_i.[A.C_{12}])}{dt} = V_i.b_{A6}[A][C_{12}]_i - V_i.(u_{A12} + d_{A12})[A.C_{12}] \tag{S25f}$$

$$\frac{d(V_e.[C_6]_e)}{dt} = \eta_6\left([C_6]_i - [C_6]_e\right) \tag{S25g}$$

$$\frac{d(V_e.[C_{12}]_e)}{dt} = \eta_{12}\left([C_{12}]_i - [C_{12}]_e\right) \tag{S25h}$$

We can apply an equilibrium assumption to the AiiA-HSL intermediate by equating (S25e) and (S25f) to zero. This gives

$$[A.C_j] = \frac{b_{Aj}}{\gamma + u_{Aj} + d_{Aj}}[A][C_j] \approx \frac{b_{Aj}}{u_{Aj} + d_{Aj}}[A][C_j] \tag{S26}$$

where the approximation to suppress $\gamma$ is due to the catalytic activity of AiiA being significantly faster than the dilution rate. Therefore, depending on the concentrations $[C_6]$ and $[C_{12}]$, AiiA is bound according to the conservation equation

$$[A_{\text{Tot}}] = [A] + [A.C_6] + [A.C_{12}] = [A].(1 + K_{A6}[C_6] + K_{A12}[C_{12}]) \tag{S27}$$

where $K_{Aj} = \frac{b_{Aj}}{u_{Aj}+d_{Aj}}$. By cancelling the binding and unbinding terms with AiiA, we are left with the simpler set of equations

$$\frac{d(V_i.[A_{\text{Tot}}])}{dt} = V_i.a_A.f_{\text{Pbad}}(\text{Ara}) - V_i.d_A.\frac{[A_{\text{Tot}}]}{1 + K_{A6}[C_6] + K_{A12}[C_{12}]} \tag{S28a}$$

$$\frac{d(V_i.[C_6]_i)}{dt} = \eta_6\left([C_6]_e - [C_6]_i\right) - V_i.d_{A6}.\frac{[A_{\text{Tot}}][C_6]_i}{1 + K_{A6}[C_6]_i + K_{A12}[C_{12}]_i} - V_i.d_C.[C_j] \tag{S28b}$$

$$\frac{d(V_i.[C_{12}]_i)}{dt} = \eta_{12}\left([C_{12}]_e - [C_{12}]_i\right) - V_i.d_{A12}.\frac{[A_{\text{Tot}}][C_{12}]_i}{1 + K_{A6}[C_6]_i + K_{A12}[C_{12}]_i} - V_i.d_C.[C_j] \tag{S28c}$$

$$\frac{d(V_e.[C_6]_e)}{dt} = \eta_6\left([C_6]_i - [C_6]_e\right) \tag{S28d}$$

$$\frac{d(V_e.[C_{12}]_e)}{dt} = \eta_{12}\left([C_{12}]_i - [C_6]_e\right) \tag{S28e}$$

As before, we seek a system of equations for the rates of change of the molecular concentrations, but distinguish between whether we assume that transport is fast enough to remove from the model or not. We additionally augment each equation system with the double receiver module.

**Slow transport model**

We start by expanding the derivatives on the left-hand sides, then defining $\gamma := \dfrac{1}{V_i}\dfrac{dV_i}{dt}$ as the specific growth rate, or *dilution* factor. Therefore, we obtain

$$\frac{d[A]}{dt} = a_A.f_{\text{Pbad}}(\text{Ara}) - (d_A + \gamma).[A] \tag{S29a}$$

$$\frac{d[C_6]_i}{dt} = \frac{1}{V_i}.\eta_6\left([C_6]_e - [C_6]_i\right) - d_{A6}.\frac{[A][C_6]_i}{1 + K_{A6}[C_6]_i + K_{A12}[C_{12}]_i} - (d_C + \gamma).[C_j] \tag{S29b}$$

$$\frac{d[C_{12}]_i}{dt} = \frac{1}{V_i}.\eta_{12}\left([C_{12}]_e - [C_{12}]_i\right) - d_{A12}.\frac{[A][C_{12}]_i}{1 + K_{A6}[C_6]_i + K_{A12}[C_{12}]_i} - (d_C + \gamma).[C_j] \tag{S29c}$$

$$\frac{d[C_6]_e}{dt} = \frac{1}{V_e}.\eta_6\left([C_6]_i - [C_6]_e\right) - \frac{1}{V_e}.\frac{dV_e}{dt} \tag{S29d}$$

$$\frac{d[C_{12}]_e}{dt} = \frac{1}{V_e}.\eta_{12}\left([C_{12}]_i - [C_6]_e\right) - \frac{1}{V_e}.\frac{dV_e}{dt} \tag{S29e}$$

Two assumptions could be applied to $V_e$: (1) $V_e$ is constant, or (2) $V_e = V_{\text{tot}} - V_i$, where $V_{\text{tot}}$ is a fixed total volume.

**Fast transport model**

If we assume that HSL transport is infinitely fast, then this results in the relations $[C_k]_e = [C_k]_i = [C_k]$ for $k = 6, 12$. However, we must be careful to maintain conservation of mass with respect to AiiA-mediated degradation. That is, we must scale the stoichiometric loss of HSL by the ratio of the extracellular and intracellular volumes. To see this, we write out equations for the new variables $[C_k]$ in terms of its constituent compartments.

$$\frac{d(V_i.[A])}{dt} = V_i.a_A.f_{\text{Pbad}}(\text{Ara}) - V_i.d_A.[A] \tag{S30a}$$

$$\frac{d(V.[C_6])}{dt} = \frac{d(V_i.[C_6]_i)}{dt} + \frac{d(V_e.[C_6]_e)}{dt} = -V_i.d_{A6}.\frac{[A][C_6]}{1 + K_{A6}[C_6] + K_{A12}[C_{12}]} - V_i.d_C.[C_6] \tag{S30b}$$

$$\frac{d(V.[C_{12}])}{dt} = \frac{d(V_i.[C_{12}]_i)}{dt} + \frac{d(V_e.[C_{12}]_e)}{dt} = -V_i.d_{A12}.\frac{[A][C_{12}]}{1 + K_{A6}[C_6] + K_{A12}[C_{12}]} - V_i.d_C.[C_{12}] \tag{S30c}$$

By dividing out the volumes from the derivatives on the left-hand sides, and assuming that $V$ (the total volume) is constant and $d_C$ is negligible, we obtain

$$\frac{d[A]}{dt} = a_A.f_{\text{Pbad}}(\text{Ara}) - (d_A + \gamma).[A] \tag{S31a}$$

$$\frac{d[C_6]}{dt} = -d_{A6}.c.\frac{[A][C_6]}{1 + K_{A6}[C_6] + K_{A12}[C_{12}]} - \tag{S31b}$$

$$\frac{d[C_{12}]}{dt} = -d_{A12}.c.\frac{[A][C_{12}]}{1 + K_{A6}[C_6] + K_{A12}[C_{12}]} \tag{S31c}$$

where $\gamma$ is the dilution factor as before and $c = \frac{V_i}{V}$ is the cell density from above.

As AiiA concentrations are never observed, we can arbitrarily rescale $[A]$, reducing the parametric complexity of the model. By substituting $[A] = k_A.a$, we obtain

$$\frac{da}{dt} = f_{\text{Pbad}}([C_6], [C_{12}]) - (d_A + \gamma)a \tag{S32a}$$

$$\frac{d[C_6]}{dt} = -c.\hat{d}_6.\frac{a.[C_6]}{1 + K_{A6}[C_6] + K_{A12}[C_{12}]} \tag{S32b}$$

$$\frac{d[C_{12}]}{dt} = -c.\hat{d}_{12}.\frac{a.[C_{12}]}{1 + K_{A6}[C_6] + K_{A12}[C_{12}]} \tag{S32c}$$

Finally, we note that the equations above can be trivially concatenated with the equations of the double receiver device described in the previous section to produce a full model of each relay device.

##### Characterization experiment

We measured the Degrader circuit in response to varying concentrations of the arabinose inducer. We used (uninformative) uniform priors on the previously uncharacterized parameters, and (informative) truncated Gaussian priors on parameters that had been previously characterized with mean and standard deviation taken from the marginal posteriors of the Receivers and Inducer circuit characterizations. We also use uninformative priors on the $B_k^{\text{back}}$ parameters, without propagating their marginal posteriors from previous circuits.

##### Implementation with GEC/CRN language

We define a module for the AiiA enzyme, which includes the modelled enzymatic degradation of C6 and C12. As before, we use a rate expression argument to arbitrarily assign a promoter transfer function to the production rates of each enzyme. However, since AiiA concentrations are never observed, the maximal production rate can be *divided out*, as explained above, and without loss of generality set to a value 1.

```
module AiiA(P,aI,growth,capacity) = {
    ->[[capacity]*aI*[P]] aiiA |
    aiiA ->{daiiA} |
    aiiA ->[[growth]*[aiiA]] |
    c6 -> [[x]*dA6*[c6]*[aiiA]] |
    c12 -> [[x]*dA12*[c12]*[aiiA]]
}
```

The GEC implementation of the Degrader circuit becomes

```
device drR33S175() = { LuxR(aR33,growth,capacity) | LasR(aS175,growth,capacity) |
     YFP(P81,aYFP,growth,capacity) | CFP(P76,aCFP,growth,capacity) }
device pBadAiiA() = { AiiA(PBAD,1.0,growth,capacity) }

system aiia = {
    directive simulation { final=36.0; points=250; plots=[[x]*([yfp]+[f530])+b530;
         [x]*([cfp]+[f480])+b480]; plotcolours=["#FFDF00"; "#ADD8E6"] }
    directive parameters [
        r = 1, { interval=Real; distribution=Uniform(0.1,10); variation=Multiple };
        K = 2, { interval=Real; distribution=Uniform(0.1,5); variation=Multiple };
        tlag = 1, { interval=Real; distribution=Uniform(0,10); variation=Multiple };
        rc=1e2,  { interval=Log; distribution=Uniform(1e0,1e5); variation=Multiple };
        b530=1e3, { interval=Real; distribution=Uniform(0.0,5e3); variation=Random };
        b480=1e3, { interval=Real; distribution=Uniform(0.0,1e4); variation=Random };
    ]
    directive rates [growth = [grow]*r*(1 - [x] / K);capacity = rc]
    directive deterministic {reltolerance=1e-12; abstolerance=1e-10}
    directive crn { cells(growth,tlag) | autofluorescence(growth,capacity) }
    | drR33S175:device
    | pBADAiiA:device
}
```

##### Characterization results

We show results for direct and ratiometric dynamic characterization for the AiiA circuit under the TargetSwitch hypothesis.

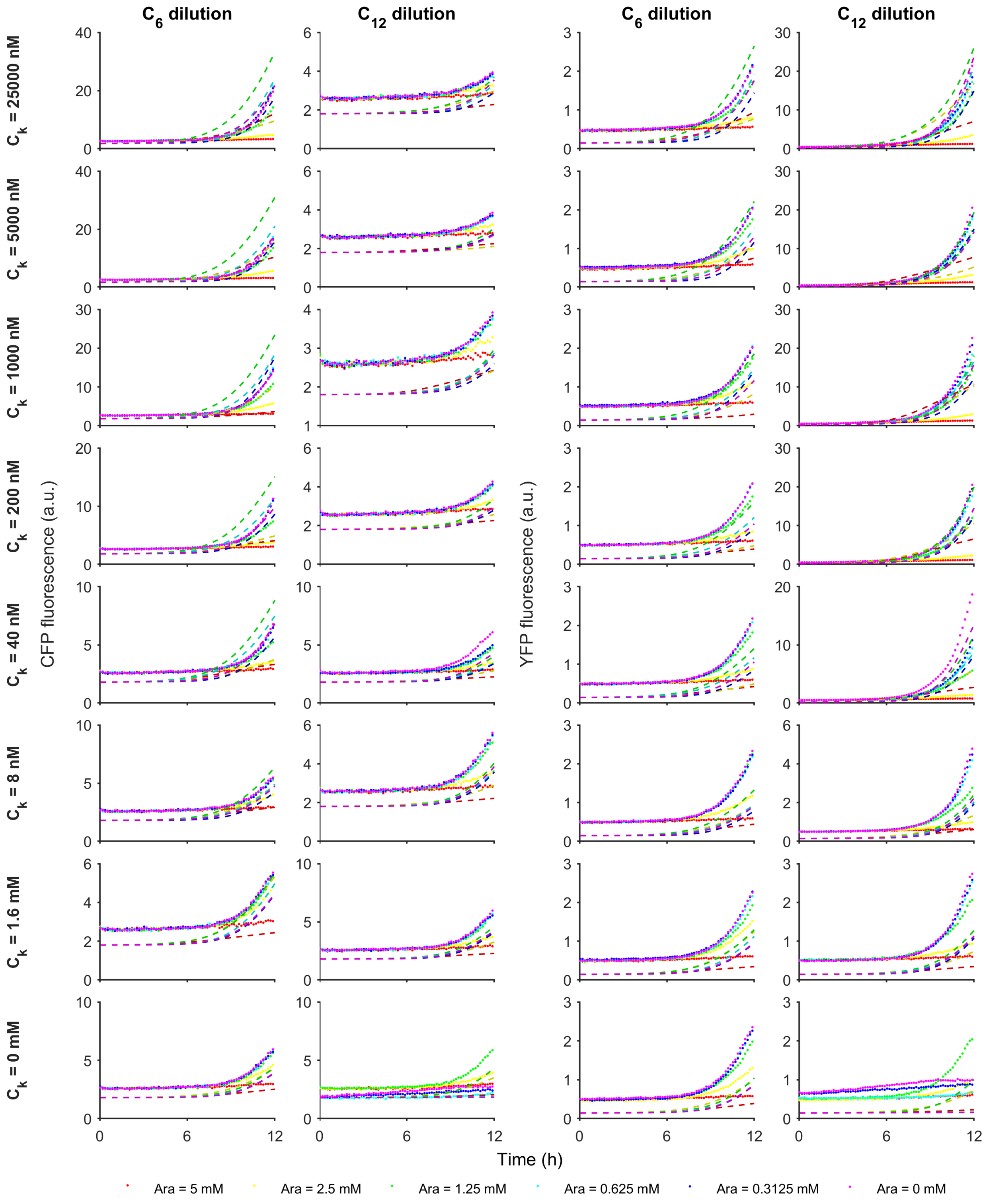

**Figure S27: Model-data comparison for the Degrader circuit with the TargetSwitch hypothesis: direct method.** Measurements of the TargetSwitch circuit are compared against simulations of the maximum likelihood parameter set from the TargetSwitch hypothesis.

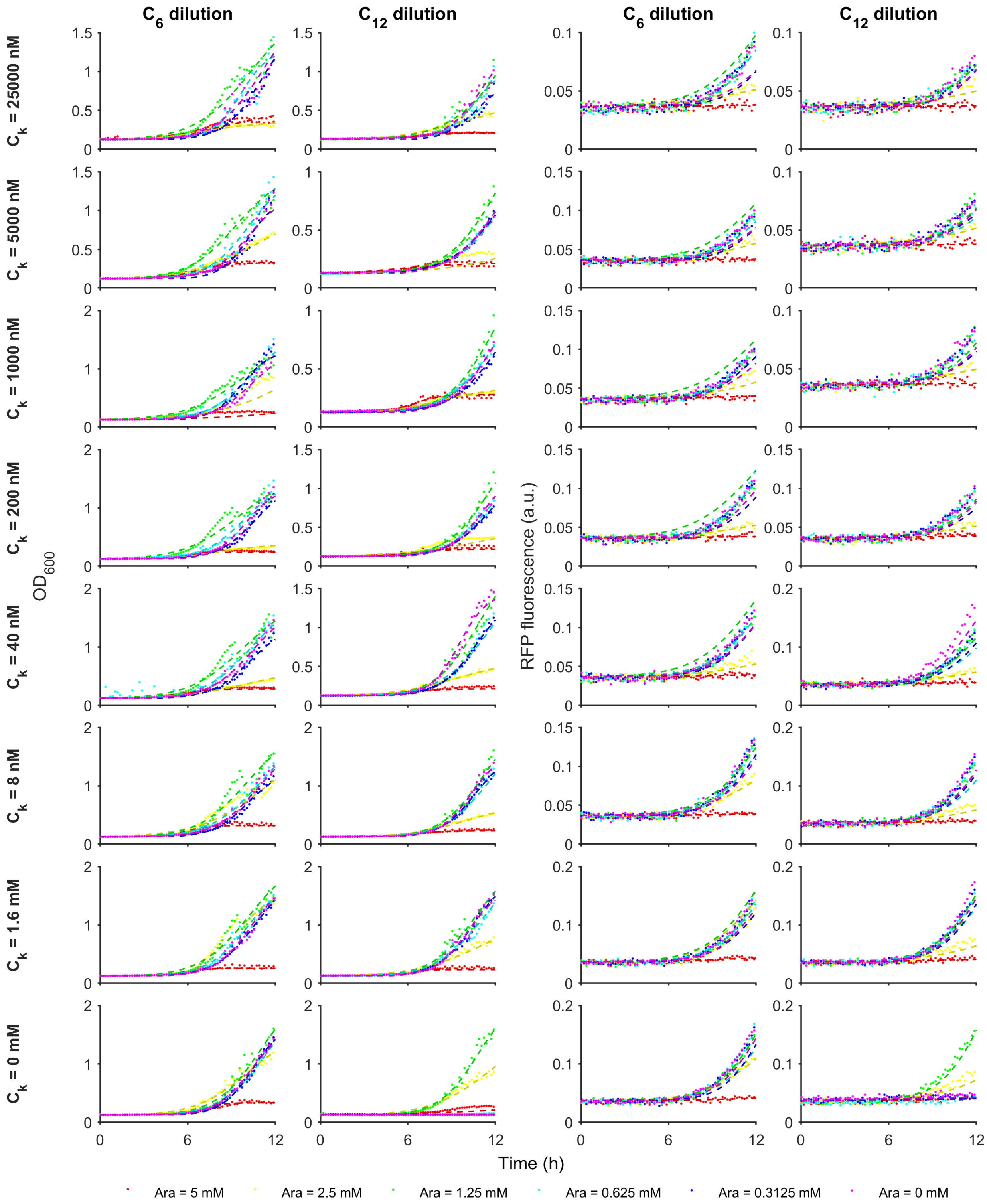

**Figure S28: Model-data comparison for the Degrader circuit with the TargetSwitch hypothesis: OD and control phase.** Measurements of the Degrader circuit are compared against simulations of the maximum likelihood parameter set from the TargetSwitch hypothesis.

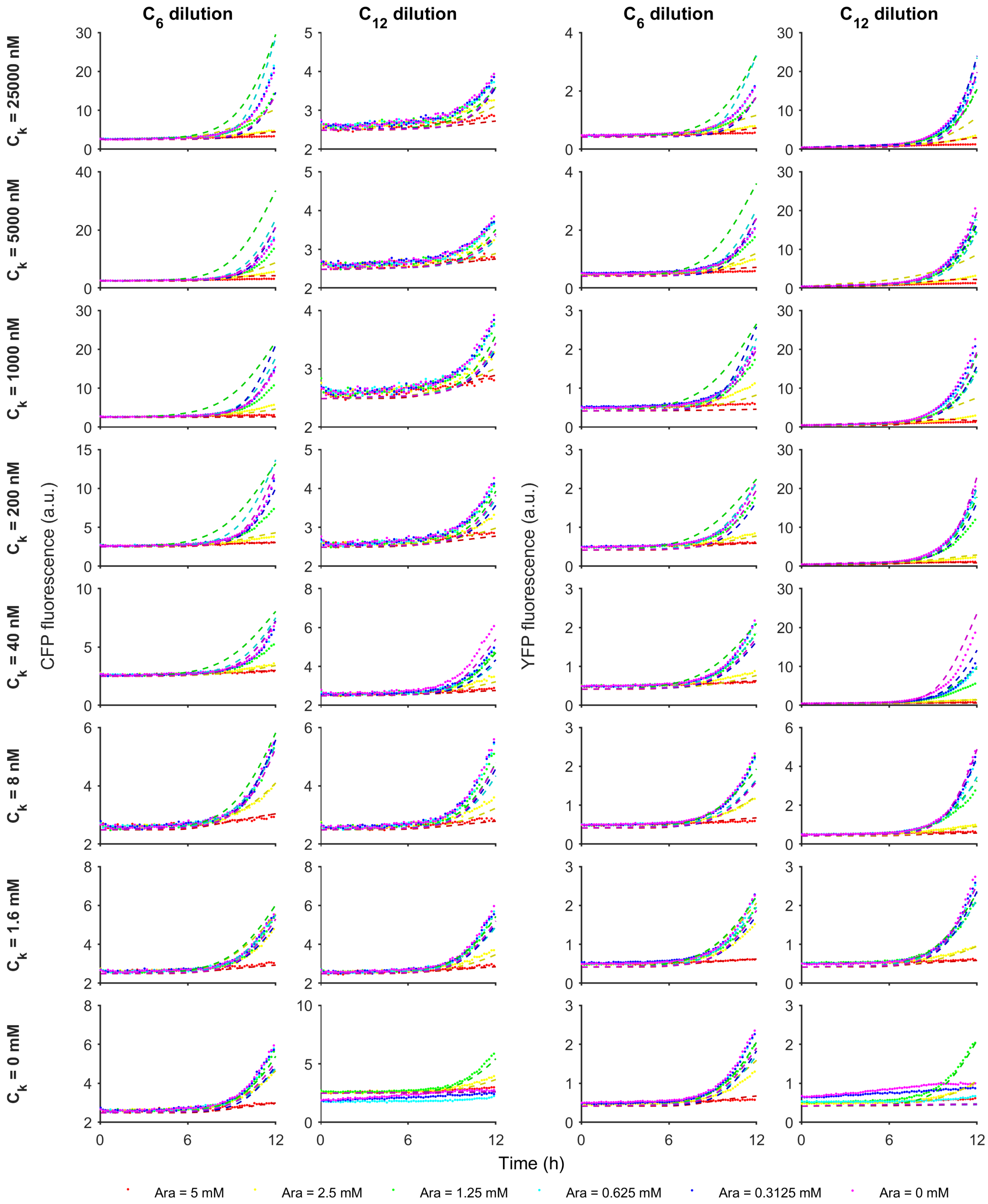

**Figure S29: Model-data comparison for the Degrader circuit with the TargetSwitch hypothesis: ratiometric method.** Measurements of the Degrader circuit are compared against simulations of the maximum likelihood parameter set from the TargetSwitch hypothesis.

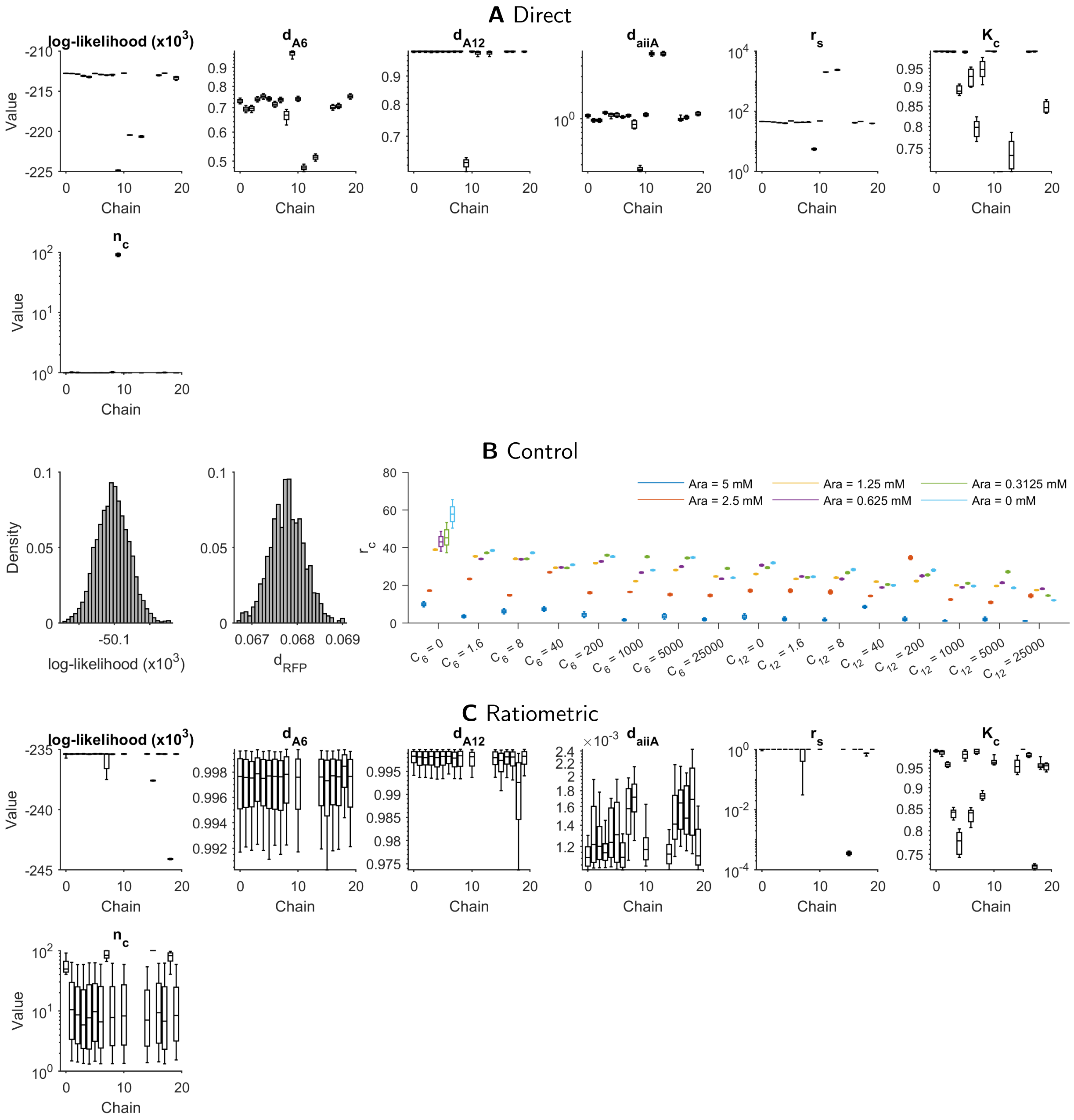

**Figure S30: Marginal parameter posterior distributions for the Degrader circuit with TargetSwitch hypothesis. A.** Inference of circuit parameters using the direct (non-ratiometric) method, which corresponds to assuming that $r_c = 1$ for all time-series traces. **B.** The control phase identifies values for $r_c$ for each time-series, to be used in ratiometric characterization. **C.** Inference of circuit parameters using the ratiometric method, using the values of $r_c$ in B.

### S3 Supplementary Figures

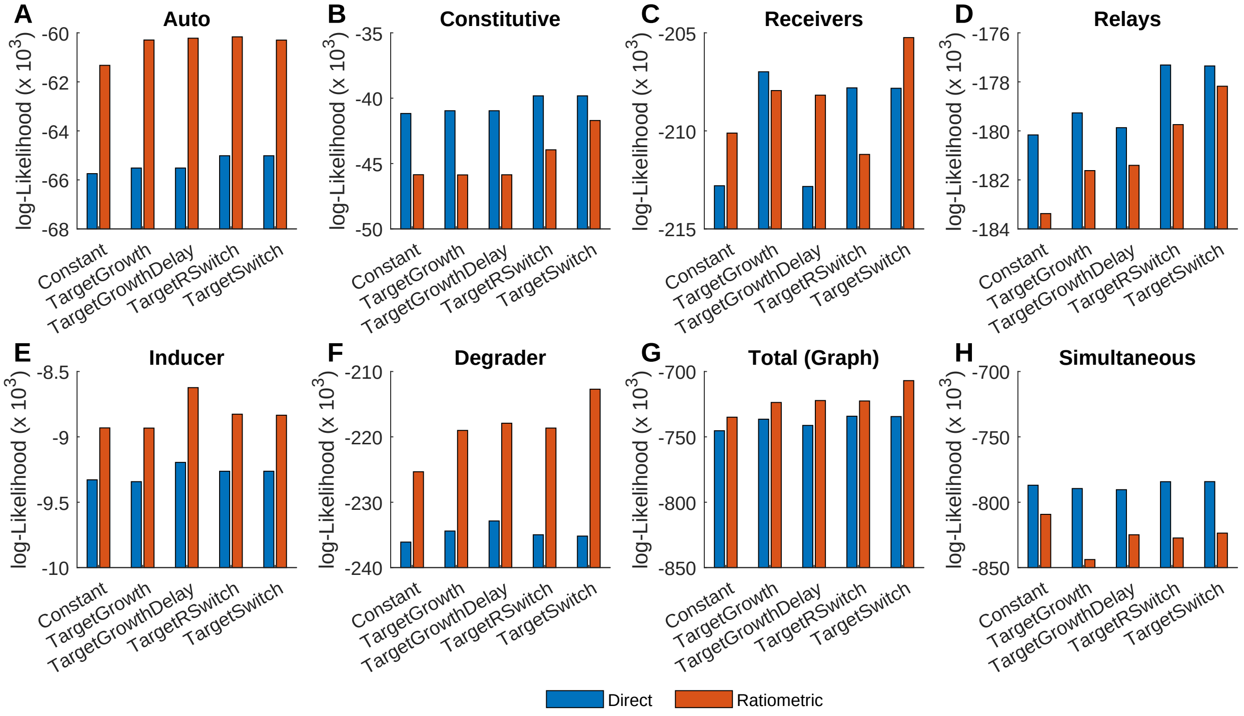

**Figure S31: Maximum-likelihood performance of dynamic characterization.** Direct and ratiometric dynamic characterization was applied sequentially to synthetic gene circuits implementing (**A**) no expression (Auto), (**B**) constitutive expression (PRPR), (**C**) induced expression of HSL signals (DR), (**D**) relayed induced expression using HSLs (Relays), (**E**) arabinose-inducible expression (Inducer ) and (**F**) disrupted HSL-inducible expression via arabinose-induced HSL degradation (AiiA). (**G**) Aggregated maximum likelihood score across A–F. (**H**) Maximum likelihood score of dynamic characterization applied simultaneously to the aforementioned circuits. In all cases, 5 different hypotheses about the dynamics of the gene expression capacity function were explored (see Table 1 for details).

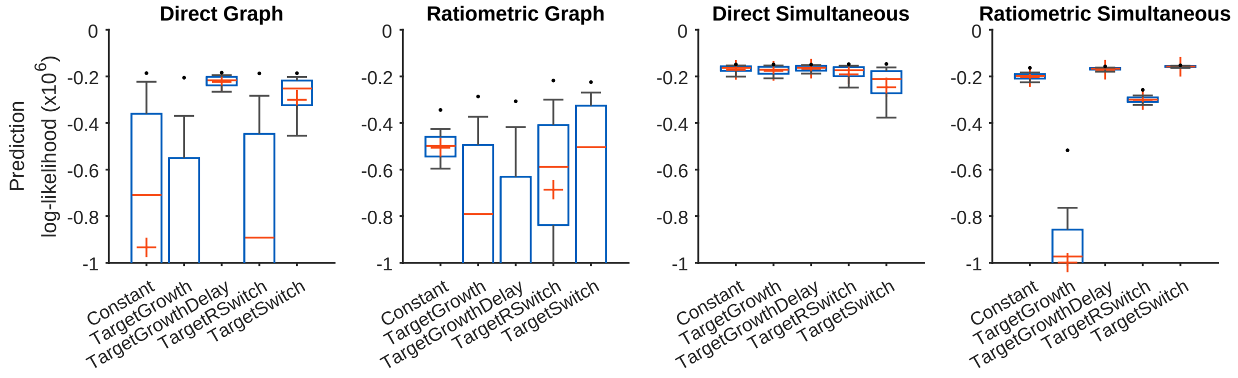

**Figure S32: Predictive performance of different gene expression capacity hypotheses.** The posterior predictive distribution against the data in Fig. 4C, and is quantified for ratiometric and direct dynamic characterization, both using the graph-based and simultaneous methods. Five hypotheses are compared for how gene expression capacity changes with culture growth. The box-plots indicate the interquartile range of samples from the marginalized parameter posteriors. The red plus symbols indicate the means of each distribution.

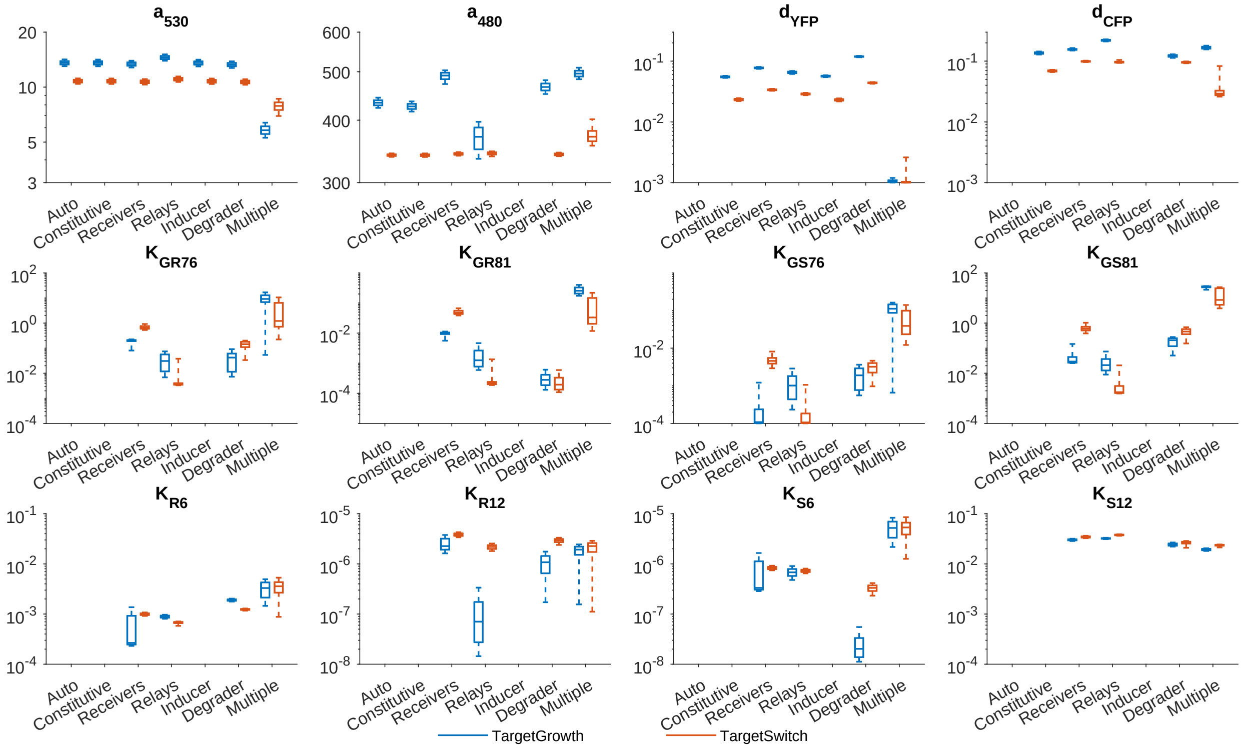

**Figure S33: Evolution of marginal parameter distributions using direct dynamic characterization.**

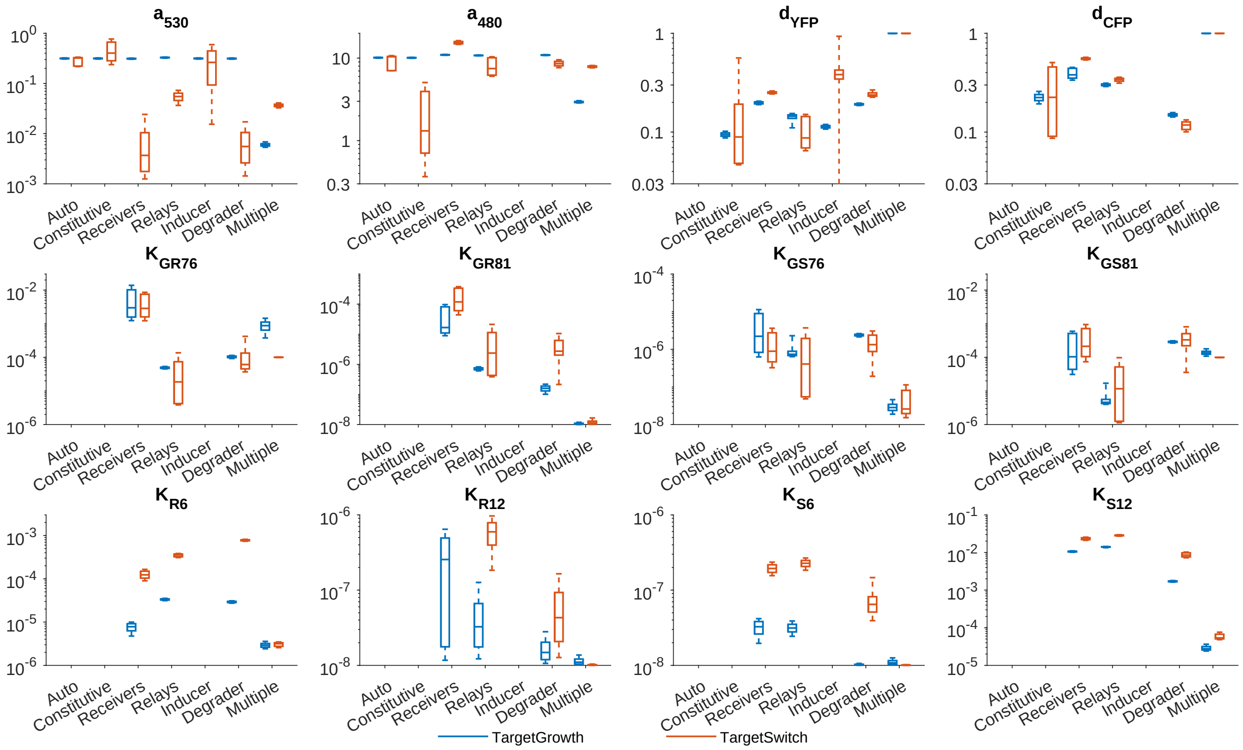

**Figure S34: Evolution of marginal parameter distributions using ratiometric dynamic characterization.**

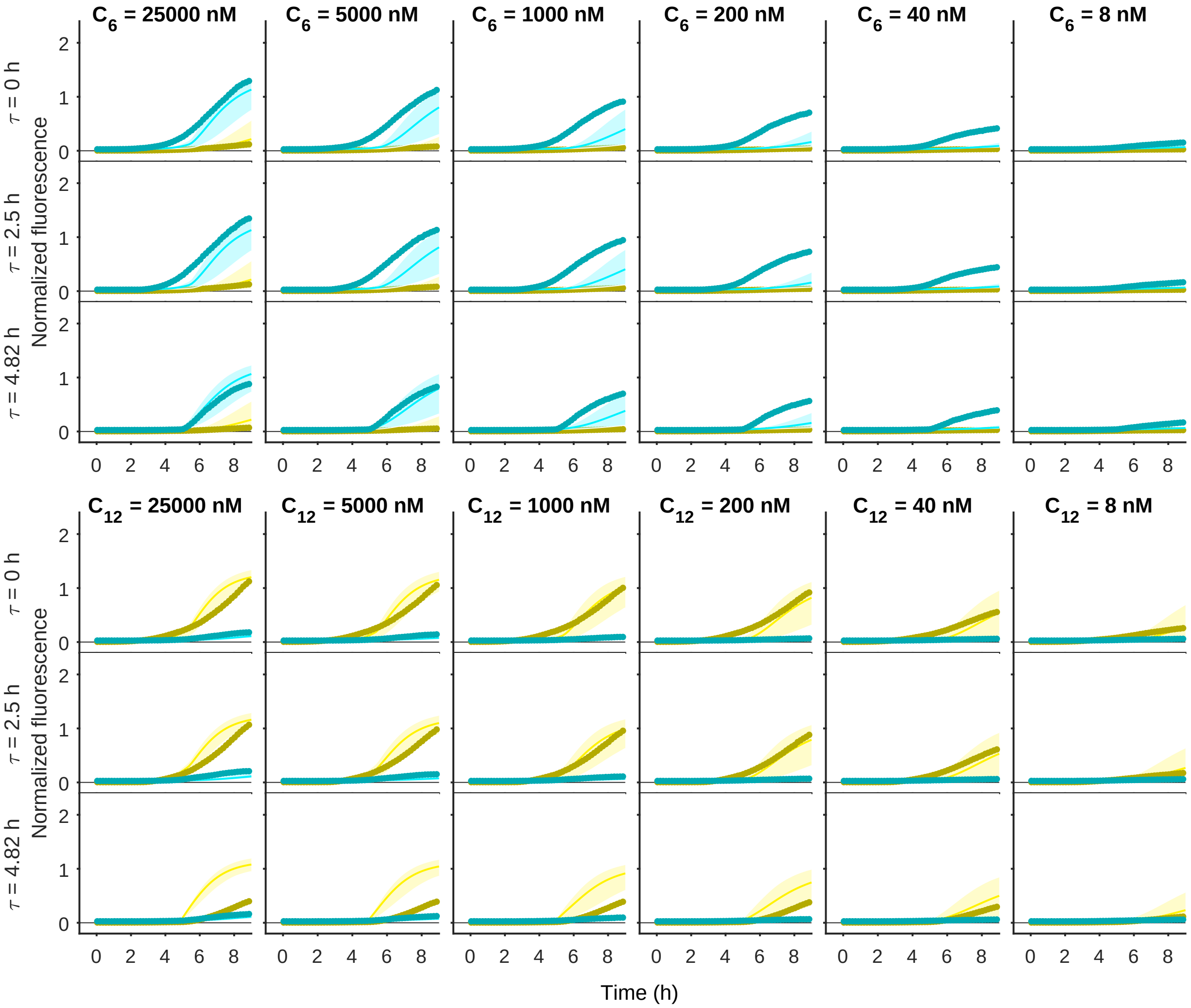

**Figure S35: Prediction of unseen data.** Comparison of the posterior predictive distribution for the TargetSwitch model using the ratiometric simultaneous method. In each column, the HSL treatment is indicated at the top, and in each row the time of HSL addition is noted on the left. eYFP and eCFP data are shown as thick darker lines (yellow and cyan respectively), with model simulations depicted as the mean (thin solid lines) and 95% credibility intervals as the shaded region.

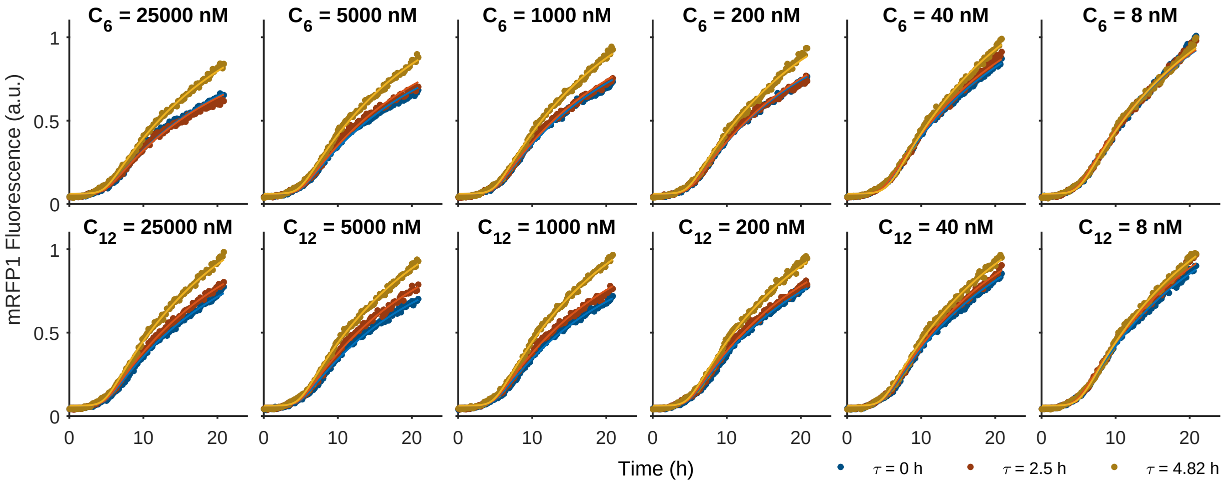

**Figure S36: RFP expression in R33-S175 following delayed HSL treatment.**

### S4 Supplementary Tables

| Plasmid name | Promoter | RBS | CDS | Origin/antibiotic |
|---|---|---|---|---|
| PRPR | PR | B0034 | eCFP; eYFP | P15/aKan |
| PBAD-YFP | PBAD | B0034 | eYFP | PSC101/Chlor |
| PBAD-AiiA | PBAD | B0034 | AiiA | PSC101/Chlor |

**Table S1:** Plasmids not previously described.

| Part Name | Part Type | BioBrick Id |
|---|---|---|
| pLas81 | Promoter | - |
| pLux76 | Promoter | - |
| pBAD | Promoter | BBa_I0500 |
| pCat | Promoter | BBa_I4033 |
| pLac | Promoter | BBa_R0011 |
| pTet | Promoter | BBa_R0040 |
| pr | Promoter | BBa_R0051 |
| rbs900 | RBS | - |
| rbsS100 | RBS | - |
| rbsS175 | RBS | - |
| rbs32 | RBS | BBa_B0032 |
| rbs33 | RBS | BBa_B0033 |
| rbs34 | RBS | BBa_B0034 |
| mrfp1 | PCR | - |
| aiia | PCR | BBa_C0060 |
| luxI | PCR | BBa_C0161 |
| lasI | PCR | BBa_C0178 |
| ecfp | PCR | BBa_E0420 |
| eyfp | PCR | BBa_E0430 |
| luxR | PCR | BBa_J69010 |
| lasR | PCR | BBa_J69121 |
| l3s2p21 | Terminator | - |
| ter1 | Terminator | BBa_B0015 |

**Table S2:** Parts Database

| Device Name | Components |
|---|---|
| auto | pr; rbs34; mrfp1; ter1 |
| prpr | pr; rbs34; eyfp; ter1;<br>pr; rbs34; ecfp; ter1 |
| drPCat | pCat; rbs34; luxR; rbs34; lasR; ter1;<br>pLas81; rbs34; eyfp; ter1;<br>plux76; rbs34; ecfp; ter1 |
| drR100S32 | pTet; rbss100; luxR; ter1;<br>pLac; rbs32; lasR; ter1;<br>pLas81; rbs34; eyfp; ter1;<br>plux76; rbs34; ecfp; ter1 |
| drR33S32 | pTet; rbs33; luxR; ter1;<br>pLac; rbs32; lasR; ter1;<br>pLas81; rbs34; eyfp; ter1;<br>plux76; rbs34; ecfp; ter1 |
| drR33S175 | pTet; rbs33; luxR; ter1;<br>pLac; rbsS175; lasR; ter1;<br>pLas81; rbs34; eyfp; ter1;<br>plux76; rbs34; ecfp; ter1 |
| relayP76LasI | pLux76; rbs900; lasI; l3s2p21 |
| relayP81LuxI | pLas81; rbs32; luxI; l3s2p21 |
| pBADYFP | pBAD; rbs34; eyfp; l3s2p21 |
| pBADAiiA | pBAD; rbs34; aiia; l3s2p21 |

**Table S3:** Device Database
